# Inhibitor-induced supercharging of kinase turnover via endogenous proteolytic circuits

**DOI:** 10.1101/2024.07.10.602881

**Authors:** Natalie S. Scholes, Martino Bertoni, Arnau Comajuncosa-Creus, Katharina Kladnik, Fabian Frommelt, Matthias Hinterndorfer, Hlib Razumkov, Polina Prokofeva, Martin P. Schwalm, Hana Imrichova, Eleonora Barone, Caroline Schätz, Andrea Rukavina, Anna Koren, Stefan Kubicek, Stefan Knapp, Nathanael S. Gray, Giulio Superti-Furga, Bernhard Kuster, Patrick Aloy, Georg E. Winter

**Affiliations:** CeMM, Research Center for Molecular Medicine of the Austrian Academy of Sciences, Vienna, Austria; Institute for Research in Biomedicine (IRB Barcelona), Barcelona, Catalonia, Spain; Department of Chemistry, Stanford School of Humanities and Sciences, Stanford University, CA, USA; Department of Chemical and Systems Biology, ChEM-H, Stanford School of Medicine, Stanford University, Stanford, CA, USA; Chair of Proteomics and Bioanalytics, Technical University of Munich, Freising, Germany; Institute of Pharmaceutical Chemistry, Goethe University, Frankfurt am Main, Germany; Stanford Cancer Institute, Stanford School of Humanities and Sciences, Stanford University, CA, USA; Center for Physiology and Pharmacology, Medical University of Vienna, Vienna, Austria; German Cancer Consortium (DKTK), partner site Munich and German Cancer Research Center (DKFZ), Heidelberg, Germany; Bavarian Biomolecular Mass Spectrometry Center (BayBioMS), Technical University of Munich, Freising, Germany; Institució Catalana de Recerca I Estudis Avançats (ICREA), Barcelona, Catalonia, Spain

## Abstract

Targeted protein degradation has emerged as a promising new pharmacological strategy. Traditionally, it relies on small molecules that induce proximity between a target protein and an E3 ubiquitin ligase to prompt target ubiquitination and degradation by the proteasome. Sporadic reports indicated that ligands designed to inhibit a target can also induce its destabilization. Among others, this has repeatedly been observed for kinase inhibitors. However, we lack an understanding of the frequency, generalizability, and mechanistic underpinnings of these phenomena. To address this knowledge gap, we generated dynamic abundance profiles of 98 kinases after cellular perturbations with 1570 kinase inhibitors, revealing 160 selective instances of inhibitor-induced kinase destabilization. Kinases prone to degradation are frequently annotated as HSP90 clients, thus affirming chaperone deprivation as an important route of destabilization. However, detailed investigation of inhibitor-induced degradation of LYN, BLK and RIPK2 revealed a differentiated, common mechanistic logic where inhibitors function by inducing a kinase state that is more efficiently cleared by endogenous degradation mechanisms. Mechanistically, effects can manifest by ligand-induced changes in cellular activity, localization, or multimerization which may be triggered by direct target engagement or network effects. Collectively, our data suggest that inhibitor-induced kinase degradation is a common event and positions supercharging of endogenous degradation circuits as an alternative to classical proximity-inducing degraders.

## Introduction

Targeted protein degradation is a pharmacological strategy that promises to overcome limitations of classical inhibitor-based approaches. Small-molecule degraders typically function by inducing proximity between a target protein and an E3 ubiquitin ligase to induce ubiquitination and proteasomal degradation of the target. Heterobifunctional PROTACs induce proximity by simultaneously engaging target and E3 with two dedicated, connected ligands. In contrast, molecular glue degraders induce proximity by binding either the target or the E3. Thereby, they chemically adapt its protein surface to prompt a highly cooperative target:ligand:E3 complex.

In addition, sporadic accounts of inhibitor-induced target degradation have been reported, for instance for inhibitors of gene-regulatory proteins such as BCL6^1,2^ or EZH2^3^ and, most frequently, for kinase inhibitors^4–6^. Many kinases require chaperones such as HSP90 for folding and maintained stability^7^. Upon kinase binding, some inhibitors, including compounds in clinical evaluation or practice, have been shown to disrupt kinase:HSP90 interactions, which in turn results in kinase destabilization^8^. This process of chaperone deprivation is well established and exemplified by the degradation of HER2 by neratinib^9^, or the degradation of LMTK3 by C28^10^. Given that hyperactive mutant kinases are frequently more reliant on HSP90, chaperone deprivation has also been associated with preferential degradation of mutant over *wildtype* kinases, such as the destabilization of EGFR^G719S^ by erlotinib^11^. However, detailed studies, for example of the mechanism of mutant selective PI3Kα degradation by taselisib and inavolisib^12,13^, revealed that mechanisms can go beyond the widely accepted framework of chaperone deprivation.

Due to the high conservation of the ATP binding site in the human kinome, many orthosteric inhibitors bind unselectively to diverse kinases. However, promiscuous inhibitors can act as rather selective destabilizers, as exemplified by sorafenib, which binds to at least 46 kinases^14^ but destabilizes only BRAF^V600E^ or RET^M918T^ ^11,15^. Importantly, this polypharmacology brings about an opportunity of “network drugging” where an inhibitor induces kinase degradation without engaging the kinase directly. Mechanistically, this can for instance occur via modulation of an upstream kinase that activates a phosphodegron, leading to the degradation of a downstream kinase, as described for WEE1^16^. Network effects can also be at play with directly acting inhibitors. For instance, mutant PI3Kα degradation by inavolisib is dependent on hyperactive HER2 signaling^13^.

Given that the discovery of monovalent kinase degraders has largely been episodic and serendipitous, we lack systematic insights into its pervasiveness and underlying fundamental mechanistic principles beyond chaperone deprivation. Due to the importance of kinases/kinase inhibitors in modern medicine (80 FDA approved drugs as of January 2024, with a further 180 in clinical trials^17^), efforts to quantify and mechanistically dissect inhibitor-induced kinase degradation could identify unanticipated therapeutic opportunities, explain adverse drug effects, or outline principles of degrader design beyond established proximity-inducing concepts.

To systematically identify inhibitor-induced kinase destabilization, we map the dynamic abundance profiles of 98 kinases after cellular perturbations with 1570 kinase inhibitors. Our efforts cover 88 canonical (*wildtype*) and 10 mutant kinases. In total we identify 232 compounds that downregulate protein levels of at least one kinase and 66 kinases that are affected by at least one compound. We discover that the predisposition of mutant kinases quantitatively and qualitatively differs from their *wildtype* counterparts. Even though frequently destabilized kinases are enriched for HSP90 clients, many of the observed degradation events cannot be explained by chaperone deprivation. Interestingly, we encounter that the propensity of a kinase to be destabilized by an inhibitor is not correlated with its degradability by PROTACs^18^. This suggests that inhibitor-induced degradation involves mechanisms that differ from traditional proximity-inducing modalities.

Detailed mechanistic follow-up of three previously undescribed kinase degraders suggests an underlying mechanistic principle in which inhibitors “supercharge” endogenous degradation circuits that preferentially recognize a particular kinase state. By inducing these states, inhibitors thus destabilize the cellular pool of a given kinase. We describe several mechanisms that lead to kinase degradation. For instance, inhibitors can induce degradation-prone kinase states by modulating kinase activity (exemplified by LYN), perturb intracellular kinase localization (BLK), and induce kinase multimerization (RIPK2). Notably, a parallel study revealed that inhibitors/degraders of IDO1, a heme enzyme that catalyzes the first step in tryptophan catabolism, function via a comparable mechanistic principle, stabilizing an apo-IDO1 state that is efficiently turned over by its endogenous E3 ligase KLHDC3 (Hennes et. al, 2024)^19^. Collectively, these findings highlight a unifying framework of inhibitor-induced target degradation that is prevalent for, but not limited to, kinases and is distinct from proximity-inducing modalities such as PROTACs or molecular glue degraders.

## Results

### Charting a monovalent degrader map via dynamic tracing of kinase abundance

To assess how inhibitors affect kinase levels, we opted for a scalable luminescent reporter setup using a lentiviral expression system where 98 kinase ORFs (88 canonical, 10 mutants) are expressed as Nanoluciferase (Nluc) fusions in K562 cells (**Fig 1A**, **Extended Data Table 1**, **Supplementary Data Table S1**). Mutants were selected based on disease-relevance or due to inducing drug resistance, as represented by a suite of EGFR mutants. Since this assay setup informs on the abundance rather than stability of a target protein, control measures were put into place, allowing us to segregate temporal inhibitor effects from global perturbations of transcription elongation or translation. Additionally, we profiled control cell lines expressing a long-lived non-kinase control target (GFP-Nluc) as well as a short-lived control (dGFP (destabilized GFP)-Nluc, see **SI Materials & Methods**). This panel of a total of 100 cell lines was dynamically assayed against an annotated library of 1570 kinase inhibitors at regular time intervals (2, 6, 10, 14 & 18 hours (h)). Compounds were selected for minimal assay interference, excluding Nluc quenchers and compounds that showed cytotoxicity during the assayed time window (**Supplementary Data Table S1**, see **SI Materials & Methods**). Scoring of the resulting extensive dataset was performed with a multi-tiered scheme using the following measures: exclusion of high variance duplicates or highly reactive compounds, scoring of individual normalized trajectories with respect to the 97 other kinases as well as against the internal degradation controls per kinase (see **Fig 1A**, SI **Materials and Methods**). This resulted in a total of 232 compounds that score and elicit destabilization across 66 of the tested kinases including 7 mutants (**Fig 1A, B**, **Supplementary Data Table S1**, **Extended Data Table 1**). Amongst all hits (see trajectories in **Supplementary Data Document 1**), we identified 160 unique kinase-compound pairs, which are denoted as selective (**Fig 1B**). Reporter half-life did not correlate (Spearman’s rank correlation = -0.381) with the frequency of scoring, supporting that the implemented controls successfully filtered out global perturbations (**Extended Data Fig 1A**).

**Fig 1.**
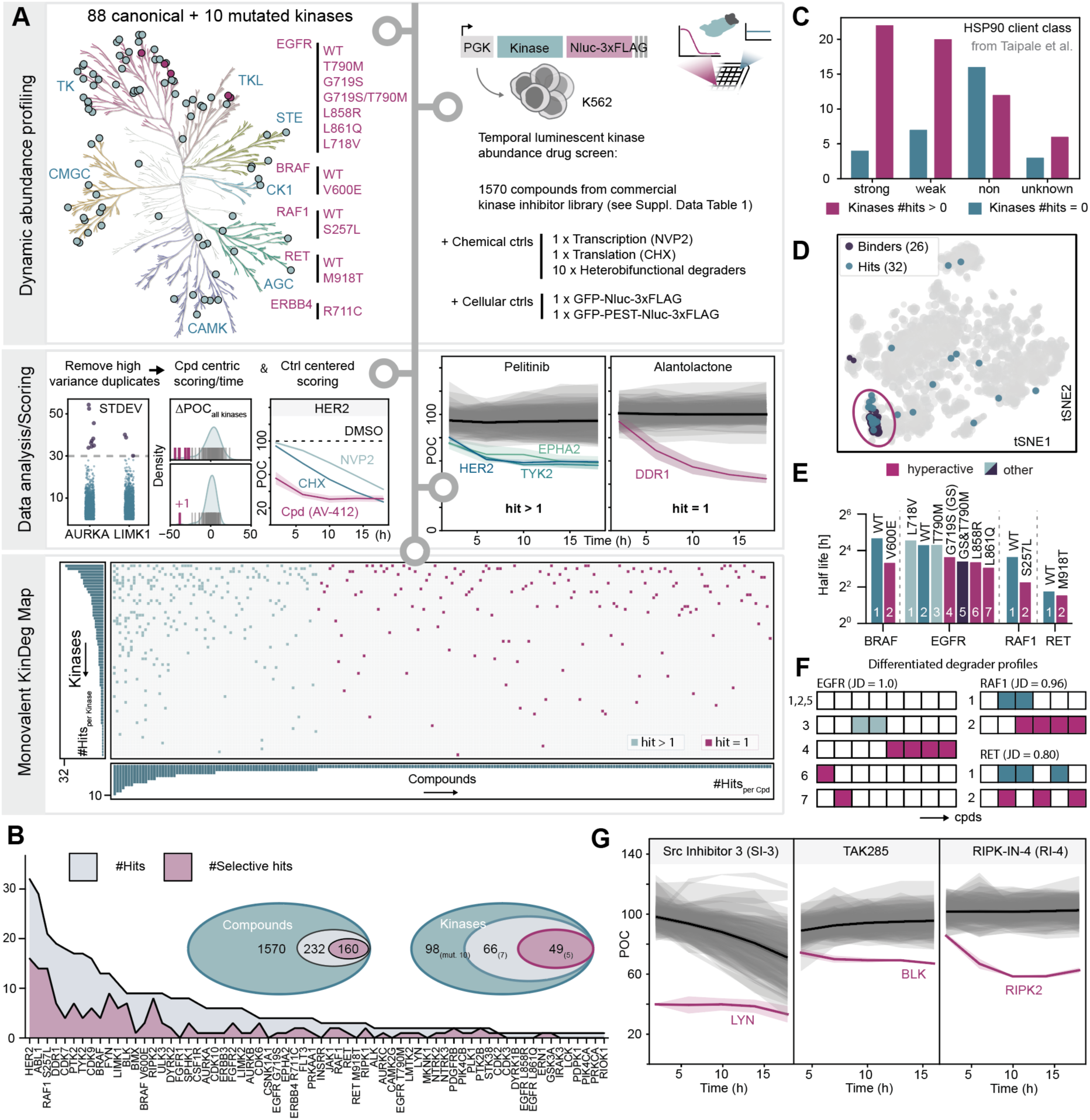
Kinase Degradation (KinDeg) map across 1570 monovalent kinase-targeting compounds. **A** Experimental and computational scheme to generate the KinDeg map. Top left: breakdown of analyzed kinases across the kinome tree, including 88 canonical kinases and 10 mutated kinases (highlighted in pink). Top right: schematic overview of drug screen setup including detailed breakdown of controls. Middle: Data processing and hit calling to classify if a compound perturbs a kinase’s abundance. Example temporal screening trajectories depicted for a small molecule with more than one downregulated kinase (hit > 1; Pelitinib) and an example for a selectively perturbed kinase (hit = 1; Alantolactone) (POC = percent over control). Bottom: resulting binary KinDeg map sorted according to the observed degradation frequencies across kinases and compounds including adjacent histograms of the summed scores in both dimensions. **B** Breakdown of hit scores across the kinases and compounds. **C** Stratified kinases for scoring at least once (downregulated) or non-scoring kinases with respect to their HSP90 client status as defined by Taipale *et al*^7^. **D** tSNE plot of compound target landscape (see **SI Materials and Methods**) across the utilized drug screening library with annotated HER2 binders (violet) and HER2 destabilizing screening hits (dark green) forming a co-cluster. **E** Comparisons of experimentally derived half-lives across mutant and canonical kinase pairs. **F** Representative hit comparison and Jaccard distance (JD) calculations across matched canonical and mutant EGFR, RAF and RET data. Number and color annotations follow the scheme depicted in (E). For the full dataset see **Supplementary Data Table 1**. **G** Temporal screening trajectories of the top three hits selected for detailed mechanism of action elucidation. All error bars are represented as confidence intervals (CI) of m = 2 (technical replicates).

Across all kinases, our top three examples based on total scores and selective hits were HER2, ABL1, and the mutant kinase RAF1^S257L^. No specific family was enriched among kinases prone to destabilization (**Extended Data Fig 1B**). When comparing our data to a recently published survey that assessed PROTAC-induced global kinase degradability^18^, no global correlation across the 41 shared kinases assessed in both studies was observed, suggesting that inhibitor-induced degradation mechanistically differs from degradation based on proximity induction (**Extended Data Fig 1C**). In contrast, when assessing the HSP90 status of each kinase^7^, both strong and weak clients had a markedly higher prevalence of being destabilized compared to non or not-defined clients, pointing to an outsized contribution of chaperone deprivation to the observed degradation events (**Fig 1C**). Among the annotated HSP90 clients was also the frequently destabilized HER2. Closer examination of the HER2-degrading compounds in our survey revealed that we successfully identified inhibitors, such as AV-412, afatinib or neratinib, which have previously been implicated as HER2 degraders that function via chaperone deprivation^9,20–22^. In addition, our data revealed new destabilizing inhibitors, such as WZ4002 or dacomitinib. Of note, many of the identified HER2 destabilizers feature covalent warheads. In line with a direct effect, introducing a C805S mutation^23^ in HER2 prevented inhibitor-induced degradation (**Extended Data Fig 1D, E**). Further supporting an on-target effect, the identified degraders (screening hits) formed a definite cluster when mapping them on their target space (**Fig 1D**)^24^.

With HER2 degraders serving as an example for directly acting degraders, we next set out to identify degradation events driven by network modulation. Mapping the experimentally identified ABL1 destabilizers on their target space, we observed a more dispersed distribution compared to HER2 degraders, yet we identified one cluster of hit compounds. In contrast to HER2 degraders, the cluster of ABL1 destabilizers did, however, not coincide with known ABL1 binders, but instead with dual PI3Kα and mTOR inhibitors (**Extended Data Fig 1F**). While we cannot exclude that the short half-life (1.5h) of ABL1 results in this prevalence of scoring, we note that the compounds failed to score in the control dGFP (half-life 1.1h) cell line (**Extended Data Fig 1G**, **SI Fig S2**), suggesting an effect beyond low baseline stability. Since the screen was conducted in the BCR-ABL driven leukemic cell line K562, many of the most potent ABL1 inhibitors were, however, eliminated from the screening library due to their acute cytotoxicity. This likely contributed to a relative underrepresentation of directly acting ABL1 degraders. Akin to HER2 and ABL1, we analyzed all remaining kinases across the available drug target space (**Extended Data** Fig 2). However, clustering of the hits on a per kinase basis failed to reveal generalizable trends, likely due to the limited number of identified degraders for most kinases and limited availability of comprehensive binding data for the assayed compounds.

Focusing our efforts on the assayed mutant kinases, we initially identified a decrease in protein half-life for activating mutations over non-activating mutations (**Fig 1E**), suggestive of an activity-stability tradeoff, in line with the observation that kinase activity is frequently counter-regulated by degradation^25^. Overall, we find distinctive degradation patterns with little overlap of hit compounds comparing *wildtype* and mutants (Jaccard distances (JD) ≥ 0.8, **Fig 1F**). While in some cases the degradation frequency corresponds to the reduced protein stability, we identify most hits for EGFR^G719S^, which is more stable than the strongly transformative mutants EGFR^L858R^ and EGFR^L861Q^. These observations imply that mechanistically, the enhanced degradability of mutant kinases goes beyond the reduced half-life and could be rooted in altered signaling circuits and protein interactomes.

After determining frequency and general features of inhibitor-induced degradation, we next set out to dissect the mechanism of action of three selective inhibitor-kinase pairs with different HSP90 client status^7^: LYN (strong client, degraded by Src inhibitor 3^26^; SI-3), BLK (weak client, degraded by TAK285^27^) and RIPK2 (non-client, degraded by RIPK-IN-4^28^; RI-4) (**Fig 1G**, **Extended Data Fig 1H**).

### Rapid destabilization of LYN kinase via Src inhibitor 3

One of the most striking effects recorded in our dataset was the degradation of the SRC family kinase LYN by the compound SI-3. Based on the measured abundance trajectories, effects of SI-3 on LYN levels were immediate with near-complete protein ablation after two hours (**Fig 1G**). Utilizing a previously described fluorescence protein stability reporter^29^, we first validated that SI-3 affects LYN stability (**Fig 2A**). The effect of SI-3 was also confirmed via immunoblot for endogenous LYN, revealing that degradation occurs in the nanomolar range (DC_50_ < 9 nM, **Fig 2B**) and already minutes after ligand exposure (t_1/2_ = 10.71 min, **Fig 2C**). Next, we sought to recapitulate the selectivity of SI-3-induced LYN degradation. Out of a total of 260 quantified kinases, including 2 of the 7 closely related SRC kinases and CSK, LYN was the most significantly destabilized kinase in a quantitative expression proteomics experiment (**Extended Data Fig 3A, B**). In line with the annotation as SRC family inhibitor and previously published kinome selectivity profiling^26^, we established that SI-3 inhibits recombinant LYN protein with an IC50 of 56.7 nM (**Extended Data Fig 3C**). Intriguingly, even though 25 of the 1570 profiled inhibitors are annotated to bind LYN, SI-3 was the only inhibitor that prompted LYN degradation (**Extended Data Fig 3D**).

**Fig 2.**
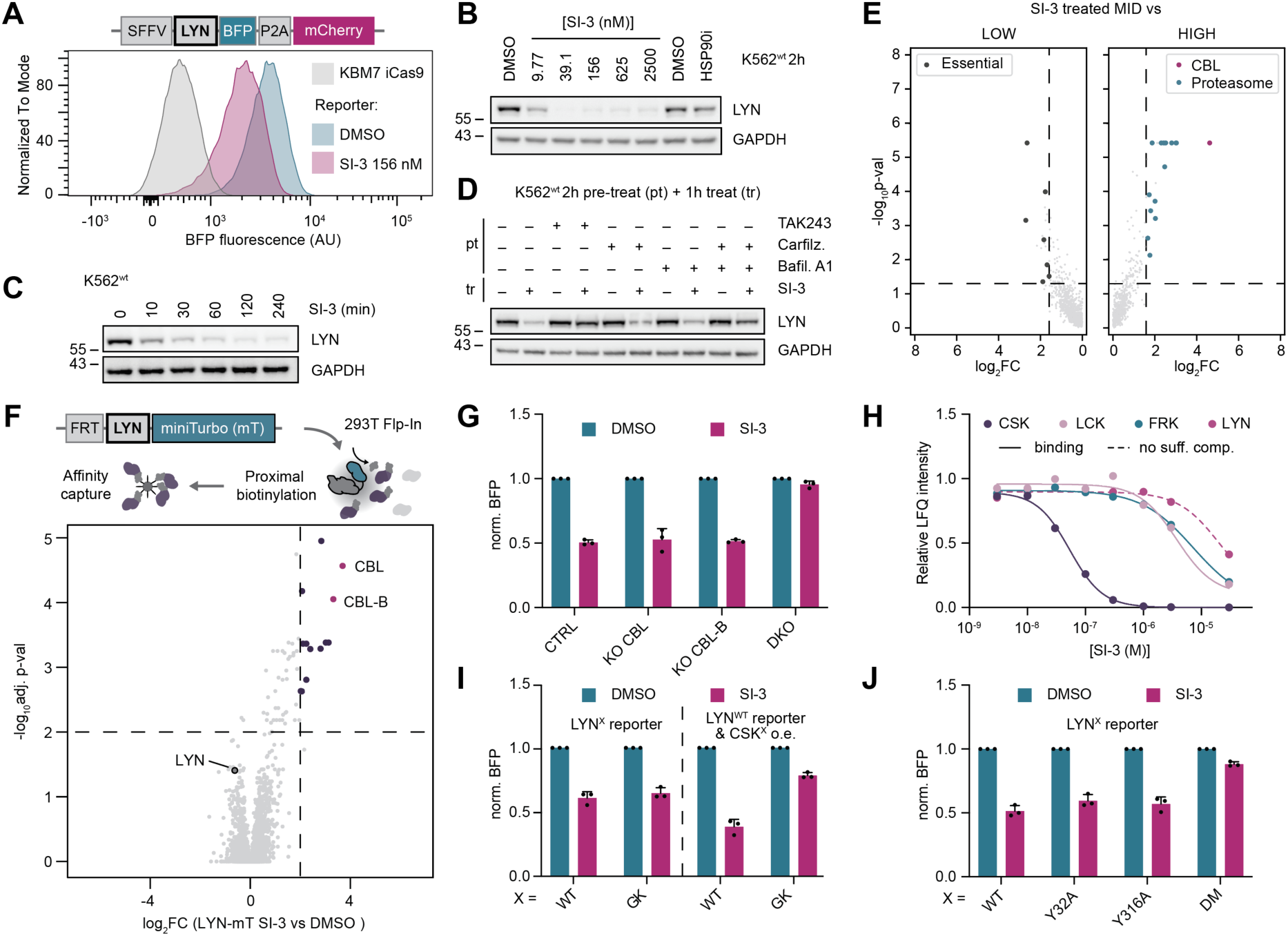
LYN is degraded rapidly by SI-3 via a canonical activity-stability switch. **A** Stability reporter setup for measuring SI-3-induced LYN degradation via flow cytometry in KBM7 iCas9 cells. **B** Dose-dependent degradation of endogenous LYN by SI-3 in K562 cells. DC_50_ < 9 nM, D_max_ = 95.55% (see quantification in **SI Fig S4B**) (n = 3). **C** Immunoblot of time-dependent LYN degradation on endogenous protein level via SI-3 (156 nM, K562 cells; see quantification in **SI Fig S4C**) (n = 3). **D** Immunoblot of chemical rescue phenotype of SI-3-induced degradation of LYN (n = 3). **E** Results of UPS-focused FACS-based CRISPR/Cas9 screen. Essential genes (Depmap 23Q4^36^) or CBL and proteasome subunits are highlighted for a p-value < 0.05 and a log_2_ fold-change (FC) of > 1.585 for the low-to-mid and high-to-mid comparisons, respectively (n = 2). **F** Differential BioID results after treatment of LYN-mT expressing 293T FlpIn cells with SI-3 or DMSO. Enriched prey is highlighted in dark lilac for a threshold of log_2_FC > 2 and -log_10_adj, p-value > 2 (adjusted p-value, BH) (n = 3). **G-J** SI-3 was utilized at 156 nM unless specified otherwise. Norm. (normalized) BFP is calculated as a raw ratio to mCherry and normalized with respect to the respective genetic perturbations or utilized constructs (see **SI Materials and Methods** for details). **G** Flow cytometric assay of LYN stability reporter upon genetic perturbation of either CTRL (sg*AAVS1*), sgRNAs targeting either *CBL* or *CBLB* (KO CBL or CBL-B), or an sgRNA targeting both E3s (dual KO, DKO) (n = 3). **H** Results of dose-ranging chemoproteomics (Kinobead profiling^37,38^) for SI-3 and selected hit kinases as well as LYN (dashed line, no sufficient competition). See **SI Fig S4G-I** for trajectories of all other hits. K_D_s: CSK = 34.60 nM, LCK = 2.987 µM, FRK = 11.43 µM. **J** Flow cytometric measurement of SI-3-mediated destabilization of the different phosphodegron dead reporter variants LYN^Y32A^, LYN^Y316A^ or LYN^Y32A+Y316A^ (double mutant, DM) compared to LYN^WT^ (n = 3). Flow cytometric assays were conducted in the cellular background of KBM7 iCas9 cells. All values represent the mean values ± SD; n = biological replicates, m = technical replicates.

While SI-3-induced LYN degradation was ubiquitin-dependent, it was not sensitive to inhibition of neddylation, thereby excluding dependency on a cullin RING E3 ubiquitin ligase (CRL) (**Fig 2D**). Moreover, degradation was not rescued by pharmacologic inhibition of the proteasome or lysosomal acidification. Only simultaneous inhibition of both degradation pathways prevented LYN degradation (**Fig 2D**). Albeit annotated as a strong HSP90 client^7^, pharmacological HSP90 inhibition did not induce LYN degradation both on endogenous and reporter setup, thus excluding chaperone deprivation as underlying mechanism (**Fig 2B**, **Extended Data Fig 3E**).

To decipher the mechanism of the SI-3-induced, potent and selective degradation of LYN, we opted for a two-pronged discovery approach geared to *(i)* identify which genes were required to induce LYN degradation following SI-3 treatment and *(ii)* chart how SI-3 treatment changed the interactome of LYN. To map genes required for SI-3-dependent LYN degradation, we performed a time-controlled, FACS-based CRISPR/Cas9 screen in KBM7 cells that feature an inducible Cas9 allele^29^ and the LYN stability reporter. This revealed the E3 ligase *CBL* as the strongest enriched gene (**Fig 2E**). CBL had previously been associated with the turnover of multiple SRC family kinases^30,31^, including LYN^32,33^. In support of a physiologically relevant interaction, we identified *CBL* as the strongest hit not only in SI-3 treated cells, but also in vehicle (DMSO)- treated conditions (**Extended Data Fig 3F**). To map how SI-3 altered LYN interactions, we opted for a BioID^34,35^ setup where LYN was expressed as fusion to the miniTurbo (mT) biotin ligase, allowing the identification of proteins that are recruited to LYN following cellular SI-3 treatment by mass spectrometry. In line with the CRISPR/Cas9 screen, this led to the identification of CBL as the strongest recruited effector (**Fig 2F**, **Extended Data Fig 3G**). Additionally, we found the closely related E3 ligase CBL-B as a strongly enriched LYN interactor in a SI-3-dependent manner, suggestive of a potential redundancy in the SI-3-induced LYN degradation. Indeed, single population-level knockout (KO) of CBL was insufficient to rescue SI-3-induced LYN degradation (**Fig 2G**), but drastically increased baseline levels (**Extended Data Fig 3H**) thus explaining why it scored as a hit in the CRISPR/Cas9 screen. Likewise, single KO of CBL-B showed an incomplete rescue. Only combined genetic disruption of both effectors fully rescued degradation (**Fig 2G**). Having identified this redundancy, we next addressed if the two E3s would separately be responsible for licensing LYN’s proteasomal or lysosomal degradation. Combining single ligase KO with pharmacological inhibition of either degradation pathway revealed that CBL-B-mediated degradation preferentially co-opts the proteasomal machinery, while degradation mediated by CBL relies on both proteasomal and lysosomal degradation (**Extended Data Fig 3I**).

Activation of LYN, for instance via IgE/FcχRI signaling, has been associated with the degradation of LYN via CBL/CBL-B^32^. Counterintuitively, this connects LYN activation to the degradation mechanism for the inhibitor SI-3, prompting us to closer investigate the cellular target spectrum of SI-3. Competitive, dose-ranging chemoproteomic profiling revealed 11 high confidence targets including 8 kinases (**Fig 2H**, **SI Fig S4G-I**). We identified CSK (K_D_ = 34.60 nM) as the most potently engaged kinase target, confirming previously reported recombinant assay data of SI-3 (IC_50_ = 4 nM CSK)^26^. Contrary to our *in vitro* data (**Extended Data Fig 3C**), LYN was only engaged at much higher concentrations (**Fig 2H**). Alongside LYN, the two SRC kinases LCK and FRK were additionally detected, however, again only at much higher concentrations. Orthogonally, we validated SI-3 target engagement with CSK using a NanoBRET displacement assay (EC50 = 15 nM, **Extended Data Fig 3J**). Based on the known role of CSK as a negative regulator of SRC kinases including LYN, we thus hypothesized that SI-3, despite inhibiting LYN, might induce indirect LYN degradation by its preferential inhibition of CSK. Similar observations have been made with a chemical genetics setup where engineered CSK could be perturbed by a mutant-selective inhibitor, leading to LYN activation and degradation^33^. To validate this hypothesis, we genetically modified either LYN or CSK to impair drug binding of SI-3 and assess consequences on SI-3-induced LYN degradation. Mutating the gatekeeper residue of LYN (T319I^39^) did not alter degradation capacity. However, as hypothesized, overexpression of the CSK gatekeeper mutant (T266M^40^) rescued degradation markedly (**Fig 2I**). This suggested inhibition of CSK as the dominant driver of SI-3 mediated LYN destabilization and that SI-3 functions by perturbing the intrinsic regulatory network. Consequently, blocking CSK-induced LYN activation by pharmacologic LYN inhibition rescued SI-3-induced LYN degradation on endogenous protein level (**Extended Data Fig 3K**) and phosphorylation of CSK’s target site LYN Y508 was reduced upon SI-3 treatment (**Extended Data Fig 3L, M**).

Prior work addressed the turnover of activated LYN, identifying Y32 as a phosphodegron important for CBL engagement^33^. However, LYN^Y32A^ was not resistant to SI-3 induced degradation, pointing to a possible additional, redundant phosphodegron (**Fig 2J**). Indeed, while degradation of LYN^wt^ was dependent on both CBL and CBL-B, we found that LYN^Y32A^ degradation solely depended on CBL (**Extended Data Fig 3N**), thus suggesting that this E3 recognizes the elusive phosphodegron. Turning back to our BioID dataset, we identified increased phosphorylation of Y316 in LYN after SI-3 treatment (**Extended Data Fig 3O**). Supporting a functional role of Y316, we found that the double mutant LYN^Y32A+Y316A^ was almost completely inert to SI-3-induced degradation (**Fig 2J**). The degradation of the single LYN^Y316A^ mutant was comparable to the degradation of *wildtype* LYN suggesting a redundancy of both phosphodegrons in SI-3-induced LYN degradation.

In summary, our data identified SI-3 as uniquely differentiated inhibitor that is sufficiently selective for CSK over LYN to exploit an endogenous activity-stability switch that ensures immediate and near-complete LYN degradation after its activation.

### Degradation by TAK285 depends on γ-secretase-induced, cytoplasmatic re-localization of BLK

From several inhibitors that cause downregulation of BLK in our assay, we focused on the selective hit TAK285. It destabilized BLK following a first order decay function without affecting abundance of any of the other assayed kinases (**Fig 1G**). First, we validated that TAK285 affects BLK stability, revealing that half-maximal degradation (DC_50_) occurs at 1 µM TAK285 and after around two hours of treatment (**Fig 3A, B**). TAK285-induced BLK degradation was ubiquitin- and proteasome-dependent, but neddylation independent and it was also independent of lysosomal degradation (**Fig 3C**, **Extended Data Fig 4A**). BLK was previously identified as a weak HSP90 client^7^ and showed sensitivity towards chaperone deprivation induced indirectly via HSP90 inhibition. HSP90i treatment followed comparable degradation kinetics and via a mechanism that was similarly UPS-dependent, but neddylation independent (**Fig 3B, C**). We thus hypothesized that TAK285 functions via the HSP90 regulatory axis.

**Fig 3.**
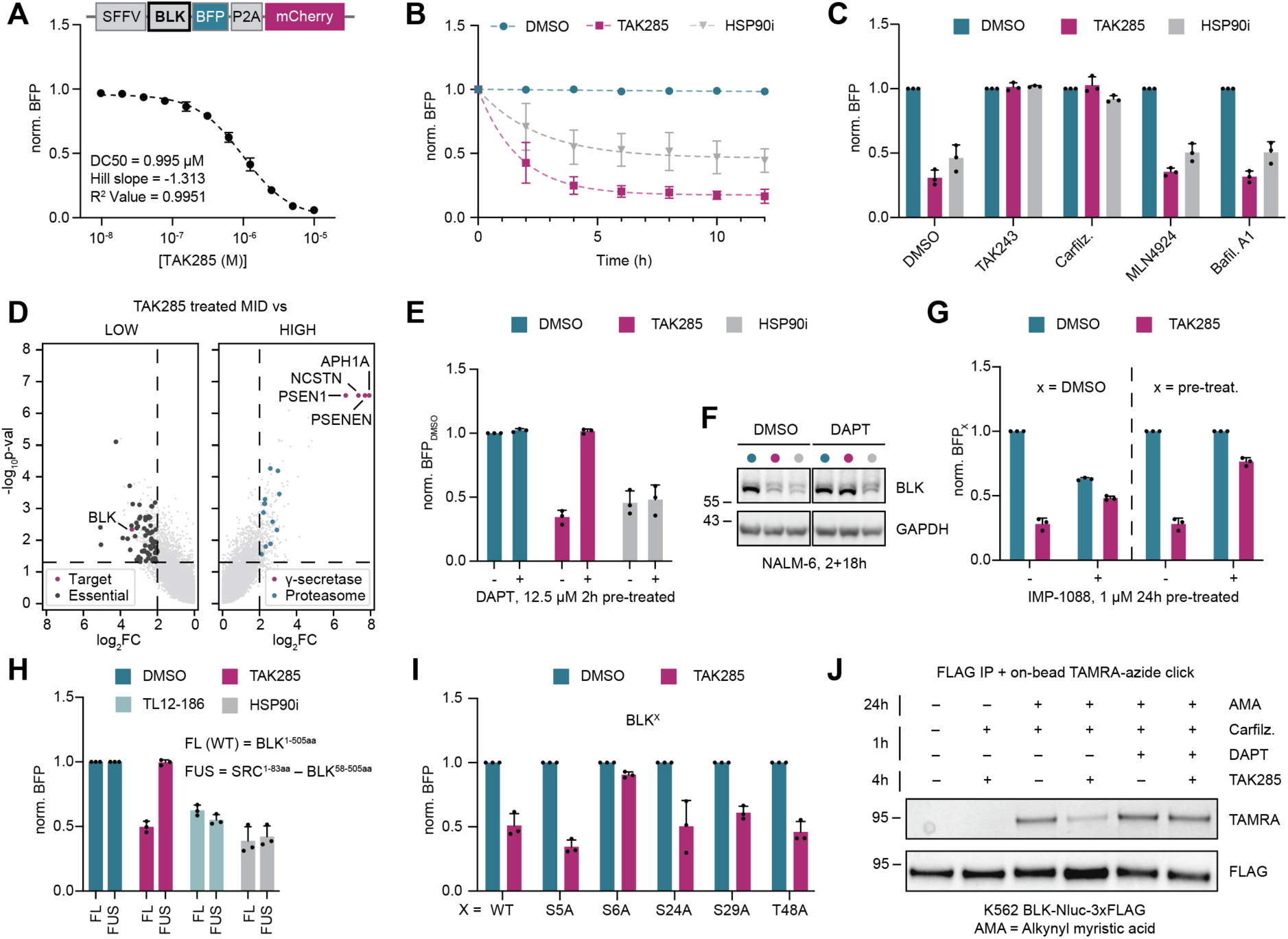
BLK is degraded by TAK285 in a γ-secretase-dependent manner. **A** Schematic of the stability reporter design used in the cell line background KBM7 iCas9. Dose response titration of TAK285 to determine the degradation of the BLK stability reporter as measured by flow cytometry. **B** Flow cytometric analysis of the time-dependent destabilization of BLK (stability reporter). TAK285 was treated at 2.5 µM and HSP90i at 10 µM. t^1/2^: TAK285 = 1.172h, HSP90i = 1.740h; D_max_: TAK285 = 82.49%, HSP90i = 53.68%, R^2^ value: TAK285 = 0.9994, HSP90i = 0.9902 (n = 3). **C** Chemical rescue of TAK285 and HSP90i mediated BLK stability reporter degradation (n = 3). TAK243 (1 µM), Carfilzomib (1 µM), MLN4924 (1 µM) and Bafilomycin A1 (Bafil. A1, 100 nM) were pre-treated for 2h, followed by 6h of 2.5 µM TAK285 treatment prior to flow cytometry. **D** Results of genome-wide FACS-based CRISPR/Cas9 screen. Essential genes (Depmap 23Q4^36^) and BLK or γ-secretase subunits and proteasome subunits are highlighted for a p-value < 0.05 and a log_2_ fold-change (FC) of > 2 for the low-to-mid and high-to-mid comparisons, respectively (n = 2). **E, F** Confirmation of the top four CRISPR/Cas9 screen hits using orthogonal chemical inhibition of the γ-secretase with DAPT (pre-treated for 2h at 12.5 µM). Degradation is abrogated for both the stability reporter system ((E), flow cytometry, 6h 2.5 µM TAK285 or 10 µM HSP90i) as well as on endogenous protein level ((F), immunoblot NALM-6 cells, 18h 2.5 µM TAK285 or 10 µM HSP90i) (n = 3). **G** Inhibition of myristoylation and measurement of BLK stability after TAK285 treatment. Myristoylation was inhibited 24h prior to 6h of 2.5 µM TAK285 treatment using the NMT1/2 inhibitor IMP-1088 at 1 µM. Left: shown as DMSO-normalized data. Right: normalized per pre-treatment (x; DMSO or IMP-1088, respectively). **H** BLK^wt^ (BLK^1-505aa^) or SRC unique domain fusion BLK (SRC^1-83aa^-BLK^58–505aa^) monitored in the stability reporter setup in KBM7 iCas9 cells and analyzed for degradation after 6h of treatment with TAK-285 (2.5 µM), TL12-186 (1 µM) and HSP90i (10 µM) using flow cytometry (n = 3). **I** Stability reporter mutant panel of all phospho-susceptible residues in the unique domain of BLK, assessed after 6h 2.5 µM TAK285 treatment using flow cytometry (n = 3). **J** Immunoprecipitation of BLK-Nluc-3xFLAG after pre-treatment with alkynyl myristic acid (AMA, 100 µM, 24h), followed by 1h Carfilzomib (1 µM) or additional DAPT (12.5 µM) incubation and treatments with DMSO or TAK285 (2.5 µM) for 4h. On-bead TAMRA click reaction and in gel-fluorescence shown in top (TAMRA) and FLAG immunoblot shown in bottom (FLAG) for immunoprecipitated (FLAG) fractions. Normalization of flow cytometry data was performed against the respective genotype or pre-treatment unless specified otherwise. Except for (F) and (J), all depicted experiments were performed in KBM7 iCas9 cells. All values represent the mean values ± SD; n = biological replicates, m = technical replicates.

To map the genetic determinants of TAK285-induced BLK degradation, we performed a FACS-based, genome-wide CRISPR/Cas9 screen (**Fig 3D**, **Extended Data Fig 4B**). Unexpectedly, we identified all four members of the γ-secretase complex (APH1A, NCSTN, PSEN1, PSENEN) as the most strongly enriched hits, suggesting a functional link to BLK degradation by TAK285. We validated involvement of the γ-secretase by pharmacological inhibition via DAPT (**Fig 3E, F**, **Extended Data Fig 4C**), as well as by genetic ablation (**Extended Fig 4D**). In contrast to TAK285, BLK degradation induced via HSP90i was independent of γ-secretase function. Hence, TAK285- induced BLK degradation appeared to be functionally differentiated from chaperone deprivation. Of note, γ-secretase subunits also scored as hits in steady-state (vehicle-) conditions of the genome-scale CRISPR/Cas9 screen (**Extended Data Fig 4B**), which we orthogonally confirmed by genetic ablation of *PSENEN* (**Extended Data Fig 4E**). This implied a role of γ-secretase in native BLK turnover. Further supporting a link between BLK and the γ-secretase complex, we identified an interaction between BLK and NCSTN when performing proximity labeling-based proteomics in BLK-mT-expressing KBM7 (**Extended Data Fig 4F**). Mining of orthogonal IP-MS data similarly revealed baseline interactions between NCSTN and BLK^41^. Amongst BLK’s interactors we identified further ADAM10 a well-known alpha-sheddase involved in pre-processing of γ- secretase substrates^42^, which is lost upon TAK285 treatment. Collectively, this data supports a functional and physical interaction between BLK and the γ-secretase upon TAK285 treatment and in steady state.

**Fig 4.**
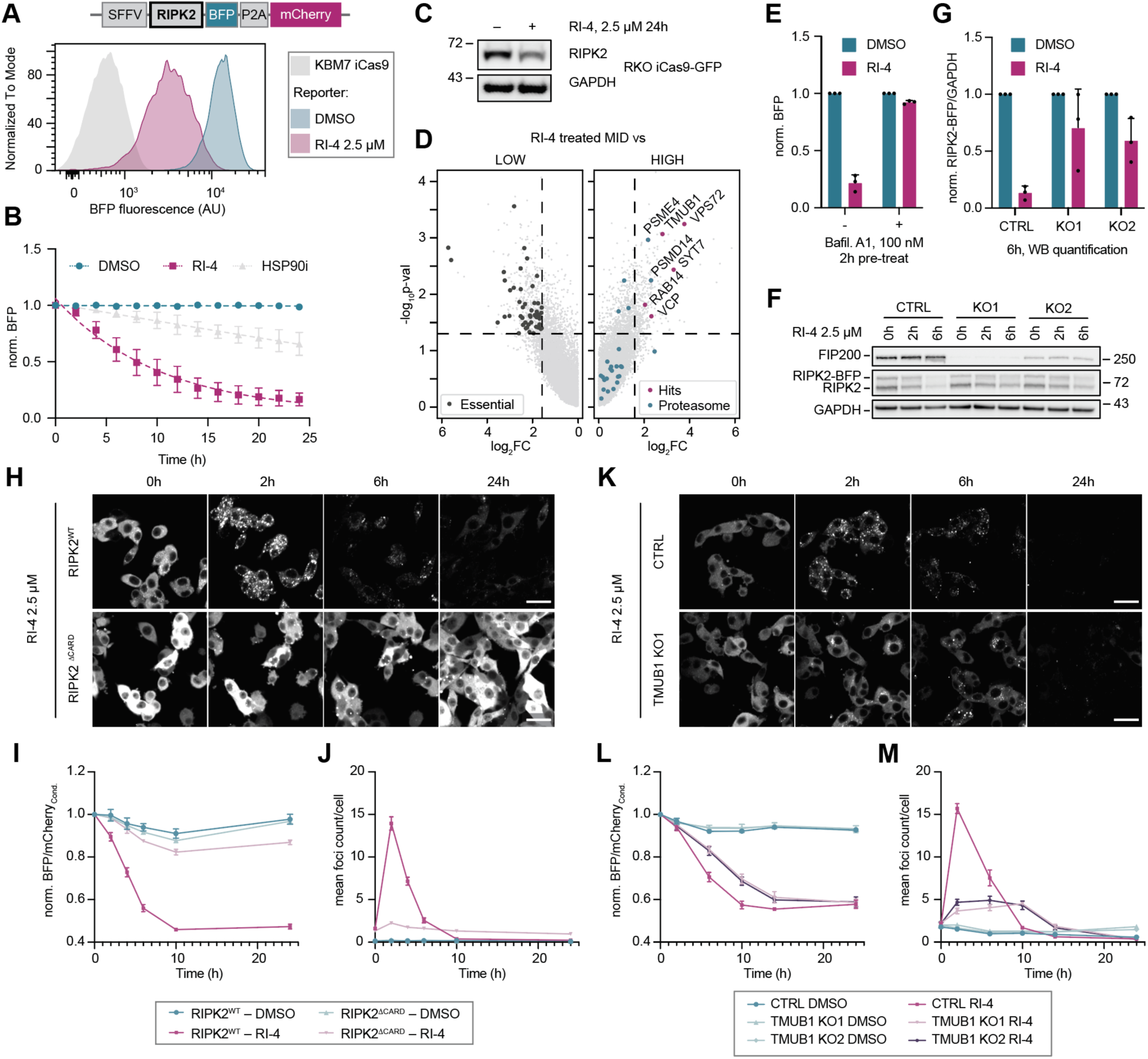
RI-4 destabilizes RIPK2 via TMUB-1-dependent multimerization and macroautophagy. **A** Stability reporter assessment of RI-4 dependent RIPK2 destabilization in KBM7 iCas9 reporter cells using flow cytometry after 18h of treatment at the indicated drug concentration. **B** Time-dependent RIPK2 stability reporter destabilization after cellular treatment with DMSO, 2.5 µM RI-4 or 10 µM HSP90i, assessed by flow cytometry (n = 3). **C** Immunoblot analysis of endogenous RIPK2 destabilization in RKO iCas9 cells (2.5 µM, 24h, n = 3). **D** Results of genome-wide FACS based CRISPR/Cas9 screen. Essential genes (Depmap 23Q4^36^) or selected hits are highlighted for a p-value (p-val) < 0.05 and a log_2_ fold-change (FC) of > 1.585 for the low-to-mid and high-to-mid comparisons, respectively. Proteasome subunits are additionally shown for the high-to-mid comparison irrespective of any cut-off (n = 2). **E** Bafilomycin A1 (Bafil. A1) pre-treatment impairs RI-4 (2.5 µM, 18h)-induced degradation as measured in the stability reporter system in KBM7 iCas9 via flow cytometry. **F** Representative immunoblot of RIPK2 stability reporter in RKO cells either expressing a control sgRNA (CTRL, sg*AAVS1*) or sgRNAs targeting *FIP200* k.o. (KO1/KO2). **G** Quantification of (F) and corresponding replicates (n = 3). **H** Representative microscopy images (BFP channel) of RIPK2^FL^ (full length) and RIPK2^ΔCARD^ stability reporter upon RI-4 treatment (2.5 µM) in RKO iCas9 cells. Scale bar = 50 µm, brightness adjusted per genotype (see **SI Fig S6B** for equally adjusted images and **Extended Data Fig 5F** for matched DMSO controls). **I, J** Quantification of RIPK2 stability (I) or mean number of RIPK2 foci per cell (J) as shown in H (n = 3, m = 2) normalized to the initial timepoint (0h) per condition. **K** Representative microscopy images (BFP channel) of RIPK2stability reporter upon RI-4 (2.5 µM) treatment in RKO iCas9 cells expressing a control sgRNA (CTRL, sg*AAVS1*) or a sgRNA targeting *TMUB1* (TMUB1 KO1) (see **Extended Data Fig 5J** for matched DMSO controls). Scale bar = 50 µm. **L, M** Quantification of RIPK2 stability (L) or mean number of RIPK2 foci per cell (M) as shown in K as well as TMUB1 KO2 and the respective DMSO controls (n = 3, m = 2). All values represent the mean values of the biological replicates ± SD; n = biological replicates, m = technical replicates.

The γ-secretase is an intramembrane protease best known for its involvement in Notch-1 and APP cleavage, albeit it also targets a selected set of kinases^42^. Hitherto cleavage is restricted to type I transmembrane proteins. Therefore, the myristoylated, membrane-anchored BLK is likely not a direct target of γ-secretase cleavage. However, we did not identify any candidate target(s) in our parallel BLK-mT and APH1A-mT BioID dataset, as no consensus type I transmembrane kinase/protein was enriched upon TAK285 treatment (**Extended Data Fig 4F**). We nevertheless surmised that differential membrane localization could underlie TAK285-induced BLK degradation. In line with this notion, perturbing membrane association of BLK via pharmacologic inhibition of N-myristoyltransferases 1/2 rescued TAK285-induced BLK degradation and, by itself, destabilized BLK (**Fig 3G**). A comparable membrane dissociation of BLK was also observed after TAK285 treatment (**Extended Data Fig 4G**). We thus concluded that TAK285 induces a membrane-to-cytosol transition of BLK, tapping into an intrinsically less stable state of the kinase. Myristoylation (and hence membrane-anchoring) of BLK and other SRC kinases occurs at the N-terminal unique domain^43^. In line with a critical role of BLK’s N-terminal domain in determining the specificity of the TAK285-induced degradation, we found that a domain swap to the N-terminal domain of SRC rescued TAK285-mediated degradation while degradation via HSP90i or BLK-targeting PROTAC TL12-186 was unaffected (**Fig 3H**). Of note, membrane association of the SRC-BLK chimera was retained (**Extended Data Fig 4I**).

Given this critical role of the N-terminal domain of BLK, we next addressed if TAK285-induced BLK degradation occurs via direct BLK binding. Cellular thermal shift assays (CETSA) did not show a thermal stabilization of BLK upon treatment with TAK285 and an *in vitro* assay revealed only partial kinase inhibition (≈ 37%, IC50 > 30 µM; **SI Fig S5F, G**). Thus, we next turned to chemical competition experiments. Blocking the active site of BLK via the covalent inhibitor acalabrutinib rescued degradation induced by the BLK-targeting PROTAC TL12-186 (**Extended Data Fig 4J**). Expectedly, this rescue was dependent on Cys319 in BLK, which is covalently engaged by acalabrutinib^23^. In contrast, TAK285-induced BLK degradation was unaffected by acalabrutinib, suggesting a mechanism that is independent of direct binding to the ATP pocket of BLK. To uncover the potential intermediate kinase targeted by TAK285, we again performed dose-ranging chemoproteomics. The standard cell lysate mix identified three binding partners: the established TAK285 target EGFR, a type-I transmembrane kinase, which is, however, not considered a γ-secretase substrate^42^, the kinase MAP2K5 (MEK5) and the non-kinase protein ERCC2 (**SI Fig S5H**). Since BLK was not detected in this standard setup, we expanded our search using an alternative cell lysate mix, which accounts for the tissue-specific expression of BLK (see **SI Material and Methods**). While BLK was now detected, TAK285 did not bind BLK (**Extended Data Fig 4K**). Collectively, these data suggested that TAK285 induces BLK re-localization independent of orthosteric BLK binding, presumably involving an additional effector still to be uncovered in future work.

Next, we focused on the role of BLK’s N-terminal unique domain to further our understanding of this process. Mutating all residues that could directly be modified via upstream phosphorylation networks led us to identify that BLK^S6A^ strongly abrogated inhibitor-induced degradation (**Fig 3J**). The conserved Ser6 residue had previously been associated with regulating myristoylation and membrane association of the related SRC-family kinase LCK^44^. Accordingly, BLK^S6A^ lost membrane association, appeared predominantly cytoplasmatic and had a lower baseline stability (**Extended Data Fig 4I, L**), phenocopying the effects of myristoylation inhibition via IMP- 1088. We thus hypothesized that TAK285 may induce a change of BLK’s myristoylation status. To test this hypothesis, we modified a previously reported method^45^ to utilize alkynyl myristic acid (AMA) as chemical probe to monitor this N-terminal myristoylation of BLK. Intriguingly, TAK285 treatment led to a loss of myristoylation, which was rescued upon γ-secretase inhibition (**Fig 3J**).

In conclusion, we found that TAK285 induces a γ-secretase-dependent dissociation of the membrane-associated BLK into the cytoplasm where BLK is intrinsically instable. TAK285- induced destabilization is independent of direct BLK binding but is encoded by its unique N-terminal domain and critically mitigated by its myristoylation status.

### RIPK-IN-4 destabilizes RIPK2 by prompting lysosomal degradation

As the final example, we turned our attention to RIPK2, a cytoplasmic kinase involved in the clearance of bacterial pathogens by linking activation of the pattern recognition receptors NOD1/NOD2 to intracellular signaling^46^. Kinome abundance trajectories revealed nine inhibitors potentially destabilizing RIPK2. Among those inhibitors, RIPK-IN-4 (RI-4, **Fig 1G**) prompted the most selective and potent degradation response (**Extended Data Fig 5A**) and was hence selected for further mechanistic workup. In accordance with RI-4’s role as RIPK2 inhibitor^28^, recombinant binding assays confirmed RIPK2 engagement, suggesting a directly induced degradation event (**Extended Data Fig 5B**). Leveraging a RIPK2 stability reporter, we next validated that RI-4 functions at the level of protein stability, inducing RIPK2 degradation over a timeframe of approximately 18h (**Fig 4A, B**). Next, we confirmed degradation of endogenous RIPK2 via immunoblotting (**Fig 4C**). Quantitative expression proteomics confirmed RI-4’s selectivity that was suggested by our dynamic abundance profiles. In addition to RIPK2, CSK emerged as the only other destabilized kinase (**Extended Data Fig 5C**). To reveal cellular effectors required for RIPK2 degradation induced by RI-4, we again turned to a time-controlled, genome-scale FACS-based CRISPR/Cas9 screen, utilizing the RIPK2 stability reporter as readout. Systems-level analysis of the identified effector genes revealed an enrichment of hits that are involved in lysosomal degradation (**Fig 4D**). Moreover, we identified the Ubiquitin-like (UBL) domain containing TMUB1 among the most strongly enriched genes. TMUB1 has previously been implied to be involved in quality control of transmembrane proteins at the ER contact site^47^. In line with the screening data, pharmacologic inhibition of lysosome acidification via Bafilomycin A1 rescued RIPK2 degradation by RI-4 (**Fig 4E**). Moreover, we found that RI-4-induced RIPK2 degradation was abrogated in cells deficient for the well-established macro-autophagy mediator FIP200^48^, but not upon population-level knockout of the proteasomal subunit PSMB5 (**Fig 4F, G**, **Extended Data Fig 5D, E**). In sum, these data supports that RIPK2 degradation by RI-4 depends on macro-autophagy.

Aiming to investigate RI-4-induced changes in RIPK2 abundance via microscopy, we observed that RIPK2 assembled into clearly discernable foci prior to degradation (**Fig 4H-J**). These structures were reminiscent of higher order RIPK2 oligomers, which are reported to form after endogenous activation, commonly referred to as RIPosomes^49^. A key feature of RIPosome formation is the essential role of RIPK2’s CARD domain^50^. Testing the effect of RI-4 on a CARD domain deficient RIPK2 variant (RIPK2 ΔCARD^aa437–524^) revealed that formation and clearance of the RI4-induced RIPK2 assemblies are likewise CARD domain dependent, thus implying a functional resemblance of both assemblies (**Fig 4H-J**). In line with observations made via FACS-based RIPK2 stability reporters, we found that clearance of RI-4-induced RIPK2 assemblies was rescued via Bafilomycin A1 treatment (**Extended Data Fig 5G**-I). This again highlighted the similarity to RIPosomes, which have likewise been shown to be turned over by autophagy^49^. To connect TMUB1 to this process, we analyzed RI-4-induced RIPK2 assembly and degradation in TMUB1 deficient cells. This revealed a marked delay in the assembly formation, as well as the total number of observed foci, and consequently a delayed degradation (**Fig 4K-M**). This implies TMUB1 as an early-acting facilitator of this process. Taken together, our data supports a model where RI-4 induces higher-order assemblies of RIPK2 via involvement of the UBL-domain containing protein TMUB1. These assemblies mimic multimers that are formed in response to physiologic stimuli by pathogens. Pathogen-induced, as well as inhibitor-induced assemblies are subsequently turned over via macro-autophagy.

## Discussion

Global and focused analysis of dynamic abundance profiles of 98 kinases after cellular exposure to 1570 annotated kinase inhibitors revealed that inhibitor-induced kinase degradation is a frequent phenomenon. Known HSP90 clients are enriched among the degradation sensitive kinases, suggesting that chaperone deprivation is likely a widespread mechanism of inhibitor-induced degradation. However, in-depth mechanistic investigation of three degradation events of kinases with graded HSP90 dependency revealed a differentiated, yet shared mechanism of action. In all cases, inhibitor-induced kinase degradation leveraged and further elevated physiological turnover mechanisms by inducing kinase states that are primed for degradation. Mechanistically, different phenomena can manifest in these instable kinase states, including altered kinase activity (LYN), changed cellular localization (BLK), or induced higher-order assemblies (RIPK2).

For the investigated kinase inhibitors, we found that induction of these states can be triggered by direct target engagement, or via network-drugging. This highlights that potent and selective off-target degradation can also result from on-target inhibition, suggesting that unbiased profiling of drug-impact on proteome abundance can, in addition to unbiased measurements of target engagement, inform on additional therapeutic opportunities or potentially unwanted side effects. Indeed, systematic proteomics profiling campaigns have revealed a stunning breadth of proteome-wide effects^51,52^. Akin to a proteomics readout, our assay informs on protein abundance and not directly on protein stability, which necessitated implementation of a set of control compounds to exclude global effects on transcription or translation. Additionally, we found that conducting time-resolved measurements vitally informed the identification of compounds that act on protein stability, akin to observations made for proteomics profiling^53^.

Globally, we do not detect a pronounced overlap between kinases that are prone to be destabilized by inhibitors and kinases that are primed for degradation via proximity-inducing modalities, such as PROTACs. Nevertheless, there is evidence that also proximity-inducing molecular glue degraders can be prospectively furnished to re-establish a physiological degradation mechanism that is disrupted by mutations, as exemplified by molecular glue degraders that induce proximity between mutant b-catenin and its physiological ligase b-TrCP^54^.

Future research will be required to understand the scope of supercharging endogenous degradation events beyond kinases. In addition to BCL6 for which BI-3802-induced multimerization enables degradation via its native E3 ligase SIAH1, a parallel manuscript reports directly acting, monovalent degraders of IDO1 (Hennes et al., 2024)^19^. IDO1 is an enzyme involved in tryptophan metabolism that has been clinically pursued as an immune-oncology target. Mechanistically, the reported ligands induce a conformational change that stabilizes IDO1 in the apo state that is not bound by heme and therefore more susceptible to turnover by CRL2^KLHDC3^, the E3 in charge of its endogenous turnover.

Collectively, these studies indicate supercharging of physiological degradation routes as a general mechanism of ligand-induced protein degradation that are complementary to proximity-inducing modalities such as PROTACs or molecular glue degraders.

## Author contributions

N.S.S. and G.E.W. conceptualized the study and wrote the manuscript with input from all authors. N.S.S. designed and executed most of the described experiments, data analysis and generated the figures. M.B. and A.C-C. generated the drug screen scoring scheme. P.A. supervised the corresponding data analysis. K.K. performed experiments, quantified immunoblots and handled BioID experiments. F.F. assisted with the BioID experiments, performed the data analysis and generated the respective figure panels. A.R. handled expression proteomics sample preparation and processing. E.B. and H.R. performed experiments. A.K. assisted with the drug screen, performed the initial data processing and normalization. M.H. supported CRISPR/Cas9 screens. M.P.S. performed NanoBRET experiments. P.P. performed and analyzed the Kinobead profiling. H.I. and C.S. analyzed sequencing data. S.Ku. supervised the drug screen. S.Kn. oversaw the NanoBRET assays. N.S.G. gave critical input to the manuscript. G.S.F. and B.K. supervised the proteomics experiments. G.E.W. has overall responsibility for the presented study.

## Acknowledgements

We are grateful to all the members of the Winter lab, in particular Marko Cigler for helpful discussions and editorial contributions, as well as Maiia Schulmann for assistance in DNA preps. We thank the Core Facility Flow Cytometry of the Medical University of Vienna for access to flow cytometry instruments and assistance with flow cytometric cell sorting as well as the CeMM Biomedical Sequencing Facility for NGS sample sequencing. We thank Thomas Hannich and Iciar Serrano of the MDP ProMet-facility at CeMM for excellent technical assistance for proteomics data acquisition. We moreover thank Johannes Zuber at the Research Institute of Molecular Pathology for sharing iCas9 cell lines and plasmids as well as Anne-Claude Gingras for providing the BioID plasmids. We would also like to thank Sascha Martens and Mads Gyrd-Hansen for helpful discussions.

CeMM, the Winter lab and the Superti-Furga lab are supported by the Austrian Academy of Sciences. The Winter lab is further supported by funding from the European Research Council (ERC) under the European Union’s Horizon 2020 research and innovation program (grant agreement 851478), as well as by funding from the Austrian Science Fund (FWF, projects P7909, P36746 and P5918723). N.S.S. is further supported by the FWF postdoctoral Esprit fellowship ESP 426 and Marie Skłodowska-Curie postdoctoral fellowship (grant agreement number: 101029199). P.A. acknowledges the support of the Generalitat de Catalunya (RIS3CAT Emergents VEIS: 001-P-001647 and 2021 SGR 00876) and the Spanish Ministerio de Ciencia, Innovación y Universidades (PID2020-119535RB-I00). A.C-C. is a recipient of an FI fellowship (2020 FI_B 00094). S.Kn. and M.P.S. are grateful for support by the Structural Genomics Consortium (SGC), a registered charity (no: 1097737) that receives funds from Bayer AG, Boehringer Ingelheim, Bristol Myers Squibb, Genentech, Genome Canada through Ontario Genomics Institute, EU/EFPIA/OICR/McGill/KTH/Diamond Innovative Medicines Initiative 2 Joint Undertaking [EUbOPEN grant 875510], Janssen, Pfizer and Takeda. S.Kn. is also supported by the German Cancer Research Center DKTK, the German Cancer Aid project TACTIC and the Frankfurt Cancer Institute (FCI). M.P.S. is funded by the Deutsche Forschungsgemeinschaft (DFG, German Research Foundation), CRC1430 (Project-ID 424228829).

## Data availability

Screening data will be made available upon publication on an interactive website to enable easy access. All processed sequencing data and proteomics data has been made available as supplementary documents (**Supplementary Data Table 2-5**). Full access to the proteomics datasets will be made available upon publication.

---

**Extended Data Fig 1.**
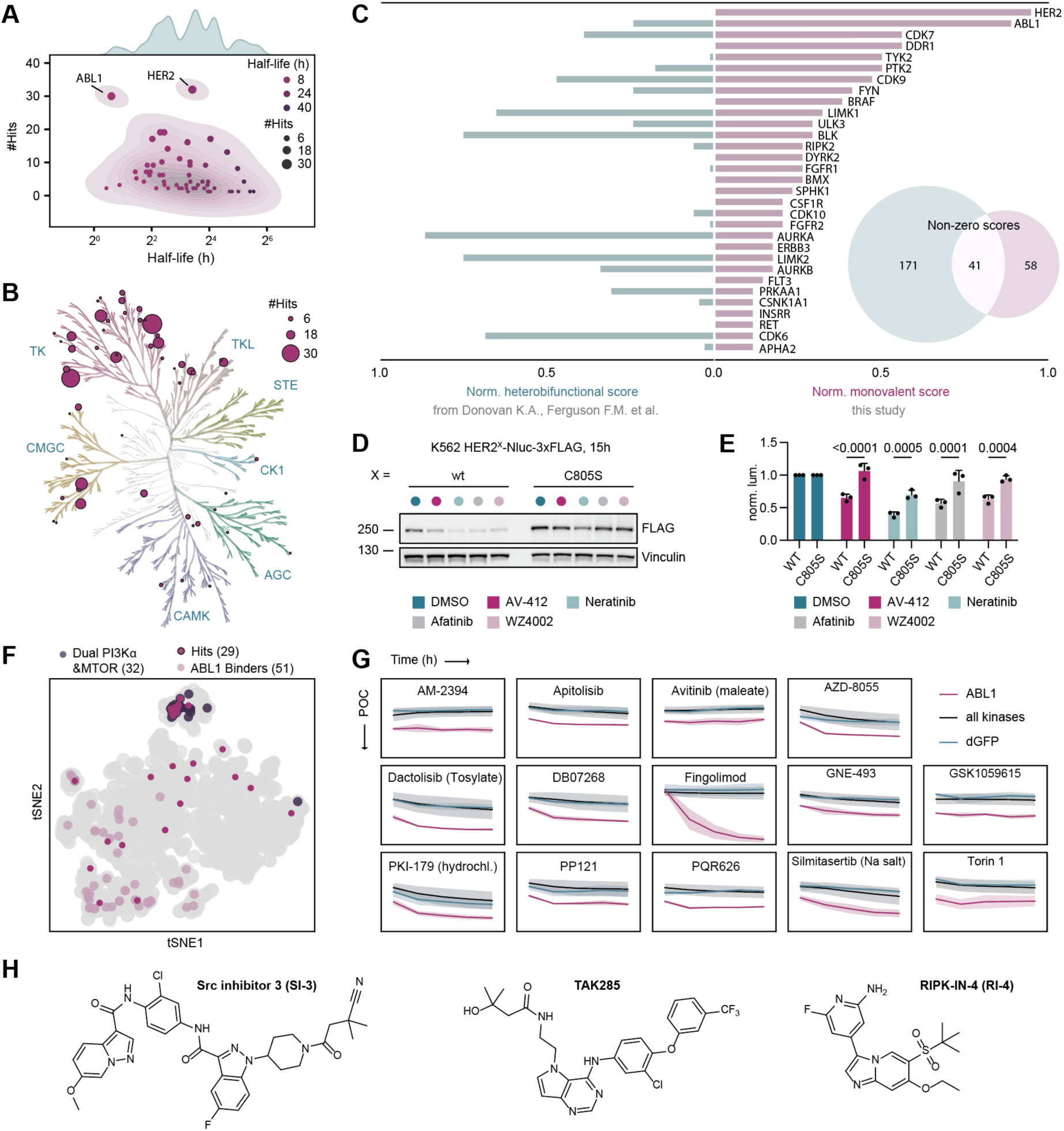
Neither half-life nor heterobifunctional degradability determine monovalent destabilization frequencies. **A** Comparison of half-life per each kinase fitted from CHX screening data plotted against the summed hit scores. Top: Half-life KDE plot with adjusted bw value of 0.5. Middle: 2D KDE plot and scatter plot of number of hits (#Hits = score) and half-life (h). **B** Summed destabilization score (#Hits) per canonical kinase mapped onto the kinome tree. **C** Comparison of monovalent degrader scores to previously reported scoring frequencies by heterobifunctional degrader molecules (heterobifunctional score) adapted from Donovan, et al^18^. The data was normalized to the highest scoring kinase per study and depicts the sorted top 32 downregulated kinases by monovalent small molecules. In total, 41 kinases were detected to be downregulated in at least one instance for both studies. **D** Western blot analysis of cell lines in E (n = 3). **E** Luminescent reporter assay of K562 HER2^wt^ or HER2^C805S^ Nluc-3xFLAG reporter cell lines treated for 15h with the indicated compounds (all 10 µM except for AV-412 (2.5 µM)) shown as normalized luminescence per genetic construct. Statistics were calculated using a two-way Anova with Šídák’s multiple comparisons test (n = 3). **F** tSNE plot of compound target landscape focusing on ABL1 hits in comparison to annotated ABL1 binders or dual PI3Kα and MTOR binders. **G** Drug screening data comparing ABL1 to the mean of all other kinases and the dGFP control. Continued in **SI Fig S2** (m = 2, error bars correspond to CI for individual trajectories and SD for the mean of all kinases). **H** Chemical structures of the three selected examples for mechanism of action elucidation. All data shown as mean of the replicates ± SD unless specified otherwise; n = biological, m = technical replicates.

**Extended Data Fig 2.**
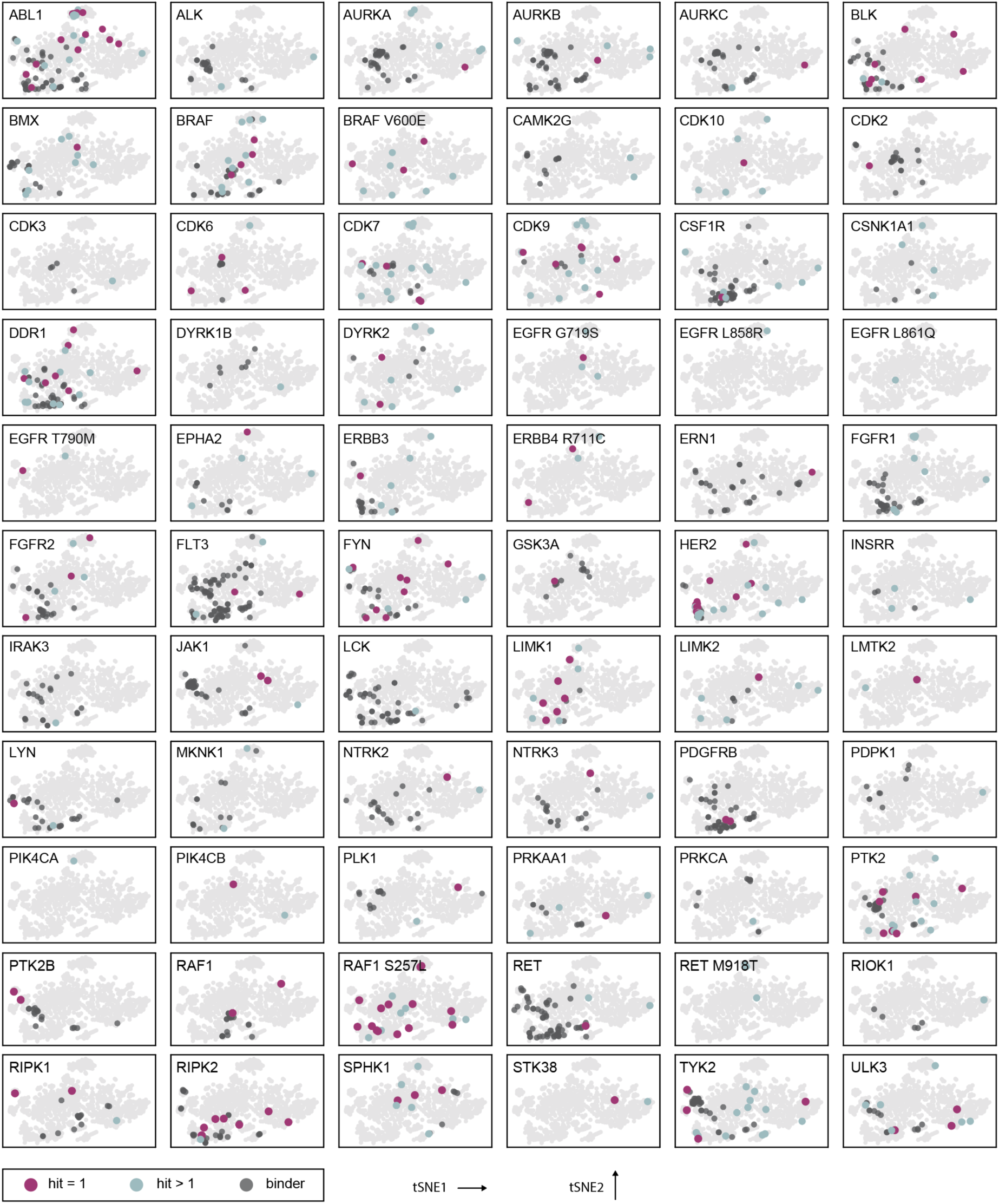
Hit and drug binding profile depicted as tSNE. Overlay of the target compound space of all screened inhibitors as annotated in ChEMBL (light grey, see **Supplementary Data Table 1**) with all identified hits and annotated kinase binders per kinase shown in the indicated color code.

**Extended Data Fig 3.**
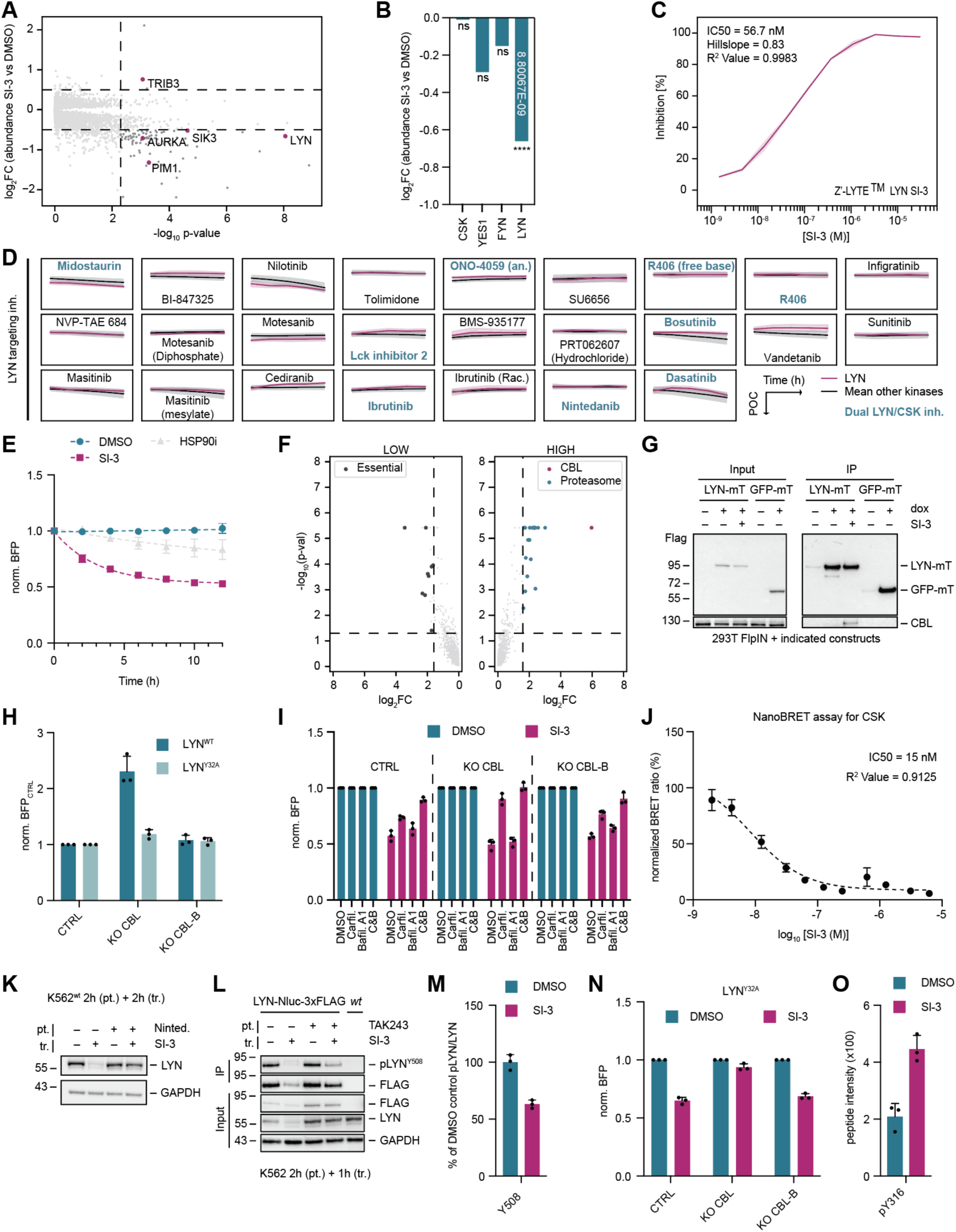
SI-3 selectively destabilizes LYN via a dual phosphodegron and dual E3 mechanism of action. **A** Expression proteomics of K562 cells after 2h of SI-3 treatment (156 nM) (n = 3). **B** Highlighted Src kinase family members and CSK abundance changes including p-values for the expression proteomics experiment shown in (A). **C** Z-LYTE’ SI-3 Lyn inhibitory assay (m = 2, error bars = CI). **D** Drug screen trajectories of compounds annotated as LYN binders (**Supplementary Data Table 1**) depicted for the LYN reporter (pink) and compared to the mean response of all other kinase reporters (black). Compounds that are annotated to bind both LYN and CSK (ChEMBL) have further been highlighted in green. Axis annotations: y-limits (POC) = 0-150 and x-limits (Time (h)) = 0-20; LYN error bars = CI and all other trajectories are shown as SD. **E** Temporal destabilization profile of LYN stability reporter for DMSO, SI-3 (156 nM) or HSP90i (10 µM) (n = 3). **F** UPS-focused CRISPR/Cas9 screen data for vehicle (DMSO) treated cells (n = 2). **G** Immunoblot of representative enrichment for the respective BioID conditions (**Fig 2F**) (n = 3). **H** Genetic k.o. (CTRL = sg*AVVS1*, KO CBL or KO CBL-B) of LYN^WT^ or LYN^Y32A^ stability reporter cell lines measured by flow cytometry. Depicted are baseline normalized values to each respective CTRL sample (n = 3). **I** Chemical rescue upon genetic k.o. for sg*AAVS1* (CTRL) or single CBL/CBL-B KO. Cells were pre-treated for 2h with 1 µM Carfilzomib (Carfil.) or 100 nM Bafilomycin A1 (Bafil. A1) or both (C&B) followed by 8h of 156 nM SI-3 treatment and flow cytometric analysis (n = 3). **J** NanoBRET measurement of tracer displacement from CSK by SI-3 (n = 2, m = 2). **K** Immunoblot analysis of K562 cells pre-treated with Nintedanib (10 µM, 2h) followed by 2h of SI-3 (156 nM) treatment (n = 3). **L** FLAG immunoprecipitation (IP) of K562 Lyn-Nluc-3xFLAG or parent (*wt*) control upon 2h pre-treatment (pt.) with 1 µM TAK243 or DMSO followed by 1h of DMSO or SI-3 treatment (n = 3). **M** Quantification of (L) using TAK243 pre-treated (degradation impaired) samples and comparing pY508 LYN to total LYN. The quantification was performed on the IP fractions (FLAG IP) for the bands corresponding to LYN-Nluc-3xFLAG and normalized to the DMSO control lane (n = 3). **N** Genetic k.o. (CTRL = sg*AVVS1*, KO CBL or KO CBL-B) for LYN^Y32A^ stability reporter cell line and SI-3 degradation assessment (8h, 156 nM) measured by flow cytometry (n = 3). **O** Phospho-peptide quantification of LYN pY316 measured in the BioID dataset and shown as raw peptide intensity (n = 3). n = biological replicates, m = technical replicates, all data shown as mean of replicates ± SD unless specified otherwise.

**Extended Data Fig 4.**
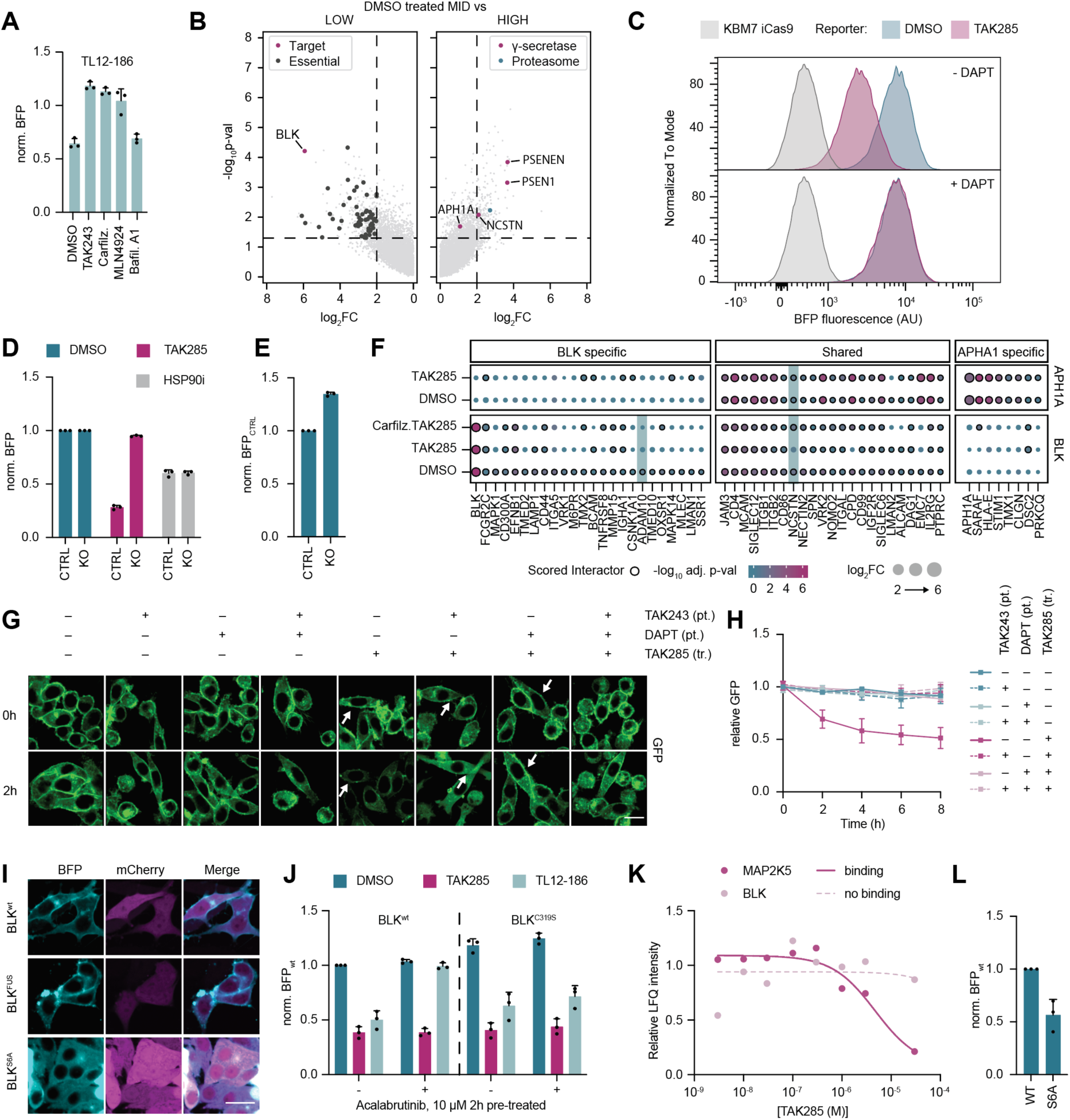
TAK285 re-localizes and degrades BLK in a γ-secretase dependent manner. **A** Scheme as in **Fig 3C** but shown for TL12-186 (1 µM, 6h) (n = 3). **B** Matched genome-wide CRISPR/Cas9 data to **Fig 3D** shown for vehicle control (DMSO) (n = 2). **C** Representative BLK stability reporter flow cytometry histogram plot for 2h DMSO or 12.5 µM DAPT pre-treated cells followed by 6h 2.5 µM TAK285 treatment. **D** Genetic k.o. of *PSENEN* (KO; γ-secretase subunit) treated for 6h with DMSO, TAK285 (2.5 µM) or HSP90i (10 µM) followed by flow cytometric analysis (n = 3). **E** *PSENEN* k.o. (KO) baseline BLK stability values measured by flow cytometry and depicted normalized to CTRL (sg*AAVS1*) (n = 3). **F** Kinase and type I transmembrane protein interaction partners for BLK and APH1A mapped by miniTurbo (mT) BioID. Interaction partners were scored in baseline (DMSO) conditions against GFP negative controls and grouped if they were shared between the baits or specifically found in BLK (BLK specific) or APH1A (APH1A specific). The type I transmembrane proteins ADAM10 and the γ-secretase subunit NCSTN are highlighted with a green shade. Interactors were further ordered in descending log_2_FC of BLK DMSO vs GFP for the BLK specific and shared interactions, and according to the descending log_2_FC of APH1A DMSO vs GFP for the APH1A specific. Dot size corresponds to log_2_FC of each protein against the GFP control. Black dot outlines indicate significantly scored interaction partners within each condition. Color gradient represents the - log_10_ of the adjusted p-value (adj. p-val, BH) (n = 3). **G** Representative images for the timepoints 0h and 2h for a BLK-GFP clonal cell line (RKO iCas9-BFP) assessed for different treatment conditions. Cells were pre-treated (pt.) for 1h with TAK243 (1 µM) and DAPT (12.5 µM) followed by DMSO or TAK285 (2.5 µM) treatment (tr.) for the indicated timeframe. White arrows highlight localization patterns. Scalebar = 25 µm (n = 2 (clonal cell lines), m = 2). See **SI Fig S5E** for additional example. 3 fields (> 90 cells) per well and 2 wells per replicate were assessed. **H** Quantification of relative GFP intensity from microscopy images shown in (G) and additionally recorded timepoints. The mean total (whole-cell) GFP abundance (y-axis) was normalized per DMSO or TAK243 pre-treatment at 0h and monitored for a total timeframe of 8h (x-axis) (n = 2 (clonal cell lines), m = 2). **I** Representative images for localization of different BLK stability reporter pools (BFP- P2A-mCherry) in RKO iCas9 GFP cell line from **Fig 3H, J**. Scalebar = 25 µm. **J** Chemical competition using 2h of 10 µM Acalabrutinib pre-treatment followed by 6h of 2.5 µM TAK285 or 1 µM TL-12-186 treatment and analysis by flow cytometry across two stability reporters: BLK^wt^ or the cysteine mutant BLK^C319S^ (n = 3, depicted normalized to BLK^wt^ as indicated in the axis label). **K** Dose-ranging chemoproteomics (Kinobead profiling) in Jurkat and MCF7 cell lysates. MAP2K5 was identified as only binder (solid pink line, K_D_ = 2.55 µM), while no binding was detected for BLK (dashed light pink line). **L** Baseline stability values for BLK^wt^ or BLK^S6A^ stability KBM7 iCas9 reporters measured by flow cytometry and depicted normalized to BLK^wt^ (n = 3). All values represent the mean values ± SD; n = biological replicates, m = technical replicates. Flow cytometry was measured in cell lines derived from KBM7 iCas9, while imaging was performed in reporter cell lines generated from RKO iCas9-GFP or RKO iCas9-BFP. BioID experiments were performed in KBM7 cells.

**Extended Data Fig 5.**
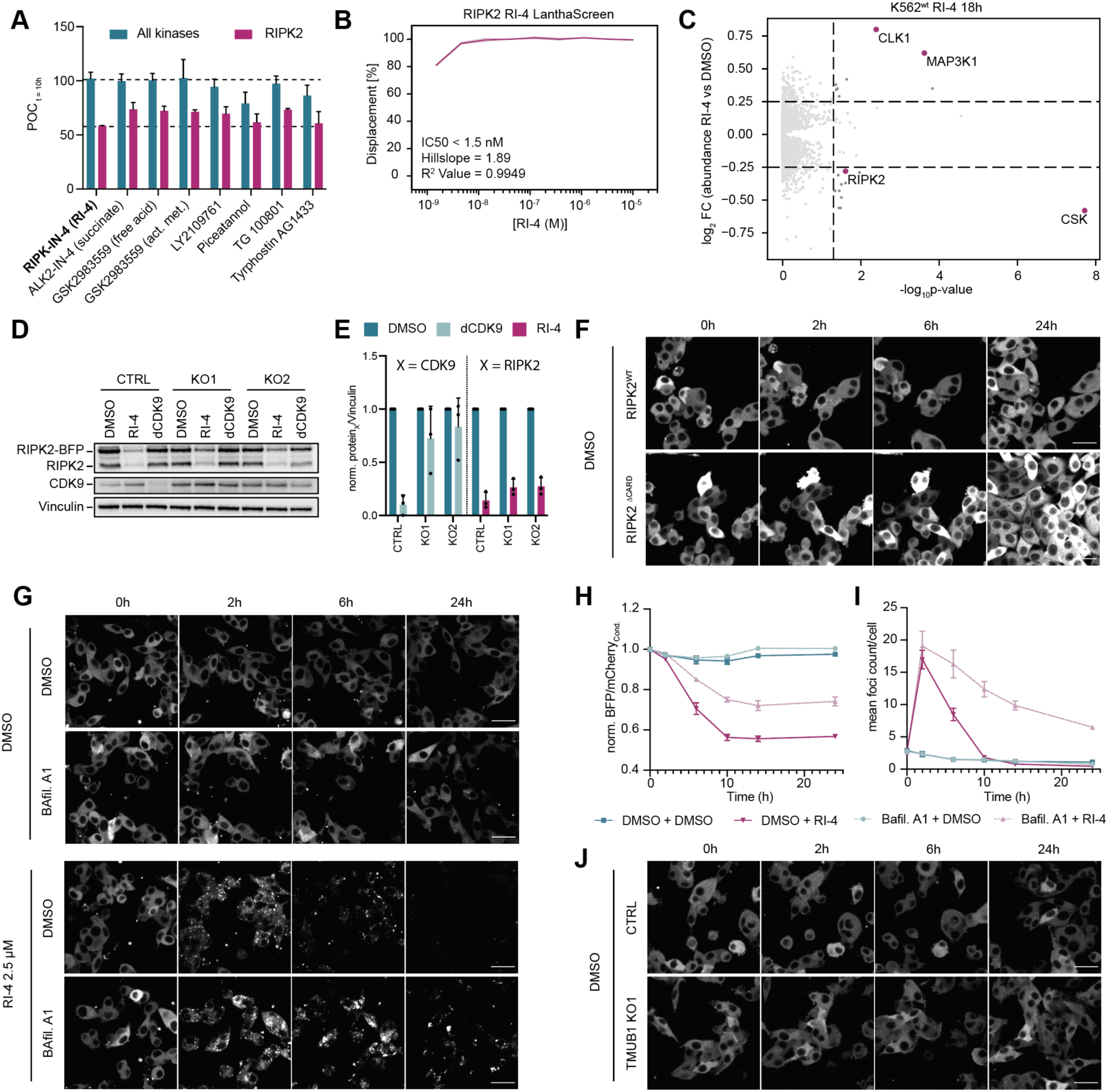
RI-4 degrades RIPK2 by inducing kinase multimerization, leading to its clearance via the lysosomal machinery. **A** Drug screen data shown as mean values ± SD for all kinases vs RIPK2 only for the 10h timepoint. Top and bottom dotted line indicated RI-4’s degradation window (m = 2). **B** RI-4 LanthaScreen drug binding data. (m = 2, error bar = CI). **C** Expression proteomics of 10 µM RI-4 treated K562 for 18h (n = 3). **D** Example immunoblot of *PSMB5* genetic k.o. of RKO iCas9 RIPK2-BFP stability reporter harboring cells expressing either a control sgRNA (CTRL, sg*AAVS1*) or sgRNAs targeting *PSMB5* (KO1/KO2) treated for 6h with either DMSO, 2.5 µM RI-4 or 1 µM dCDK9. **E** Quantification of (D) and respective replicates for CDK9 or RIPK2 abundance for either dCDK9 or RI-4 treated samples, respectively. Values are shown in relation to Vinculin and normalized to DMSO (n = 3). **F** Representative microscopy images (BFP channel) of RIPK2^FL^ (full length) and RIPK2^ΔCARD^ stability reporter in RKO iCas9 cells for vehicle control (DMSO) treatments matched treatments shown in **Fig 4H-J**. **G** Representative microscopy images (BFP channel) of RIPK2 stability reporter upon RI-4 treatment in RKO iCas9 cells for either DMSO or Bafilomycin A1 (Bafil. A1) pre-treated (2h) cells followed by DMSO or RI-4 (2.5 µM) treatment for a total timeframe of 24h. **H/I** Quantification of RIPK2 stability (H) or mean number of RIPK2 foci per cell (I) as shown in G (n = 3, m = 2). **J** Representative microscopy images (BFP channel) of RIPK2 stability reporter in RKO iCas9 cells for CTRL or TMUB1 KO1 vehicle control (DMSO) treatments matched to RI-4 treatments shown in **Fig 4K-M**. All values represent the mean values ± SD unless specified otherwise; n = biological replicates, m = technical replicates. Scale bars = 50 µm.

**Extended Data Table 1:**
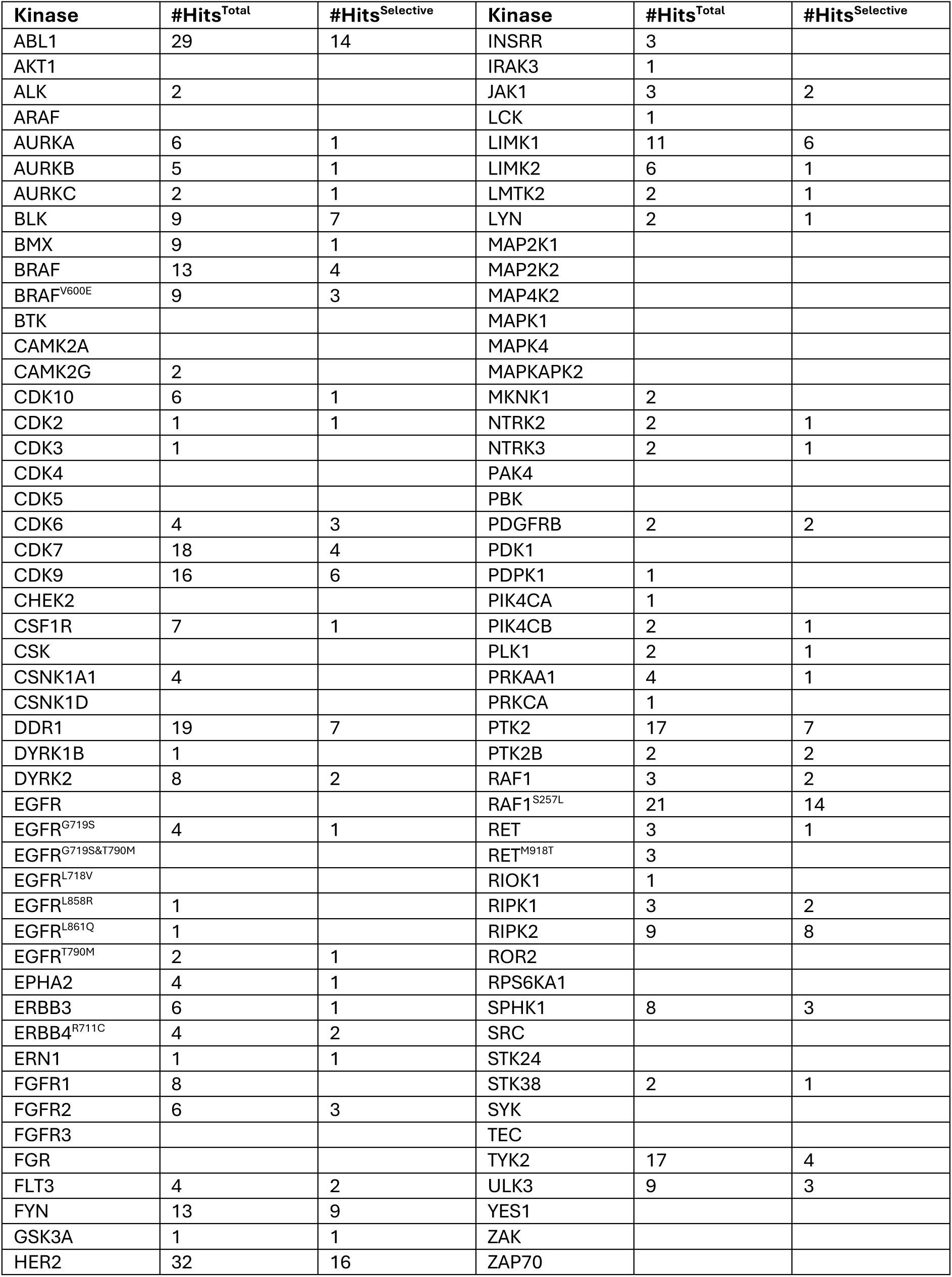
Screened kinases and hit number breakdowns. The kinase column indicates the kinase screened as Nluc-3xFLAG reporter cell line in K562. Number of hits (#Hits) are indicated for the respective total and selective (only one kinase scoring per compound) counts across 1570 compounds.

| Kinase | #Hits<sup>Total</sup> | #Hits<sup>Selective</sup> | Kinase | #Hits<sup>Total</sup> | #Hits<sup>Selective</sup> |
| --- | --- | --- | --- | --- | --- |
| ABL1 | 29 | 14 | INSRR | 3 |  |
| AKT1 |  |  | IRAK3 | 1 |  |
| ALK | 2 |  | JAK1 | 3 | 2 |
| ARAF |  |  | LCK | 1 |  |
| AURKA | 6 | 1 | LIMK1 | 11 | 6 |
| AURKB | 5 | 1 | LIMK2 | 6 | 1 |
| AURKC | 2 | 1 | LMTK2 | 2 | 1 |
| BLK | 9 | 7 | LYN | 2 | 1 |
| BMX | 9 | 1 | MAP2K1 |  |  |
| BRAF | 13 | 4 | MAP2K2 |  |  |
| BRAF<sup>V600E</sup> | 9 | 3 | MAP4K2 |  |  |
| BTK |  |  | MAPK1 |  |  |
| CAMK2A |  |  | MAPK4 |  |  |
| CAMK2G | 2 |  | MAPKAPK2 |  |  |
| CDK10 | 6 | 1 | MKNK1 | 2 |  |
| CDK2 | 1 | 1 | NTRK2 | 2 | 1 |
| CDK3 | 1 |  | NTRK3 | 2 | 1 |
| CDK4 |  |  | PAK4 |  |  |
| CDK5 |  |  | PBK |  |  |
| CDK6 | 4 | 3 | PDGFRB | 2 | 2 |
| CDK7 | 18 | 4 | PDK1 |  |  |
| CDK9 | 16 | 6 | PDPK1 | 1 |  |
| CHEK2 |  |  | PIK4CA | 1 |  |
| CSF1R | 7 | 1 | PIK4CB | 2 | 1 |
| CSK |  |  | PLK1 | 2 | 1 |
| CSNK1A1 | 4 |  | PRKAA1 | 4 | 1 |
| CSNK1D |  |  | PRKCA | 1 |  |
| DDR1 | 19 | 7 | PTK2 | 17 | 7 |
| DYRK1B | 1 |  | PTK2B | 2 | 2 |
| DYRK2 | 8 | 2 | RAF1 | 3 | 2 |
| EGFR |  |  | RAF1<sup>S257L</sup> | 21 | 14 |
| EGFR<sup>G719S</sup> | 4 | 1 | RET | 3 | 1 |
| EGFR<sup>G719S&T790M</sup> |  |  | RET<sup>M918T</sup> | 3 |  |
| EGFR<sup>L718V</sup> |  |  | RIOK1 | 1 |  |
| EGFR<sup>L858R</sup> | 1 |  | RIPK1 | 3 | 2 |
| EGFR<sup>L861Q</sup> | 1 |  | RIPK2 | 9 | 8 |
| EGFR<sup>T790M</sup> | 2 | 1 | ROR2 |  |  |
| EPHA2 | 4 | 1 | RPS6KA1 |  |  |
| ERBB3 | 6 | 1 | SPHK1 | 8 | 3 |
| ERBB4<sup>R711C</sup> | 4 | 2 | SRC |  |  |
| ERN1 | 1 | 1 | STK24 |  |  |
| FGFR1 | 8 |  | STK38 | 2 | 1 |
| FGFR2 | 6 | 3 | SYK |  |  |
| FGFR3 |  |  | TEC |  |  |
| FGR |  |  | TYK2 | 17 | 4 |
| FLT3 | 4 | 2 | ULK3 | 9 | 3 |
| FYN | 13 | 9 | YES1 |  |  |
| GSK3A | 1 | 1 | ZAK |  |  |
| HER2 | 32 | 16 | ZAP70 |  |  |

