## Supplementary information for "Inhibitor-induced supercharging of kinase turnover via endogenous proteolytic circuits"

#### Materials and Methods

##### Cell lines and cell culture

KBM7 cells (obtained from T. Brummelkamp) and KBM7 iCas9 cells (a gift from J. Zuber) were grown in IMDM (Thermo Fisher Scientific) supplemented with 10% heat-inactivated FBS (Sigma-Aldrich) and 1% penicillin/streptomycin (Pen/Strep, Gibco™). RKO iCas9-GFP and iCas9-BFP (gifted by J. Zuber), K562 (purchased from ATCC) and NALM-6 (obtained from A. Villunger) were cultured in RPMI 1640 (Thermo Fisher Scientific) supplemented with 10% FBS and 1% Pen/Strep. 293T lentiviral packaging cells (obtained from Clontech) and Flp-In™ T-REx™ 293 (obtained from Invitrogen™) were cultured in DMEM (Thermo Fisher Scientific) supplemented with 10% FBS and 1% Pen/Strep.

For competitive Kinobead pulldowns, Jurkat, MCF7, K-562, COLO-205 and MV-4-11 cells were cultured in RPMI 1640 medium (Biochrom GmbH) supplemented with 10% (v/v) FBS (Biochrom GmbH). SK-N-BE(2) cells were grown in DMEM/Ham’s F-12 (1:1) supplemented with 10% (v/v) FBS and OVCAR-8 cells were cultured in IMDM medium (Biochrom GmbH) supplemented with 10% (v/v) FBS.

Cell lines were cultured at 37 °C and 5% $CO_2$ in a humidified incubator and were regularly tested for mycoplasma contamination.

##### Plasmids and cloning

All plasmid preparation, unless specified otherwise, was performed in Stable Competent *E. coli* (NEB) or in case of destination vectors in One Shot™ ccdB Survival™ 2 T1R Competent Cells (Invitrogen) according to the manufacturers instructions.

The pLEX305-ccdB-Nluc-3xFLAG luminescent reporter vector was generated as a destination vector starting from pLEX_305-ccdB-dTAG destination vector (Addgene #91798) by restriction digest with AgeI and MluI and T4 DNA ligation (NEB) of a synthesized gene block (TWIST) containing the Nluc sequence and a c-terminal 3xFLAG tag (5’- *cgggcaaaaccggtgtcttcacactcgaagatttcgttggggactggcgacagacagccggctacaacc tggaccaagtccttgaacagggaggtgtgtccagtttgtttcagaatctcggggtgtccgtaactccgatccaaaggattgt cctgagcggtgaaaatgggctgaagatcgacatccatgtcatcatcccgtatgaaggtctgagcggcgaccaaatgggc cagatcgaaaaaatttttaaggtggtgtaccctgtggatgatcatcactttaaggtgatcctgcactatggcacactggtaa tcgacggggttacgccgaacatgatcgactatttcggacggccgtatgaaggcatcgccgtgttcgacggcaaaaagat cactgtaacagggaccctgtggaacggcaacaaaattatcgacgagcgcctgatcaaccccgacggctccctgctgtt ccgagtaaccatcaacggagtgaccggctggcggctgtgcgaacgcattctggcggactacaaggaccacgacggtg actacaaggaccacgacatcgactacaaggacgacgacgacaagtagtaaacgcgttgacgatgg* -3’).

To generate a destabilised version of the vector, an identical gene block was synthesised with the addition of the PEST sequence (Promega) 5’- *aattctcacggctttccgcctgaggttgaa gagcaagccgccggtacattgcctatgtcctgcgcacaagaaagcggtatggaccggcacccagccgcttgtgcttca gctcgatcaacgtc* -3’ upstream of the stop codon. pENTR223 Gateway® entry vectors for the kinases were obtained from Hahn/Root Labs Human Kinases ORF Kit[1,2] (Addgene Kit

#1000000014) with the exception of FYN and MAPK4, which were purchased seperately (BCCM, LMBP ORF81088-E05, LMBP ORF81100-B12). A pENTR221-GFP vector was generated via BP gateway cloning (Invitrogen™), starting from the PCR amplified GFP sequence of pCAG-GFP[3] (gifted by Connie Cepko, Addgene #11150) and insertion into the empty pDONR221 (Invitrogen™, #12536017). Final luminescent reporter vectors were generated by LR gateway cloning following the manufacturers recommendations (Invitrogen™, incubation was routinely run over night (o.n.) at 25 °C prior to heat inactivation and transformation). Correct insert size was assessed by analytical digest and in-frame cloning verified by sequencing (Microsynth, Austria).

Single point mutations with the exception of BLK S5A and S6A were generated from the respective pENTR223 plasmids using either the Q5® site-directed mutagenesis kit (see primers in **SI Table S1** "Method" SDM, NEB) or by Q5 (NEB) PCR amplification (see primers in **SI Table S1** "Method" PCR), followed by 1h of DpnI digest (NEB) and direct transformation into DH5α *E. coli* (NEB). Stability vectors were generated by digesting the previously published[4] plasmid backbone pRRL_SFFV_empty_BFP_P2A_mCherry with SalI and BamHI, prior to insertion of the PCR amplified kinase of interest using the NEBuilder® HiFi DNA Assembly Master mix (NEB) following the manufacturers instructions. Primers were designed with the NEBuilder® Assembly tool (see sequences in **SI Table S2**). BLK S5A and S6A stability reporter plasmids were generated analogously using the corresponding mutated primer pairs.

The pRRL_SFFV_CSK*_EF1a_iRFP670 was generated by restriction digest of SalI and XhoI of pRRL.SFFV.DACF16.EF1as.iRFP670[4] and insertion of the PCR amplified CSK fragment using the NEBuilder® HiFi DNA Assembly Master Mix (NEB). The gateway vectors pcDNA5_FRT_ccdB_3xFLAG_miniTurbo and pSTV6_ccdB_3xFLAG_miniTurbo (kindly provided by Anne-Claude Gingras), as well as, pRRL_EF1a_ccdB_emGFP_IRES_HygroR (gifted by Giulio Superti-Furga) formed the basis for the generation of the remaining kinase reporter vectors via LR gateway cloning (Invitrogen™). For the LYN BioID dataset pENTR223_LYN or pENTR221_GFP was cloned into pcDNA5_FRT_ccdB_3xFLAG _miniTurbo, and for the BLK BioID dataset pENTR223-BLK or pENTR223-APH1A (BCCM, LMBP ORF81047-H06) and pENTR221-GFP were cloned into pSTV6_ccdB_3xFLAG _miniTurbo vector.

sgRNAs were cloned into a single sgRNA vector pLenti-U6-IT-EF1a-Thy1.1-P2A-Neo or dual sgRNA vector pLentiDual-hU6-IT-mU6-IT-EF1a-Thy1.1-P2A-Neo (both kind gifts from Johannes Zuber) as previously reported[4]. The sgRNA sequences are described in **SI Table S3** and were designed using the VBC score[5].

All inserted DNA sequences were verified by Sanger Sequencing (Microsynth, Austria).

##### Cell line generation via lentiviral transduction

With exception of the generation of the LYN BioID cell lines (see **Generation of cell lines via flp recombinase**), all cell lines were generated via transduction of lentivirus. For virus production, 293T lentiviral packaging cells were transfected at 70 % confluence with the to be packaged plasmid in addition to the two packaging plasmids (pCMVR8.74 helper, pMD2.G envelope, both gifted by Didier Trono (Addgene #22036 and #12259) using polyethylenimine (PEI MAX® MW 40000, Polysciences). Viral supernatants were collected 60h post-transfection and cell debris removed using a 0.45 µm poly-ethersulfone filter.

For transduction, 1 million cells (K562 for luminescent reporters, KBM7, KBM7 iCas9 or RKO iCas9-GFP/BFP for all other reporters) per 2 mL were transduced with 250 µL virus solution and 8 µg/mL polybrene. If required, virus volume was adjusted to achieve the desired transduction efficiency. 24h post transduction, cells were expanded. For luminescent reporter cell lines, selection was performed with puromycin (1 µg/mL, Gibco) starting 48h after cell recovery. Subsequently cell pools were subjected to QC by means of immunoblot analysis using the c-terminal FLAG epitope tag, as well as assessment of luminescent levels using NanoGlo® Luciferase Assay System (Promega). For the latter, $10^5$ cells were seeded for each reporter cell pool in 30 µL on a 384 well plate and luminescence was measured on a Victor™ X3 2030 Multilabel Reader (Perkin Elmer).

The three cell lines generated for the BLK BioID experiment (performed in KBM7) were likewise selected with puromycin (1 µg/mL, Gibco). Generated cell pools were quality controlled after cell recovery for construct expression 24h post doxycycline treatment (1 µg/mL, PanReac AppliChem). Both correct fusion size and biotinylating efficiency were tested via immunoblot. The latter was conducted by additional incubation of cells with 100 µM biotin for varying timeframes and blotting for the biotinylated proteome using an Anti-Biotin antibody (see **Immunoblotting**).

sgRNA vector containing cells were selected using G418/Neomycin (1 mg/mL, Gibco) 72h after transduction. Completion of selection or sgRNA transduction efficiency was assessed via staining with APC anti-mouse Thy1.1 antibody (1:400, 202526, BioLegend) in Human TruStain FcX Fc Receptor Blocking Solution (1:1000, 422302, BioLegend) for 5 min at 4 ˚C, followed by 2 x PBS washing and subsequent analysis via flow cytometry. Genetic knockouts (k.o.) were generated by induction of the tightly inducible Cas9 cassette via doxycycline (0.4 µg/mL, PanReac AppliChem) for a timeframe of 48h up to 1 week (see **SI Table S3** for incubation times per sgRNA) prior to analysis via immunoblot, imaging or flow cytometry.

All fluorescent reporter cell lines were either used directly for flow cytometry or selected via FACS using a CytoFLEX SRT Benchtop Cell Sorter. In the first round of sorting, pools of reporter positive cells (see **SI Fig S3**) were enriched. For selected cell lines, single cells were sorted, expanded and utilized for flow cytometry, FACS-based CRISPR/Cas9 screens or imaging experiments. Specifically, the main stability reporters generated in KBM7 iCas9 for $LYN^{wt}$, $BLK^{wt}$ and $RIPK2^{wt}$ were used as clonal cell lines. In addition, the $BLK^{wt}$-GFP reporters in RKO iCas9-BFP were also used as clones. The remainder of the stability reporters generated for imaging purposes in RKO iCas9-GFP were used as sorted pools. The KBM7 iCas9 $LYN^{Y32A}$ stability reporter was used as a sorted cell pool, whereas the suite of KBM7 iCas9 BLK mutant stability reporters were used without sorting and instead analysis was performed on the reporter positive cell gate. For the latter, a matched unsorted $BLK^{wt}$ stability reporter was hence used as control in the corresponding datasets. All genetic knockouts were performed in a pooled format after G418 selection as detailed above.

##### Generation of cell lines via flp recombinase

Flp-In™ T-REx™ 293 cells were transfected with 200 ng of LYN-mT and GFP-mT plasmids and 2 µg pOG44 vector (Invitrogen™, V600520) using Lipofectamine 2000 (Invitrogen™) according to the manufacturers instructions in a 6-well format. 24h post-transfection, cells were expanded to a 10 cm dish and after an additional 24h, cell selection was iniated with 200 µg/mL hygromycin B (Roth). Cells were passaged and contained under hygromycin B selection for 4 weeks, prior to expansion and assessment of expression and biotinylation capacity analogue to described for the BioID cell lines generated via lentiviral transduction.

##### Temporal luminescent drug screen

Compounds from the kinase inhibitor library including 10 PROTAC controls (see **Compounds**) and respective transcription and translation compounds were dispensed via an Echo 550 system into white 1536-well plates (PerkinElmer, 6004684) at the appropriate concentrations (0.5-10 µM, see **Supplementary Data Table 1**; 10 µM for CHX and 1 µM for NVP-2). Plates were sealed and stored at -20 °C. On the day of the drug screen, plates were equilibrated to room temperature. Next, 5 µL of 1:100 Endurazine (Promega) in buffered RPMI (complete RPMI supplemented with 50 mM HEPES pH7, (Sigma-Aldrich, H0887)) were pre-laid into each well using a liquid dispenser (Thermo Fisher Scientific, Multidrop Combi). Then, 5 µL of cells at a density of 640.000 cells/mL in buffered RPMI were dispensed on top. Cells were transferred to an incubator (humidified chamber, 37 °C, 5% $CO_2$) and luminescent signal was measured every 4h from 2h to 18h post-seeding using an EnVision plate reader (Revvity). Raw luminescence signals were subsequently normalized for intra-plate effects. Finally, compound effects were quantified by calculating percentage of control (POC) based on averaged, outlier-corrected DMSO (100%) and positive control (CHIR-99021; 0%) wells, for each plate and timepoint individually (**SI Fig S1A**). Only compounds that passed an initial pre-selection step were used for the final drug screen. The pre-screen was performed following the identical steps but only for the two control cell lines GFP-Nluc and dGFP-Nluc and at two concentrations (2.5 µM or 10 µM for 10 mM stock compounds and 0.5 and 2 µM for 2 mM stock concentrations). Compounds were eliminated if any of the normalised POC data was smaller than 48 or larger than 150 or if the relative change to the 2h timepoint was bigger than 0.58. In cases were only the higher concentrations fulfilled these criteria, the corresponding lower concentration was used. In total thus 1620 compounds including 10 PROTACs were used for the drug screen (see **Supplementary Data Table 1**).

##### Data analysis of luminescent drug screen

Following the normalization of the initial drug screening data, we performed additional data processing in order to obtain a binary active/inactive classification for each compound-kinase pair.

First, we implemented a filter to exclude compound-kinase pairs exhibiting high variability across replicates (STDEV > 30, **Fig 1**) consistently across all five time points, resulting in the removal of 138 compound-kinase pairs. Overall, the proportion of pairs having 0 time points with high standard deviation was 99.67%. We further filtered out compounds that exhibited high reactivity against all kinases, considering them false positives due to their low initial 2h time point (35 compounds with a median POC across kinases < 70). We further excluded the three non-small molecules Disitertide (TFA), Pep2m myristoylated (TFA), pm26TGF-β1 (TFA) from our analysis. To ensure comparability of time series and to eliminate bias towards absolute POC values, we centered the compound-kinase series around 100 POC relative to the 2h time point. This centering process was first applied across kinases and then across compounds.

We employed the time series of CHX, NVP2 and DMSO controls to assess whether compound-kinase pairs significantly deviated from each control. For each kinase and time point, we independently calculated the normalized compound z-score. A compound was considered to significantly reduce the kinase readout if it exhibited a substantial reduction (2 sigma) compared to the null model of the controls. We utilized the same methodology to calculate z-scores of compounds concerning the distributions of all other compounds. We also calculated the z-scores normalizing against only the initial 2-hour time point to capture significant changes relative to the initial conditions.

This process resulted in eight normalization schemes: against CHX, NVP2, DMSO and compounds, considering both time point-independent and initial time point-dependent situations. Each normalization offered varying selectivity over the compound-kinase time series, and we then expressed the scores as the count of significantly decreased time points (2 sigma).

Finally, we conducted a parameter scan to define a query for selecting hit compounds by combining the scores and specifying the minimal number of significantly deviated time points for each normalization scheme and the overall total combined via ‘or’ operators. We determined the normalization score thresholds for the query by minimizing the false discovery rate. This was achieved using the 10 PROTAC controls and their respective kinase targets as a reference for true positives as well as a manually curated inclusion list. The query that reflected our constraints is the following:

[(DMSO_norm>=5) || (CHX_norm>=2) || (CPD_norm>=5) || (CHX_norm2h>=5)] || (TOT>=10)

Meaning that all the positive kinase-compound pairs have to globally score 10 or more, or having a normalized score above the determined threshold in at least one of the individual screens (**SI Fig S1B**). For the rare instance of a missing timepoint (mainly associated with the kinase reporters for CDK4, 7 and 9) the score was corrected by +1. One compound was excluded from further data analysis due to scoring in > 10 instances. The final KinDeg scores are shown in **Supplementary Data Table 1** (including the annotation of excluded compounds). The final hit kinase trajectories are shown in **Supplementary Document 1**.

The screening data was used to fit the half lifes of each kinase. This was perfromed by fitting the equation $100*e^{(-x*tau)}$ in Python (3.7.6) and the package scipy (1.4.1) to each kinase’s CHX screening trajectory. tSNE plots were generated with sklearn and matplotlib (1.0.1 and 3.5.3, respectively) from ChEMBL drug binding data processed as described in the Chemical Checker (CC)[6] and compounds were characterized with CC global bioactivity signatures. Chaperone client status was mapped from Taipale et al.[7] to the respective canonical kinases (see **Supplementary Data Table 1**). Jaccard distances (JD) between kinase hit profiles were calculated as 1 – Jaccard Similarities (JS), JS being the size of the intersection divided by the size of the union of two compound (hit) sets. Kinome trees were depicted using http://www.kinhub.org/kinmap/index.html.

#### Immunoblotting

Cell pellets (1-2 million cells per treatment) were lysed in UREA lysis buffer (8 M Urea, 1% CHAPS, 50 mM Tris-HCL pH 8) for 30 min shaking at 4 ˚C and 1200 rpm. Next samples were cleared by centrifugation for 15 min (20000 x g, 4 ˚C) and quantified using the Pierce BCA Protein Assay Kit (Thermo Fisher Scientific) according to the manufacturer's instructions. Finally, samples were diluted with Bolt LDS sample buffer (4x) (Invitrogen) supplemented with f.c. 10% beta-mercaptoethanol (Sigma-Aldrich) and denatured for 10 min at 70 ˚C. 20 µg per protein sample were separated on a Bolt 4-12 % Bis-Tris Plus Gel (10-17 wells) (Invitrogen) using the Color Prestained Protein Standard, Broad Range (10-250 kDa, NEB) as a marker. After transfer to a nitrocellulose membrane, membranes were stained by Ponceau-S. Next, the membranes were blocked with 5% milk in TBS-T (30 min, RT) and then incubated with primary antibodies *o.n.* at 4°C in TBS-T. The following day, membranes were washed thrice with TBS-T followed by 1h RT incubation with the respective secondary antibodies if required. Finally, membranes were again washed thrice before analysis on a Chemidoc and using Pierce ECL Western Blotting Substrate (Thermo Fisher Scientific). Following antibodies and dilutions were used: GAPDH (1:5000; Santa Cruz Biotechnology, sc-365062), GAPDH (1:5000; Santa Cruz Biotechnology, sc-47724), Vinculin (1:500; Szabo Scandic, SACSC-25336), FLAG (1:2000; Cell Signalling Technology, #2368), LYN (1:1000; Cell Signalling Technology, #2796), BLK (1:1000; Cell Signalling Technology, #3262), RIPK2 (1:1000; Cell Signalling Technology, 4142S), Phospho-Lyn (Tyr507) (1:1000; Cell Signalling Technology, #2731), FIP200 (1:1000; Cell Signalling Technology, #12436), CDK9 (1:1000; Cell Signalling Technology, #2316), HRP-conjugated Anti-Biotin (1:1000; Cell Signalling, #7075), Peroxidase-conjugated Goat Anti-Rabbit IgG (1:10000; Jackson ImmunoResearch 111-035-003), Peroxidase-conjugated Goat Anti-Mouse IgG (1:5000; Jackson ImmunoResearch JAC115035003). For quantifications the accompanying ChemiDoc ImageLab™ software was used, normalized against the respective loading control and plotted as fold changes with respect to each genotype's DMSO control or 0h timepoint. The data was plotted as mean from three independent biological replicates ± SD. Replicates and uncropped images are shown in **SI Fig S7-18**.

#### Compounds

Carfilzomib (Cay17554-5) and Bafilomycin A1 (Cay11038) were purchased from Cayman, HSP90i (4-(4-(23-Dihydro-14-benzodioxin-6-yl)-5-methyl-1H-pyrazol-3-yl)-6-ethylresorcinol), 385920) was obtained from Calbiochem®. All other small molecule inhibitors were sourced from MedChemExpress. These include the Kinase inhibitor library (see **Supplementary Data Table 1**, 1996 compounds, HY-L009), MLN4924 (HY-70062), TAK-243 (HY-100487), TAK-285 (TAK285, HY-15196), Src inhibitor 3 (SI-3, HY-130254), RIPK-IN-4 (RI-4, HY-107978), AV-412 (HY-10346), Neratinib (HY-32721), Afatinib (HY-10261), WZ4002 (HY-12026), Nintedanib (HY-50904), DAPT (HY-13027), alkynyl myristic acid (HY-140335), THAL-SNS-032 (dCDK9, HY-123937) and NVP-2 (HY-12214A). Cycloheximide (CHX) was purchased from Cell Signaling Technology (2112S).

All compounds were dissolved in DMSO (Sigma-Aldrich, D1435) as 1 mM, 10 mM, 20 mM or 100 mM stock solutions. Working dilutions were prepared as 1000 x or 2000 x stock solutions. The kinase inhibitor library was delivered as 2 mM or 10 mM stock solutions (see **Supplementary Data Table 1**).

##### Flow cytometry

Cells were treated with the compound, concentrations and timeframes indicated in the respective figure legends and the fluorescent channels of interest were subsequently analyzed on a LSR Fortessa (BD Biosciences) with the BD FACSDiva software (v9.0). The data was analyzed with FlowJo (10.6.2) as outlined in **SI Fig S3** and the resulting mean BFP and mCherry values were exported for further processing. BFP/mCherry ratios were calculated after background subtraction (from matched *wt* cells) and normalized to either each pre-treatment or genetic variant (referred to as norm. BFP in the figure legends) or normalized to a specific condition as indicated in the respective subscripts e.g. DMSO in **Fig 3E**. Decay functions were fitted using Y=(Y0 - Plateau)*e(-K*X) + Plateau and dose responses fitted using Y = Bottom + (Top - Bottom)/(1+(IC50/X)n) where n = the Hill slope using the in-built functions of GraphPad Prism (10.0.3) and nonlinear regression fitting. Matched mCherry flow histograms to **Fig 2A, 4A** and **Extended Data Fig 4C** are shown in **SI Fig S4A**, **S5A** and **S6A**, respectively.

##### FACS-based CRISPR/Cas9 screen

The screens were performed as previously described[4]. First, cells were transduced at an MOI of 0.1-0.2 with lentivirus harboring the respective sgRNA library, prepared as described in **Cell line generation via lentiviral transduction** to achieve a 1000 x representation per sgRNA. For LYN the previously published UPS-focused sgRNA library[8] (7801 sgRNAs) and for BLK and RIPK2 a genome-wide library[5,9] was used. 72h after transduction, the transduction rate was assessed by staining with APC anti-mouse Thy1.1 antibody (1:400, 202526, BioLegend) and Human TruStain FcX Fc Receptor Blocking Solution (1:1000, 422302, BioLegend) for 5 min at 4 °C. Next, selection with G418 (1 mg/mL, Gibco) was initiated. Cells were maintained in G418 positive media for at least 14 days, splitting cells every 48-72h. For the screen, Cas9 expression was induced with doxycycline (0.4 µg/mL, PanReac AppliChem) and after 72h cells were treated with DMSO or the respective inhibitors (SI-3 156 nM, 8h; TAK285, 6h; RI-4 2.5 µM, 18h). Cells were centrifuged for 5 min at 500 x g and stained with APC anti-mouse Thy1.1 antibody (1:400, 202526, BioLegend), Zombie NIR Fixable Viability Dye (1:1000, BioLegend) and Human TruStain FcX Fc Receptor Blocking Solution (1:1000, 422302, BioLegend) for 5 min at 4 °C. Subsequently cells were fixed with BD Fixation buffer 4 % (Thermo Fisher Scientific™ Pierce™) for 45 min at 4 °C followed by two washes with PBS and resuspension in FACS buffer (PBS, 5 % FBS and 1 mM EDTA) for storage at 4 °C. All staining steps were performed in the dark. Cells were sorted within 48h of fixation.

Sorting was performed on a BD FACSAria Fusion (70 µm nozzle, BD Biosciences, BD FACSDiva software (v8.0.2)). First, cells were strained through a 35-µm nylon mesh. Next, cells were sorted for the 5% highest and lowest BFP expressing cells as well as 30% of the mid-fraction (see gating strategy in **SI Fig S3**). For each replicate and condition, cells corresponding to at least a 500-fold (genome-wide), or 1000-fold (UPS-focused) library representation were sorted.

After sorting, high, low and mid-fractions were pooled per replicate and lysed *o.n.* (14h) at 55 ˚C shaking at 1200 rpm in lysis buffer (10 mM Tris-HCl, 150 mM NaCl, 10 mM EDTA, 0.1 % SDS) supplemented with proteinase K (New England Biolabs). The following day, RNAse was removed with DNAse-free RNAse (Thermo Fisher Scientific) for 2h at 37 ˚C. Lysates were stored at -20 ˚C until further processing.

For DNA extraction, two rounds of phenol extraction (UltraPure™ Buffer-Saturated Phenol, Thermo Fisher Scientific, 15513039) using phase Lock Gel tubes (VWR, 7332477) followed by isopropanol precipitation *o.n.* at -20 ˚C were performed. Next, samples were barcoded using a two-step PCR protocol (AmpliTaq Gold polymerase, Invitrogen, 4311818). After each PCR step, amplicons were cleaned up with Mag-Bind TotalPure NGS beads (Omega Biotek) using the manufacturers protocol for double-sided selection. Final NGS libraries were pooled at equimolar ratios and sequenced on a HiSeq 3000 or NovaSeq 6000 platform (Illumina).

The resulting reads were trimmed using fastx-toolkit (v0.0.14) and subsequently aligned (Bowtie2 v2.4.5) and quantified (featureCounts v2.0.1). The corresponding workflows are available at https://github.com/ZuberLab/crispr-process-nf/tree/566f6d46bbcc2a3f49f51bbc96b9820f408ec4a3 and https://github.com/ZuberLab/crispr-mageck-nf/tree/c75a90f670698bfa78bfd8be-786d6e5d6d4fc455. Gene-level enrichment was calculated by comparing each high or low population to the corresponding mid population using the median-normalized read counts. The resulting $log_2FC$ and p-values as well as the number of scoring and total quantified sgRNAs per gene can be found in **Supplementary Data Table 3**.

##### High-content confocal imaging and data analysis

Cells were imaged using a PerkinElmer Opera Phenix automated microscope run on the Harmony® software (4.9 or later versions) and utilising the pre-set filter settings for DAPI (BFP), AF-488 (GFP), mCherry and brightfield. Exposure was set to < 400 ms per channel, and BFP and GFP channels were seperated during acquisition if required. Cells were seeded 24h prior to imaging into 384 or 96-well (CellCarrier Ultra, Revvity) to achieve a final cell density of 40-60%. Drugs were added imediately prior to imaging as indicated in the figure legends.

Cells were segmented using cellpose[10] (0.6.5-foss-2020b) using either the mCherry (RIPK2) or GFP (BLK) channel and an adjusted diameter of 38 or 50, respectively. Next, relevant features and fluorescence were extracted using custom-built cellprofiler pipelines (4.1.3-foss-2020b).

In all instances, “ConvertImageToObjects” (convert to boolean image (no), preserve original labels (yes)) was used to generate the primary objects. Next, for RIPK2, “EnhanceOrSuppressFeatures” was applied (Operation = Enhance, Type = Speckles, Size 6, Speed and accuracy = Fast) followed by “IdentifyPrimaryObjects” (diameter = 2-20, thresholding strategy = global, method = manual, threshold = 0.0016, smoothing scale 1.3488, method clumped objects&draw lines between clumped objects = Intensity, automatic smoothing and distance calculation enabled, holes filled in after both thresholding and declumping). “RelateObjects” was applied to assign the resulting speckles per cell object. Finally, “MeasureObjectIntensity” and “MeasureObjectSizeShape” were applied for measuring the respective parameters across the speckles and cell objects, before exporting the data to a database for further processing via self-written python scripts. For the RIPK2 data associated with panels **Fig 4H-J** and **Extended Data Fig 5F**, due to the different absolute BFP fluorescence values of the constructs for RIPK2$^{wt}$ and RIPK2$^{\Delta CARD}$, two steps were added prior to “EnhanceOrSuppressFeatures”. Namely, “ExpandOrShrinkObjects” was applied to eliminate cell boundaries (Operation = shrink by a specified number of pixels, pixels = 4) followed by “ImageMath”, which was used to calculate the BFP to mCherry ratio. The thresholds in “IdentifyPrimaryObjects” were thus adapted to 0.2 instead of 0.0016.

For the BLK-GFP cell clones, only module “MeasureObjectIntensity” was applied after object classification. Corresponding data was exported to a spreadsheet for further processing.

In all instances, Python (3.7.6) was used to annotate the resulting data (condition, replicate) and normalise the data. Normalised data was then exported and depicted in GraphPad Prism (10.0.3). In all cases, data was averaged per biological replicate of the mean values per cell. SD was correspondingly calculated across the biological replicates.

#### NanoBRET

The assay was performed as described previously[11]. In brief, full-length CSK and LYN were obtained as plasmids cloned in frame with an N-terminal Nluc-fusion (kind gift from Promega). Plasmids were transfected into HEK293T cells using FuGENE HD (Promega, E2312), and proteins were allowed to express for 20h. Serially diluted inhibitor and NanoBRET K4 Tracer (Promega, TracerDB ID: T000037) at the Tracer KD concentration taken from TracerDB[12] were pipetted into white 384-well plates (Greiner 781207) using an ECHO acoustic dispenser (Labcyte). The transfected cells were added and reseeded at a density of $2 \times 10^5$ cells/mL after trypsinization and resuspending in Opti-MEM without phenol red (Life Technologies). The system was allowed to equilibrate for 3h (37 °C, 5% $CO_2$) prior to bioluminescence resonance energy transfer (BRET) measurements. To measure BRET, NanoBRET NanoGlo Substrate and extracellular Nluc Inhibitor (Promega, N2540) was added as per the manufacturer’s protocol, and filtered luminescence was measured on a PHERAstar plate reader (BMG Labtech) equipped with a luminescence filter pair (450 nm BP filter (donor) and 610 nm LP filter (acceptor)). Competitive displacement data were then analyzed using GraphPad Prism (10.0.3) software using a normalized 3-parameter curve fit with the following equation: Y = 100/(1 + 10(X − log IC50)).

#### Commercial recombinant binding/inhibitory assays

*In vitro* kinase inhibitory or kinase binding assays were performed using the SelectScreen platform (Thermo Fisher Scientific). TAK285 (BLK) and Src inhibitor 3 (SI-3, LYN) were screened using the Z'-LYTE assay, while RIPK-IN-4 (RI-4, RIPK2) was screened with the LanthaScreen Eu Kinase Binding Assay according to their respective assay availability. 3-

fold dilutions were performed starting from 30 µM and in presence of ATP, using its standard apparent $K_M$ per kinase.

#### Immunoprecipitation

Cell pellets (10 million cells) were lysed in 900 µL IP lysis buffer (50 mM Tris-HCL (ph7.4), 150 mM sodium chloride, 0.1% TritonX-100, 1 mM EDTA and 5 mM magnesium chloride, 1x protease inhibitors) followed by lysate clearance, protein quantification and immonoprecipitation as described in **Immunoprecipitation and on-bead TAMRA click**. After immunoprecipitation and sample washing, proteins were then directly eluted using 70 µL as final volume prior to analysis via immunoblot. For blocking 5% BSA in TBS-T was used instead of 5% milk in TBS-T and the phospho LYN Y507 antibody was diluted 1:1000 in TBS-T containting 3% BSA and 0.1% sodium azide.

#### Immunoprecipitation, on-bead TAMRA click and in-gel fluorescence

Cell pellets (15 million cells per condition) were lysed in 900 µL NP40 lysis buffer (DPBS with 1.5 mM magnesium chloride, 1% NP40, 1x protease inhibitors, 1 x benzonase) for 30 min on ice. Lysates were cleared by centrifugation (20 min, 4 ˚C, 20000 x g) and quantified using the Pierce BCA Protein Assay Kit (Thermo Fisher Scientific) according to the manufacturer's instructions. Next, samples were normalized to 1 mg per input and pre-activated anti-Flag magnetic beads (Sigma-Aldrich) were added, followed by the incubation for 3h at 4˚C on a rotating wheel. Beads were washed thrice with lysis buffer. After removal of the supernatant, 56 µL of click-mix (170 µM TAMRA (5-TAMRA-Azide CLK-FA008 Jena Biosciences), 230 µM copper sulfate, THPTA 1.15 mM, HCl 5 mM, sodium ascorbate 5 mM, in PBS) are added per sample. Finally, 18 µL of elution buffer (4x Laemmli buffer supplemented with f.c. 10% beta-mercaptoethanol) are added and the samples boiled at 95˚C for 10 min prior to loading 20 µL of supernatant and analysis via SDS-page. Prior to the transfer for immunoblotting and continuation as described in section **Immunoblotting**, the SDS-page was imaged on a ChemiDoc using the Alexa 546 channel and Coomassie for the ladder.

#### Data plotting and statistical analysis

All data is represented as the mean of technical or biological replicates ± SD (standard deviation) or ± CI (confidence interval). Data points were calculated as described in the respective sections.

Imaging data, all data related to the drug screen, proteomics, CRISPR screen, as well as *in vitro* kinase binding/inhibitory assay were plotted with seaborn (0.12.2) and matplotlib (3.4.2) in Python (Python 3.7.6). Standard packages such as numpy (1.21.5), pandas (1.0.1) and scipy (1.4.1) were correspondingly used for data handling, processing, normalization, statistical calculations and/or data fitting. Immunoblot quantifications and data associated with flow cytometry (except for flow histograms) were plotted in GraphPad Prism (10.0.3). Statistical tests and data fitting for the corresponding datasets were calculated directly with in-build functions as detailed in the respective sections.

Flow histograms were exported from FlowJo (10.6.2). Representative images of microscopy experiments were prepared using Fiji (ImageJ, 2.1.1/1.53i).

**Preparation of BioID MS-samples**

Bait-mT expression was induced 24h prior to iniating cell treatments using 1 µg/mL doxycycline. The following day, the respective inhibitors (TAK285, 2.5 µM or SI-3, 156 nM) or vehicle control (DMSO, across all cell lines including the two GFP-mT versions) and 100 µM Biotin (Sigma-Aldrich, B4501) were added to the cells for 1h. For BLK-mT an additional condition including 2h Carfilzomib (1 µM) pre-treatment prior to TAK-285 addition was generated. After the treatments, 20 million cells per condition and replicate were harvested by centrifugation followed by two ice-cold PBS washes. Resulting cell pellets were snap frozen on dry ice and stored at -80 ˚C until further processing.

The following protocol has been adapted from Offensperger *et al*[13]. All steps were carried out with Protein LoBind tubes (Eppendorf) and HPLC-grade reagents. Briefly, for lysis cell pellets were resuspended in 250 µL lysis buffer (PBS supplemented with 1% SDS (Sigma-Aldrich, 71736), 2 mM magnesium chloride (Invitrogen, AM9530G), protease inhibitors (Thermo Fisher Scientific, 78437) and benzonase (Merck, US170746-3)). Samples were vortexed and incubated at 37 ˚C (300 rpm, 30 min) followed by centrifugation at 18000 x g (4 ˚C, 30 min). Supernatant was transferred to fresh tubes and protein concentration was measured using Pierce™ 660nm Protein Assay Reagent (Thermo Fisher Scientific, 22660) following the manufactures instructions. Per sample 1 mg of total protein were diluted up to a final volume of 300 µL with lysis buffer. Next, 30 µL of 50 mM TCEP (Sigma-Aldrich, 75259, diluted in $H_2O$) was added, samples vortexed and incubated for 1h at 56 ˚C shaking at 300 rpm. Then 80 µL of 1M HEPES (pH 7.5, AppliChem, A6916) were added, followed by 45 µL of 200 mM iodoacetamide (Sigma-Aldrich, I1149). Samples were again vortexed and incubated at 25 ˚C and 300 rpm. Pierce™ Streptavidin Agarose (Thermo Fisher Scientific, 20353) resin was prepared by centrifugation for 30 s followed by two PBS washes. Next the protein samples were added and incubated on a rotator for 1h at RT in the dark. Finally, samples were washed twice utilizing 1x pre-washed Mini BioSpin columns (Bio-Rad, 7326207) with wash buffer 1 (0.2% SDS in 1x PBS), followed by 16 washes with wash buffer 2 (8 M urea in 1x PBS) and four washes with PBS. For elution, slurry was resuspended in 2x digestion buffer (50 mM ammonium bicarbonate, 200 mM guanidine hydrochloride, 1 mM calcium chloride, in $H_2O$) and transferred again to a fresh tube. Subsequently, supernatant was removed and 250 µL digestion buffer, as well as, freshly supplied 10 µL of trypsin solution (0.1 µg/µL, Promega, V5117) added, prior to incubation *o.n.* on a rotating wheel (14h).

The following day, beads were centrifuged briefly (30 s) and supernatant transferred into a fresh tube. Resin was washed 1x using 200 µL $H_2O$, which was added to the already separated supernatant. Peptides were cleaned up with self-made stage tips columns. These were prepared from 1 mm circles of an Epore C18 disk inserted into a 200 µL tip. On top of the C18 disk, 24 µL Oligo R3 solution (15 mg/mL in acetonitrile (ACN)) were added prior to 1 min of centrifugation (1000 x g). The column was then washed twice with 100 µL ACN (1000 x g for 1 min) and equilibrated twice with 200 µL 0.1% TFA (3 min at 1000 x g). Samples were acidified with 30% TFA (1% final concentration) prior to loading of the samples in two fractions onto the column (1000 x g for 3 min). One wash with

200 µL 0.1% TFA (3 min at 1000 x g) was followed by a double elution step using 50 µL elution buffer (90% ACN, 0.01% TFA, in $H_2O$) each. Eluted peptides were dried using a vacuum centrifuge (45 ˚C) and stored at -20 ˚C. Next samples were TMT-labelled with the TMTpro 18-plex Label Reagent Set (Thermo Fisher Scientific, A52045) following the manufacturer's instructions. Subsequently, labelled peptides were pooled and fractionated using on-tip high pH fractionation. 1 mL of 20 mM ammonium formate (pH 10) was added per 320 µL of pooled sample and added again to self-made C18 columns, prepared as stated above except for the final wash steps, which were performed with 200 µL of 20 mM ammonium formate pH 10 instead of ACN. Samples were loaded in fractions of 250 µL followed by a wash with 200 µL of 20 mM ammonium formate pH 10. Centrifugation at each step was carried out for 3 min and 1000 x g. Elution was carried out in five fractions (2 min at 1000 x g) with buffers containing 20 mM ammonium formate (pH10) and different percentages of ACN (16%, 20%, 24%, 28%, 80%). First 50 µL were used per respective buffer, followed by 20 µL per buffer (2 min 1000 x g each). Next, all fractions were dried using a vacuum centrifuge at 45 ˚C and the resulting dried peptides were stored at -20 ˚C until data acquisition.

**Sample preparation for full proteome profiling**

Per condition 20 million cells were lysed in 300 µL lysis buffer (50 mM HEPES, pH 8 supplemented with 1 mM PMSF, protease inhibitor cocktail (Sigma-Aldrich) and 2% SDS). Cells were homogenized by pipetting and incubated at room temperature (RT) for 20 min. Next samples were sonicated (Covaris S2 high-performance ultrasonicator) for 150 s. Lysates were clarified by centrifugation at 20000 x g for 5 min at RT. Extracted protein amounts were determined by BCA (Pierce BCA Protein Assay, # 23227). For each sample 200 µg protein was digested using a filter-aided sample preparation (FASP) protocol essentially according to published procedures[14].

In short, proteins were reduced by addition of DTT (final concentration 83.3 mM), followed by incubation at 95 ˚C for 5 min. After cooling samples to RT, samples were mixed with 200 µL freshly prepared 8 M urea in 100 mM Tris-HCl at pH 8.5 (UA-buffer) and added onto FASP filter units (Merck Millipore). For buffer exchange, samples were centrifuged at 14000 x g for 15 min at 20 ˚C and residual SDS was washed by an additional washing step with 200 µL UA-buffer. All subsequent centrifugation steps were done at 14000 x g for 15 min at 20 ˚C. Proteins were alkylated by addition of iodoacetamide (50 mM final concentration) and incubated for 30 min at RT in the dark. Samples were washed three times with 100 µL UA-buffer followed by three washes with 100 µL TEAB buffer (Sigma-Aldrich). Proteins were digested by addition of sequencing-grade trypsin at a ratio of 1:50 at 37 ˚C *o.n.*.

To collect peptides, 50 µL of 50 mM TEAB buffer was added and samples were centrifuged. Filters were additionally washed with 50 µL of 0.5 M NaCl and the flowthroughs of both washing steps were pooled. Peptides were cleaned-up by C18 with peptide desalting spin-columns (Thermo Fisher Scientific). The peptides of each condition were labelled with TMTpro 18plex reagents according to the manufacturer (Thermo Fisher Scientific). After 1h of labelling, 1 µL of each channel was pooled together, quenched and cleaned-up by C18 and concentrated under reduced pressure. This test-mix was measured by DDA employing an OT/OT and quantification on MS2 level. The test-mix was used to calculate the median signal intensity of each TMTpro channel. The ratios to the lowest median channel intensity were derived and all channels were normalized to equalize labelling efficiency. The pooled channels were quenched, and samples were cleaned up by C18. As additional QC of channel normalization another test-pool was injected. After pooling all samples, an aliquot of 100 µL corresponding to roughly 450 µg was cleaned-up by C18 and resuspended in 10 mM ammonium formate buffer pH 10. Peptides were separated on an C18 reversed phase column (150 x 2.0 mm Gemini-NX, 3 µm C18 110Å, Phenomenex) by liquid chromatography into 96 time-window based fractions operating at 50 µL/min constant flow rate. A total of 36 fractions were collected, employing a previously described pooling strategy[15]. Samples were fractionated into glass-vials with 5 µL 30% TFA to acidify samples after fractionation. Fractions were dried under reduced pressure and reconstituted in 0.1% TFA for MS analysis. Additional information with regards to the reagents can be found in **SI Table S4**.

##### LC-MS/MS data acquisition of BioID and full proteome samples

Mass spectrometry data were acquired on an Orbitrap Fusion Lumos Tribrid mass spectrometer (Thermo Fisher Scientific) coupled to a Dionex Ultimate 3000 RSLCnano system (Thermo Fisher Scientific) interfaced with a Nanospray Flex Ion Source (Thermo Fisher Scientific). Peptides were loaded on a trap column (PepMap 100 C18, 5 µm, 5 × 0.3 mm, ThermoFisher Scientific) at a constant flow rate of 10 µL/min with 0.1% TFA in HPLC-grade H2O.

Next, the trap column was switched in-line, and peptides were separated on an analytical column (50 cm, 75 mm inner diameter) in-house packed with ReproSil-Pur 120 C18-AQ, 3 mm (Dr. Maisch HPLC GmbH) fitted to an ESI emitter fused silica (20 µm ID x 7 cm L x 365 µm OD; Orifice ID: 10 µm, CoAnn Technologies) kept at 50˚C. For the analysis, an analytical gradient of 190 min operated at a constant flow rate of 230 nL/min was used. The HPLC was operated with buffer A (0.4% formic acid in HPLC-grade $H_2O$), and buffer B (0.4% formic acid in ACN).

The analytical gradient comprised of the following steps: 0 to 4 min constant 6% buffer B, 4 -5 min from 6 to 9%, followed by an increase to 30% buffer B from 5 to 146 min, followed by an increase to 65% buffer B from 146 to 154 min, and a flush at 100% buffer B. The column was re-equilibrated at 6% buffer B from 167 to 190 min. Samples were acquired in data-dependent acquisition (DDA) mode using a maximum of 10 dependent scans (TopN approach) with synchronous precursor selection (SPS) enabled. Peptides were ionized by applying a constant voltage of 1.8 kV. MS1 precursor survey scans for MS2 and MS3 levels were acquired with scan range of 400 - 1600 m/z and a resolution of 120000 (at 200 m/z) in the Orbitrap. The automatic gain control (AGC) was set to 'standard' with amaximum injection time 50 ms. Precursor ions were filtered by charge state (2-5) excluding undetermined charge states with a dynamic exclusion (60 s with a ±10 ppm window), and monoisotopic precursor selection.The MS1 precursor intensity threshold was set to 5.0e3. For MSn data analysis, a charge-state filter was used to select precursors for data-dependent scanning. In MS2 analysis, spectra were obtained using one charge state per branch (from z=2 to z=5) in a dual-pressure linear ion trap (ITMS2). Ions were isolated employing a quadrupole isolation window with an isolation window of

+/- 0.7. Fragmentation was achieved by collision-induced dissociation (CID) with a fixed normalized collision energy of 35% and an CID activation time of 10 ms. For MS2 scans, the normalized AGC target was set to 200% with a maximum injection time of 35 ms. For MS3 scans, precursor ions were isolated using SPS waveform with varying isolation windows for charge stats: 1.3 m/z for z=2, 1.2 m/z for z=3, 0.8 m/z for z= 4 and 0.7 m/z for z = 5. Fragment ions were further fragmented by high-energy collision induced dissociation (HCD) at a fixed activation energy at 45% collision energy. The AGC target was set to 300% with a maximum injection time of 100 ms. Orbitrap scan range was set to 100 - 500 m/z at a resolution of 50000. Xcalibur Version 4.3.73.11 and Tune 3.4.3072.18 were used to operate the instrument.

**Processing of BioID raw MS-injections**

MS-raw files were processed with the Proteome Discoverer software (PD, Thermo Fisher Scientific, version 2.4.1.15). For the LYN dataset, a subset of 9 TMT-channels (126, 127N, 127C, 132C, 133N, 133C, 134N, 134C, 135N) were used, whereas for the BLK and APH1A experiment the full channel set was processed. The two BioID datasets were independently processed.

The peptide identification search was performed with Sequest HT, searching for fully tryptic peptides with maximal 2 missed cleavages and a minimum peptide length of 6 and a maximum of 144 amino acids. The precursor mass tolerance was set to 10 ppm and fragment ion mass tolerance was restricted to 0.6 Da. Spectra were searched against the canonical human protein database obtained from UniProtKB (download 05.11.2021, 20,304 sequences) appended with an in-house generated list of common lab contaminants (298 sequences) and streptavidin. As variable modification Methionine oxidation (+15.994 Da), Deamidation (0.984 Da), phosphorylation on Serine, Threonine and Tyrosine (+79.966 Da) and N-terminal specific acetylation (+42.011 Da), methionine loss (-131.040 Da) and acetylation with methionine loss (-89.030 Da) with a maximum number of three variable modification of the same type per peptide. Carbamidomethylation (+57.021 Da) of cysteine residues and tandem mass tag (TMT) 18-plex labeling of peptide N termini and lysine residues (+304.207 Da) were used as static modification. PSM and peptide FDR were controlled by Percolator at 1% respectively. Obtained results were filtered to include only spectrum matches with a Sequest HT cross-correlation factor (Xcorr) larger or equal to 0.9. Phopshosites needed a minimum site-probability of 75 corresponding to the high confidence threshold. For protein abundance inference, only high-confidence proteotypic peptides were included.

Protein and peptide intensities were derived from TMTpro reporter ion intensities. The reporter abundances were based on signal-to-noise (S/N) values if applicable, else reporter ion intensities were used. Correction of isotopic impurities was enabled. A co-isolation threshold for isolation interference of precursors was set to 80%. Further, to remove noisy signals, an average TMTpro reporter ion S/N threshold smaller or equal to 10 was used with an addtional SPS mass matches threshold of 65%, removing peptides with strong interferences. The obtained data was normalized using the sum total peptide amount and scaled to all average. For normalization and to derive protein abundances all quantified peptides were used. Protein ratios and $\log_2$FC were directly calculated from the grouped protein abundances, without missing value imputation. Abundance changes were tested for their significance, by an ANOVA on individual proteins across biological triplicates. P-values were corrected for multiple-testing with Benjamini-Hochberg.

##### Data analysis and representation of BioID data

The protein level PD output was used for further analysis. For the LYN BioID experiments, 4325 UniProtKB accessions were identified and for BLK, APH1A BioID experiments 2962 UniProtKB accessions were found. From the datasets proteins flagged as contaminates and proteins without quantification values were removed, resulting in 3564 UniProtKB accessions for the LYN dataset and 1870 UniProtKB accessions for the BLK, APH1A dataset. For subsequent data analysis, the PD derived normalized intensities, $log_2FC$ and the adjusted p-value (BH) were used. For the LYN experiment the analysis focused on significantly changed proteins upon SI-3 treatment versus vehicle/baseline control (DMSO). To identify enriched proximity interactions, a combined threshold of a $log_2FC \geq 2$ and an adjusted p-value $\leq 0.01$ against GFP controls were employed. The same thresholds were used to identify differentially changed interactions in the LYN SI-3-treated samples against LYN DMSO treated control. For the BLK/APH1A dataset additional scoring of proximity interaction partners was performed employing SAINTq[16]. For this, the total sum normalized protein intensities per replicate and condition were grouped together and scored against GFP negative controls (DMSO). SAINTq was performed on protein level (parameters: normalize_control = false, input_level = protein, compress_n_ctrl = 3, and max score across bait replicates). Proteins with a $log_2FC \geq 0.5$, a SAINTq-score ≥ 0.99 and a BFDR ≤ 0.01 were considered as high-confidence proximity interaction partners. As further filter, the CRAPome[17] frequency was mapped to each prey protein, using a 10% frequency threshold. Bait proteins were excluded from the CRAPome filter. Further, prey proteins which were annotated as kinases and type I transmembrane proteins in UniProtKB were filtered. These annotated interactors were further filtered for significant changes against GFP negative controls, employing the adjusted p-value (BH) from the ANOVA hypothesis test performed within PD. The obtained proximity interactors were intersected between BLK and APH1A, revealing on the one hand bait specific and on the other hand shared preys. Data analysis and visualizations were generated employing the statistical software R (version 4.3.1). The resulting processed datasets can be found in **Supplementary Data Table 4**. Normalisation results and additional individual vulcano plots or scatter plots of SAINTq results can be found in **SI Fig S4D-E** for LYN and **SI Fig S5 B-D** for BLK/APH1A.

##### Processing and data analysis of full proteome profiling data

The full proteome data was processed in Proteome Discoverer v.2.4.1.15 deriving protein intensities using the TMTpro 18 report ion quantities.

Peptide identification search was performed with Sequest HT searching for fully tryptic peptides of a minimum of 6 to up to 144 amino acids length and allowing for max. 2 missed cleavage-sites. Precursor mass tolerance was to 10 ppm and fragment ion mass tolerance was restricted to 0.6 Da. The search was performed against the canonical human protein database obtained from UniProtKB (download 12.11.2020) appended with an in-house generated list of common lab contaminants and streptavidin.

As variable modification methionine oxidation (+15.994 Da) and N-terminal specific acetylation (+42.011 Da), methionine loss (-131.040 Da) and acetylation with methionine loss (-89.030 Da) were set. The maximum number variable modification of the same type was limited to 3. As static modification carbamidomethylation (+57.021 Da) of cysteine residues and tandem mass tag (TMT) 18-plex labeling of peptide N termini and lysine residues (+304.207 Da) were set. PSM and peptide FDR were controlled with Percolator at 1% respectively. Obtained results were filtered to include spectrum matches with a Sequest HT cross-correlation factor (Xcorr) ≥ 1 and strict Percolator target FDR filters. Fur further analysis only peptides scored with high confidence and proteins at least identified with 1 proteotypic peptide were used. Protein and peptide intensities were derived from TMTpro reporter ion intensities. The reporter abundances were based on signal-to-noise (S/N) values if applicable, else report ion intensities were used. Correction of isotopic impurities was enabled. Co-isolation threshold of 70% for isolation interference of precursors was used. Further, to remove noisy signals, an average TMTpro reporter ion S/N threshold ≤ 10 was used. A SPS mass match threshold of at least 65% was applied. For report ion-based quantification unique and razored peptides were considered. The obtained data was normalized using the sum total peptide amount. For normalization and to derive protein abundances all quantified peptides per protein were used. Protein ratios and $\log_2$FC were calculated from the grouped protein abundances, without missing value imputation. To test for differentially abundant proteins, an ANOVA on individual proteins across all biological replicates (n=3) was performed. For further analysis, the 7665 ProteinGroups with a high-confidence score and quantitative values were used. For each drug-treated condition the $\log_2$FC and p-values were derived against DMSO/baseline control conditions. Depending on the duration of the treatment, either the 8 or the 18h negative control was used. The resulting processed datasets can be found in **Supplementary Data Table 2**.

#### Kinobead profiling

Cells were lysed in 0.8 % IGEPAL, 50 mM Tris-HCl pH 7.5, 5% glycerol, 1.5 mM magnesium choride, 150 mM sodium chloride, 1 mM sodium orthovanadate, 25 mM sodium fluoride, 1 mM DTT, protease inhibitors (SigmaFast, Sigma) and phosphatase inhibitors (prepared in-house according to Phosphatase inhibitor cocktail 1, 2 and 3 from Sigma-Aldrich). The cell lysate mixes used for compound profiling was generated either from COLO-205, K-562, SK-N-BE(2), MV-4-11 and OVCAR-8 lysates (standard 5 CL (cell line) mix) or Jurkat and MCF7 mixed in equivalent ratios, the protein concentration was determined by Bradford.

Kinobeads pulldown experiments were performed as previously described[18]. In brief, 2.5 mg of the cell lysate mixture was pre-incubated with increasing compound concentrations (DMSO, 3 nM, 10 nM, 30 nM, 100 nM, 300 nM, 1 µM, 3 µM, 30 µM) for 45 min at 4 °C in an end-over-end shaker in either of the two lysate mixes. Afterwards, lysates were incubated with Kinobeads (17 µL settled beads) for 30 min at 4 °C. The beads were washed and bound proteins were reduced with 50 mM DTT in 8 M Urea, 40 mM Tris HCl (pH 7.4) for 30 min at room temperature. After alkylation with 55 mM CAA proteins were digested with trypsin over night at 37 °C. Peptides were desalted using C18 StageTips and dried down in a SpeedVac. Peptides were analyzed via LC-MS/MS on a Dionex Ultimate3000 nano HPLC coupled online to an Orbitrap Fusion Lumos (Thermo Fisher Scientific) mass spectrometer. Peptides were delivered to a trap column (100 µm x 2 cm, packed in-house with Reprosil-Gold C18 ODS-3.5 µm resin, Dr. Maisch, Ammerbuch) and washed at a flow rate of 5 µl/min in solvent A (0.1% formic acid, 5 % DMSO in HPLC grade water). Peptides were then separated on an analytical column (75 µm x 40 cm, packed in house with Rprosil-Gold C18 3 µm resin, Dr. Maisch, Ammerbuch) using a 52 min gradient ranging from 4-32% solvent B (0.1% formic acid, 5% DMSO in acetonitrile) in solvent A at a flow rate of 300 nL/min. The mass spectrometer was operated in a data dependent mode, automatically switching between MS1 and MS2 spectra. MS1 spectra were acquired over a mass-to-charge ratio (m/z) range of 360-1300 m/z at a resolution of 60.000 in the Obitrap using a maximum injection time of 50 ms and an automatic gain control (AGC) target value of 4e5. Up to 12 peptide precursors were isolated (isolation width of 1.7 Th, maximum injection time of 75 ms, AGC value of 5e4), fragmented by HCD using 30% normalized collision energy (NCE) and analyzed in the Orbitrap at a resolution of 15,000. The dynamic exclusion duration of fragmented precursor ions was set to 30s.

Peptide and protein identification and quantification was performed using MaxQuant (1.5.3.30) by searching the tandem MS data against all canonical protein sequences as annotated in the UniProtKB reference database using the embedded search engine Andromeda. Carbamidomethylated cysteine was set as fixed modification and phosphorylation of serine, threonine and tyrosine, oxidation of methionine and N-terminal protein acetylation as variable modifications. Trypsin/P was specified as the proteolytic enzyme and up to two missed cleavages were allowed. The minimum length of amino acids was set to seven and all data were adjusted to 1% PSM and 1% protein FDR. Label-free quantification and match between runs were enabled within MaxQuant. For the Kinobeads competition binding assays, protein intensities were normalized to the respective DMSO control and IC50 and EC50 values were deduced by a four-parameter log-logistic regression using an internal pipeline that utilizes the drc package in R. An apparent dissociation constant (Kdapp) was calculated by multiplying the estimated EC50 with a protein-dependent correction faction. The correction factor of a protein is defined as the ratio of the amount of protein captured from two consecutive pulldowns of the same DMSO control lysate. Targets of the compounds are annotated manually. A protein is considered a target if the resulting binding curve shows a sigmoidal curve shape with a dose dependent decrease of binding to the beads. Additionally, the number of unique peptides and MSMS counts per condition as well as the protein intensity in the DMSO control are taken into account. The resulting fitted parameters in addition to the normalised intensities can be found in **Supplementary Data Table 5**.

### SI Tables

#### Table S1 Primers for mutagenesis

| Vector name | Primer fw | Primer rev | Method |
|---|---|---|---|
| pENTR223 EGFR T790M | TGCAACTCATCATGCAGCTCATGCCCTTC | GAAGGGCATGAGCTGCATGATGAGTTGCA | PCR |
| pENTR223 EGFR G719S | AAGATCAAAGTGCTGAGCTCCGGTGCGTTCG | CGAACGCACCGGAGCTCAGCACTTTGATCTT | PCR |
| pENTR223 EGFR L858R | CACAGATTTTGGGCGGGCCAAACTGCTGGGTGCGGAAG | CAGCAGTTTGGCCCGCCCAAAATCTGTGATCTTGACATGCT GCG | PCR |
| pENTR223 EGFR L861Q | GCTGGCCAAACAGCTGGGTGCGGAAGAGAAAGAATAC | TCCGCACCCAGCTGTTTGGCCAGCCCAAAATCTGTG | PCR |
| pENTR223 EGFR L718V | AAGATCAAAGTGGTGGGCTCCGGTGCGTTCG | CGAACGCACCGGAGCCCACCACTTTGATCTT | PCR |
| pENTR223 BRAF V600E | GATTTTGGTCTAGCTACAGAGAAATCTCGATGGAGTG | CACTCCATCGAGATTTCTCTGTAGCTAGACCAAAATC | PCR |
| pENTR223 RAF1 S256L | CTCTCCCAGAGGCAGAGGCTGACATCCACACCTAATGTCCA C | CATTAGGTGTGGATGTCAGCCTCTGCCTCTGGGAGAGGGAA C | PCR |
| pENTR223 RET M918T | GATTCCAGTTAAATGGACGGCAATTGAATCCCTTTTTGATCAT ATC | TCAAAAAGGGATTCAATTGCCGTCCATTTAACTGGAATCCGA CCCTG | PCR |
| pENTR223 ERBB4 R711C | CAGCACCCAATCAAGCTCAACTTTGTATTTTGAAAGAAAC | GTTTCTTTCAAAATACAAAGTTGAGCTTGATTGGGTGCTG | PCR |
| pENTR233 LYN Y32A | GCCGTGAGAGATCCAACGTCCAATAAAC | AATAGTTCTTTCAGTATTACGTACTG | SDM |
| pENTR233 LYN Y316A | GGAGCCCATTGCGATCATCACCGAGTAC | TCCCTGGTGACCACAG | SDM |
| pENTR233 LYN T319I | TCGAGTACATGGCCAAGGGCAG | TGATGATGTAAATGGGCTCCTCCCTG | SDM |
| pENTR233 CSK T266I | TGGAGTACATGGCCAAGGGGAG | TGACGATGTAGAGCCCGCCCTTC | SDM |
| pENTR223 BLK S24A | GGGCCAATGGGCGCCCCTGAAGG | TTGTCCTTCTCTTTGATCG | SDM |
| pENTR223 BLK T48A | CTTCAACCACCTTGCTCCTCCACCGCCCGATGAAC | CATCGGGCGGTGGAGGAGCAAGGTGGTTGAAGACAACCAG G | PCR |
| pENTR223BLK S29A | CCTGAAGGTCGCGGCCCAAGACAAGGAC | GGGCTCCATTGGCCC | PCR |
| pENTR223BLK C319S | AGCCTGCTGGATTTCCTGAAGACAGATG | TCCTCTGGCCATGTACTCGGTGAC | SDM |
| pENTR223 RIPK2 ΔCARD | GTGGTTTCTAGATCACCATC | GGCTATACCAGGCTGCAGAC | SDM |

SDM = site directed mutagenesis, PCR = direct PCR to transformation see section **Plasmids and cloning**

#### Table S2 Gibson primer sequences

| Vector name | Primer fw | Primer rev | Vector BB |
|---|---|---|---|
| pRRL_SFFV_LYN*_BFP_P 2A_mCherry | CGCCAGTCCTCCGAGTCGACGGCATGGGATGTATAAAATCA AAAGGGAA | CCCGAGCCACCTCCGGATCCAGGCTGCTGCTGGTATTGC | SR |
| pRRL_SFFV_CSK*_EF1a_i RFP670 | CGGCGCGCCAGTCCTCCGAGTCGACCATGTCAGCAATACA GGCCG | CTGACGGGCACCGGAGCCACTCGAGTCACAGGTGCAGCT CGTG | iRFP670 |
| pRRL_SFFV_BLK*_BFP_P 2A_mCherry | CGCCAGTCCTCCGAGTCGACGGCACCATGGGGCT | CCCGAGCCACCTCCGGATCCGGGCTGCAGCTCGTACTG | SR |
| pRRL_SFFV_BLK_S5A_BF P_P2A_mCherry | CGCCAGTCCTCCGAGTCGACGGCACCATGGGGCTGGTAG CGAGCAAAAAGCCGGACAAGGAAAAG | CCCGAGCCACCTCCGGATCCGGGCTGCAGCTCGTACTG | SR |
| pRRL_SFFV_BLK_S6A_BF P_P2A_mCherry | CGCCAGTCCTCCGAGTCGACGGCACCATGGGGCTGGTAAG TGCGAAAAAGCCGGACAAGGAAAAG | CCCGAGCCACCTCCGGATCCGGGCTGCAGCTCGTACTG | SR |
| pRRL_SFFV_SRC_BLK_FU S_BFP_P2A_mCherry | CGGCGCGCCAGTCCTCCGAGTCGACGCCACCATGGGTAG CAACAAGAGCAAGC | ACCACGAAATGCTTGTCTTCGGCCAGCGGGCCCGCC | SR |
| | GAAGACAAGCATTTCGTGGTGGC | CCCGAGCCACCTCCGGATCCGGGCTGCAGCTCGTACTG | |
| pRRL_SFFV_RIPK2*_BFP_ P2A_mCherry | CGCCAGTCCTCCGAGTCGACGGCACCATGAACGGGG | CCCGAGCCACCTCCGGATCCGGGCTGCAGCTCGTACTG | SR |

BB = backbone ; SR = stability reporter (pRRL_SFFV_empty_BFP_P2A_mCherry); iRFP670 = pRRL_SFFV_DCAF16_EF1a_iRFP670; * or mutated versions thereof

#### Table S3 sgRNA sequences

| KO | sgRNA sequence 1 | sgRNA sequence 2 (dual only) | Dox |
|---|---|---|---|
| CBL | GCAGGTCTAAGATATAAGGT | – | 96h |
| CBL-B | GTCCCAAACACATCCCGA | – | 96h |
| AAVS1 | GCTGTGCCCCGATGCACAC | – | 96h |
| CBL | GCCAGGATGAGTTACAGCA | GCTGTGCCCCGATGCACAC | 1 week |
| CBL-B | GCTCCGGAAAGAGCATCCT | GTTGCACTCGATTGGGACAG | 1 week |
| CBL & CBL-B | GCCAGGATGAGTTACAGCA | GTTGCACTCGATTGGGACAG | 1 week |
| AAVS1 | GCTCCGGAAAGAGCATCCT | GCTGTGCCCCGATGCACAC | 1 week |
| PSENEN | GTTGACCAACCAGAGAAA | – | 1 week |
| TMUB1 (1) | GGAGCCAATGGTGTCGTG | – | 96h |
| TMUB1 (2) | GCACGAGGGGCTCCTGCG | – | 96h |
| PSMB5 (1) | TTTGTACTGATACACCATGT | – | 48h |
| PSMB5 (2) | GCTTCATGGAACAACCACCC | – | 48h |
| FIP200 (1) | TTTCTAACAGCTCTATTACG | – | 96h |
| FIP200 (2) | ACTACGATTGACACTAAAGA | – | 96h |

**Table S4: Reagents and ressources for MS experiments** of sections "Sample preparation for full proteome profiling", "LC-MS/MS data acquisition of BioID and full proteome samples", "Data analysis and representation of BioID data" and "Processing and data analysis of full proteome profiling data"

| REAGENT or RESOURCE | SOURCE | IDENTIFIER |
|---|---|---|
| LiChrosolv® Water | MERCK KgaA, Germany | # 1.15333.2500 |
| PMSF | BioChemica | # A0999.0005 |
| Protease inhibitor cocktail | Sigma-Aldrich, Germany | # P8340 |
| SDS | SERVA Electrophoresis GmbH, Germany | #20765 |
| HEPES | Merck, KgaA, Germany | # 1.10110.0250 |
| Covaris S2 high performance ultrasonicator | Covaris, USA | NA |
| Dithiothreitol, for molecular biology, minimum 99% titration | Sigma-Aldrich, Germany | # D9779-5G |
| Urea | Sigma-Aldrich, Germany | # U0631-500G |
| Tris(hydroxymethyl)aminomethane (Tris/HCl) ultrapure grade ≥99.9% | Sigma-Aldrich, Germany | # 154563 |
| Microcon 30, Ultracel YM-30 | Merck Millipore, Ireland | # MRCF0R030 |
| Iodoacetamide | Sigma-Aldrich, Germany | # I1149-5G |
| Triethylammonium Bicarbonate Buffer (TEAB), 1 M, pH 8.5 | Sigma-Aldrich, Germany | # 17902 |
| Sequencing Grade Modified Trypsin | Madison, WI, USA | # V5113 |
| NaCl | Sigma-Aldrich, Germany | # S7653 |
| PEPTIDE DESALTING COLUMNS 50PC | Thermo Fisher Scientific, USA | #89851 |
| TMTpro 18-plex label reagent set | Thermo Fisher Scientific, USA | # A52045 |
| Agilent 1200 series | Agilent | NA |
| 150 x 2.0 mm Gemini-NX, 3 µm C18 110Å | Phenomenex, Torrance, USA | NA |
| TFA Uvasol (trifluoroacetic acid) | MERCK KgaA, Germany | # 1.08262.0100 |
| 0.1% TFA | Sigma-Aldrich, Germany | # 34978-2.5L-R |
| Orbitrap Fusion Lumos Tribrid mass spectrometer | Thermo Fisher Scientific, USA | NA |
| **MS data acquisition** | | |
| Dionex Ultimate 3000 RSLCnano system | Thermo Fisher Scientific, USA | NA |
| Nanospray Flex Ion Source | Thermo Fisher Scientific, USA | NA |
| PepMap 100 C18, 5 µm, 5 × 0.3 mm | Thermo Fisher Scientific, USA | NA |
| Fused Silica Capillary ID 75 µm, 50 cm length | Polymicro Technologies LLC | TSP075150 |
| ESI Emitter Fused Silica: 20 µm ID x 7 cm L x 365 µm OD; Orifice ID: 10 µm | CoAnn Technologies, USA | TIP1002010C-5 |
| ReproSil-Pur 120 C18-AQ, 3 µm | Dr. Maisch, Germany | # r13.aq. |
| Suprapur ® Formic acid 98-100% | Merck KgaA, Germany | # 1.11670.1000 |
| Acetonitril for HPLC LC-MS Grade | VWR Chemical, USA | # 83640.32 |
| **Data processing and analysis** | | |
| Discoverer version: v.2.4.1.15 | Thermo Fisher Scientific | NA |
| SAINTq | Teo G, et al.[16] | NA |
| R version 4.3.1 | The R Foundation | NA |

#### SI Figures

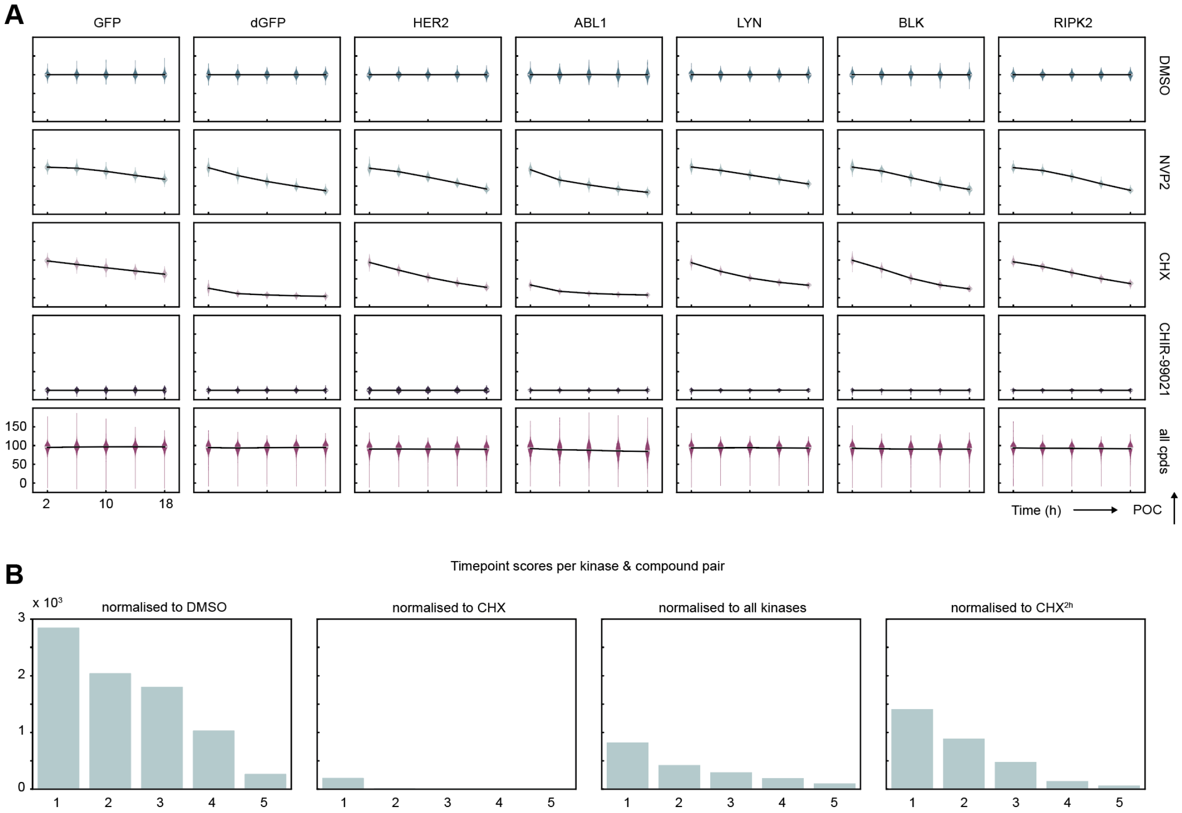


**Fig S1 Normalisation and overview of scoring distribution. A** Selected examples for kinases and controls highlighted throughout the paper. Depiction of the screening trajectories as line (black) and violin plot (kernel density estimate (KDE) for data including a horizontal grey line highlighting the median) for all the controls and averaged across the screened compounds (all cpds, not including the PROTAC controls). DMSO is shown in green, NVP2 in light green, CHX (cycloheximide) in light pink, CHIR-99021 in dark lilac and the average of all cpds in pink. DMSO and CHIR-99021 formed the basis of the data normalization to generate the POC values. CHX trajectories were used for determining each reporter's half-life. **B** Score overview of each kinase and compound pair across the different time points for each chosen normalization strategy. A score of 5 indicates the kinase-cpd trajectory scored for each recorded timepoint, a score of 1 indicates scoring for a single timepoint across the recorded trajectory. Recorded time stamps: 2,6,10,14 and 18h.

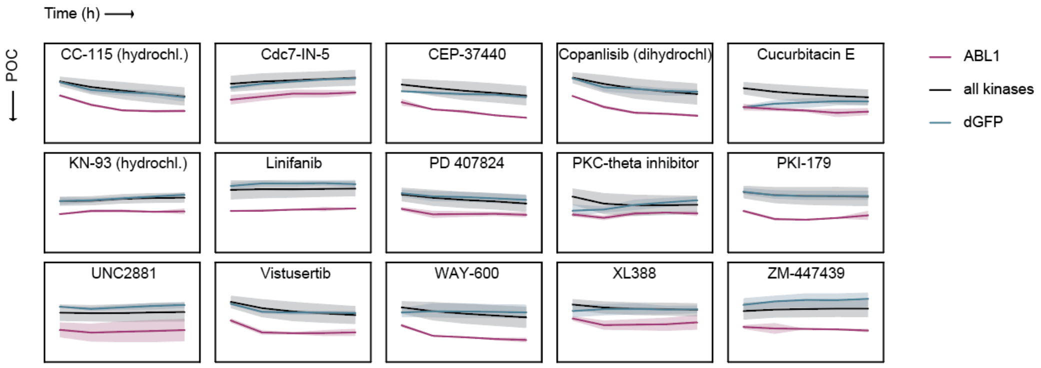


**Fig S2 Continuation from Extended Data Fig 1G.** Drug screening data comparing ABL1 to the mean of all other kinases and the dGFP control trajectory (m = 2, error bars correspond to the CI for individual trajectories and the SD for the mean of all kinases).

Gating strategy: iCas9 GFP with Thy1.1 Neo sgRNAs & sorted/unsorted reporter examples

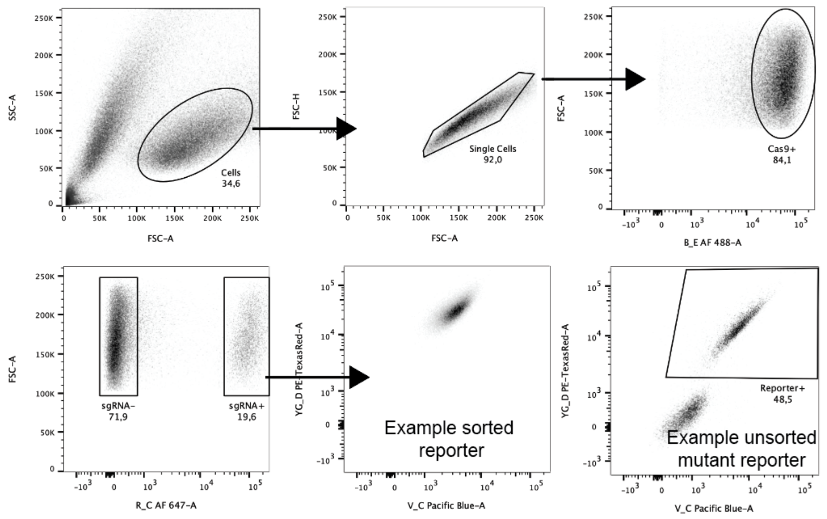


Gating strategy: CRISPR screen

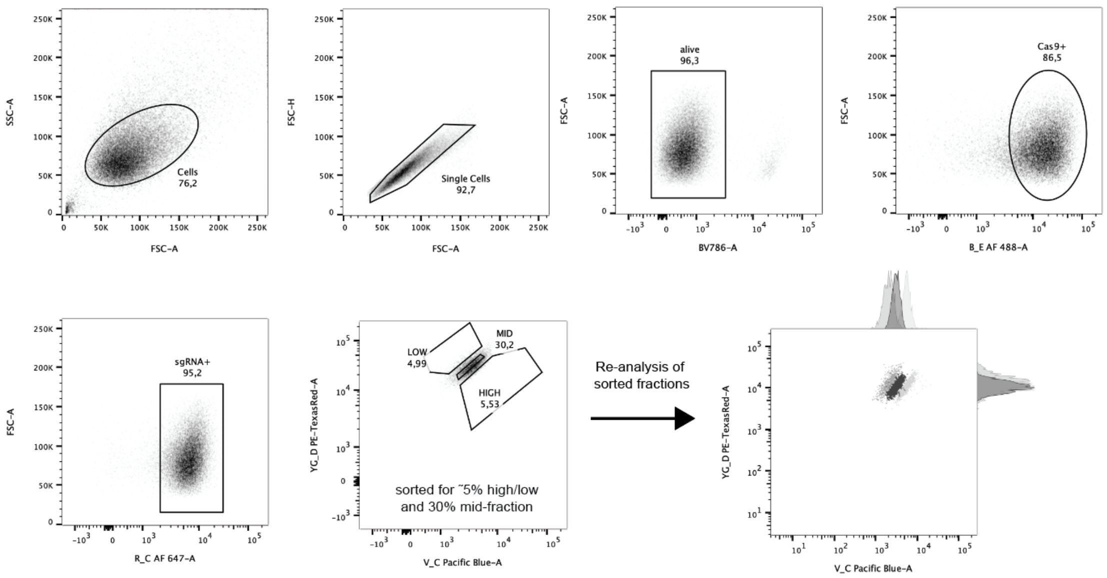


**Fig S3 Gating strategies applied across Fig 2-4 and Extended Fig 3-5. Top:** Hierarchy of applied gating scheme for either sorted reporters (pool and clonal or unsorted cell lines. iCas9+ (AF488/GFP) and Thy1.1+ (AF647) gating was skipped in case of non-induced Cas9 (non-induced k.o. experiments), and/or in case cells had been selected for harboring the sgRNA using Neomycin. **Bottom:** Gating strategy applied for all CRISPR/Cas9 screens and exemplified re-analysis of the sorted fraction after FACS.

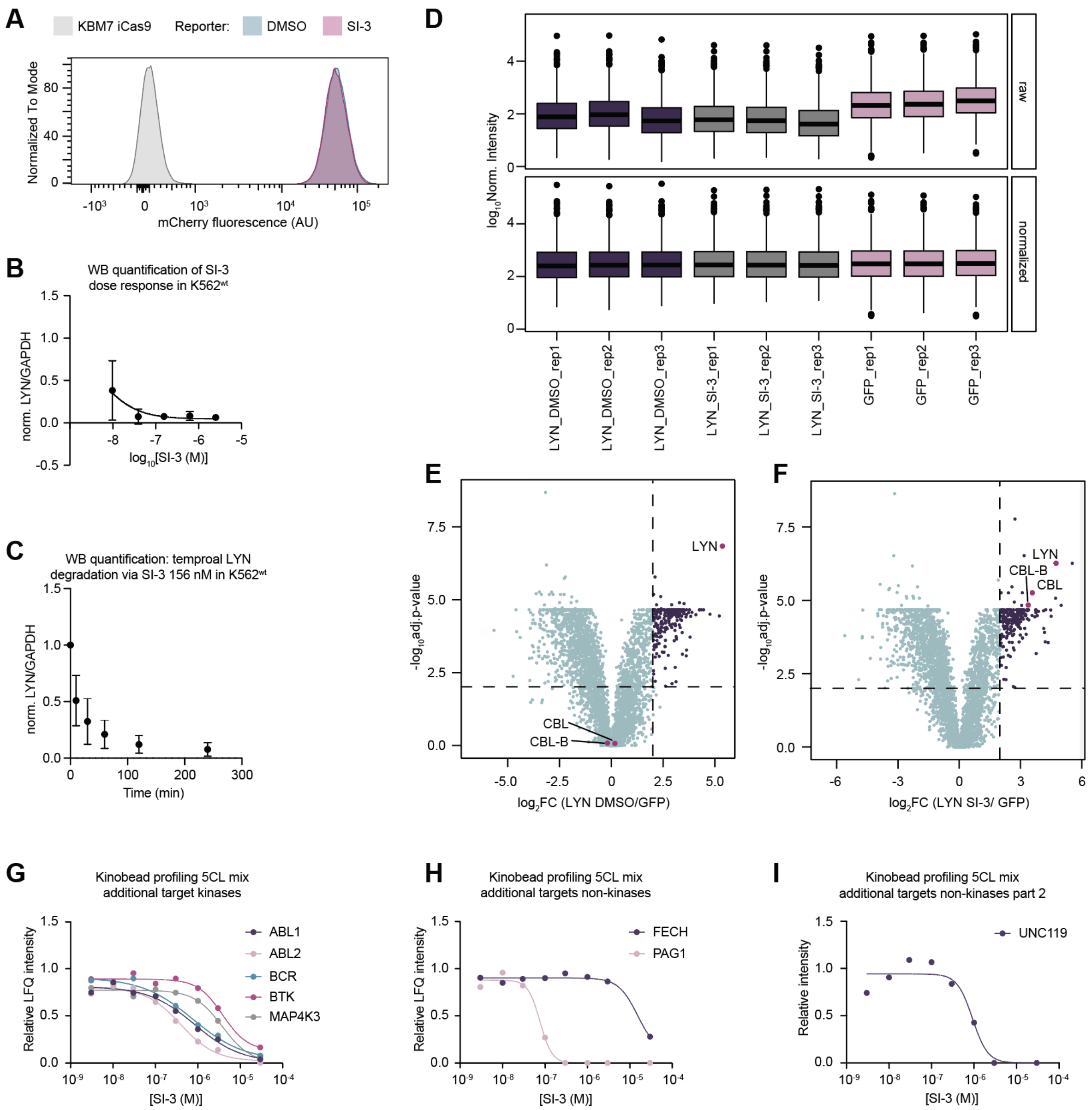


**Fig S4 Additional data to LYN datasets. A** Flow histogram matched to **Fig 2A** showing no change of mCherry fluorescence upon SI-3 treatment (8h, 156 nM). **B** Quantification of endogenous dose response measured via immunblot at 2h after treatment of SI-3 with varying concentrations (9.77 nM – 2.5 µM, see example replicate in **Fig 2B**). Black line indicates the nonlinear fit of a dose response (inhibitor vs response with three parameters). $D_{max}$ = 95.55%, IC50 = 4.595 nM, $R^2$ = 0.8767. Data is shown relative to the GAPDH loading control and normalised to DMSO. **C** Quantification of data shown in **Fig 2C** and respective replicates monitoring endogenous LYN degradation for 156 nM SI-3 treatment over time by immunblot. Data is shown relative to the GAPDH loading control and normalised to DMSO. **D** Distribution of protein intensities ($log_{10}$) before and after total ion current (TIC) normalization across LYN, LYN SI-3 treated and GFP negative controls. Solid line represents the median, box limits show the IQR and its whiskers 1.5 x IQR. Outliers are visualized as black dots (n = 3). **E** Differential enrichment of proteins in LYN DMSO control against GFP negative controls. The x-axis represents the $log_2FC$ of each protein against the signal in the GFP negative control. The y-axis shows the $-log_{10}$ of the adjusted p-value (BH correction). As thresholds for significantly enriched proteins a $log_2FC$ of larger or equal to 2 and an adjusted p-value smaller or equal to 0.01 were used (dotted lines). Not significantly enriched proteins are shown in light green, whereas significantly enriched proteins are indicated in purple. The LYN kinase bait protein, and the two E3-ligases CBL and CBL-B are shown in pink. **F** Differential enrichment of proteins in LYN SI-3 treated against GFP negative controls. The x-axis represents the $log_2$FC of each protein against the signal in the GFP negative control. The y-axis shows the -$log_{10}$ of the adjusted p-value (BH correction). As thresholds for significantly enriched proteins a $log_2$FC of larger or equal to 2 and an adjusted p-value smaller or equal to 0.01 were used (dotted lines). Not significantly enriched proteins are shown in light green, whereas significantly enriched proteins are indicated in purple. The LYN kinase bait protein, and the two E3-ligases CBL and CBL-B are highlighted in pink. n = biological replicates. **G** Kinobead profiling for additional kinase examples for SI-3 in the standard 5-cell line lysate (5CL) mix showing the relative LFQ intenstiy (y-axis) against the drug concentration (x-axis). Following $K_D$s were determined: ABL1 = 353 nM, ABL2 = 40.2 nM, BCR = 127 nM, BTK = 3.08 µM, MAP4K3 = 9.70 µM. **H** Plot as in G for for additional non-kinase examples Following $K_D$s were determined: FECH = 6.35 µM, PAG1 = 48.4 nM **I** Plot as in G but modified to show the relative intensity (y-axis) for UNC119, $K_D$ = 24.1 nM.

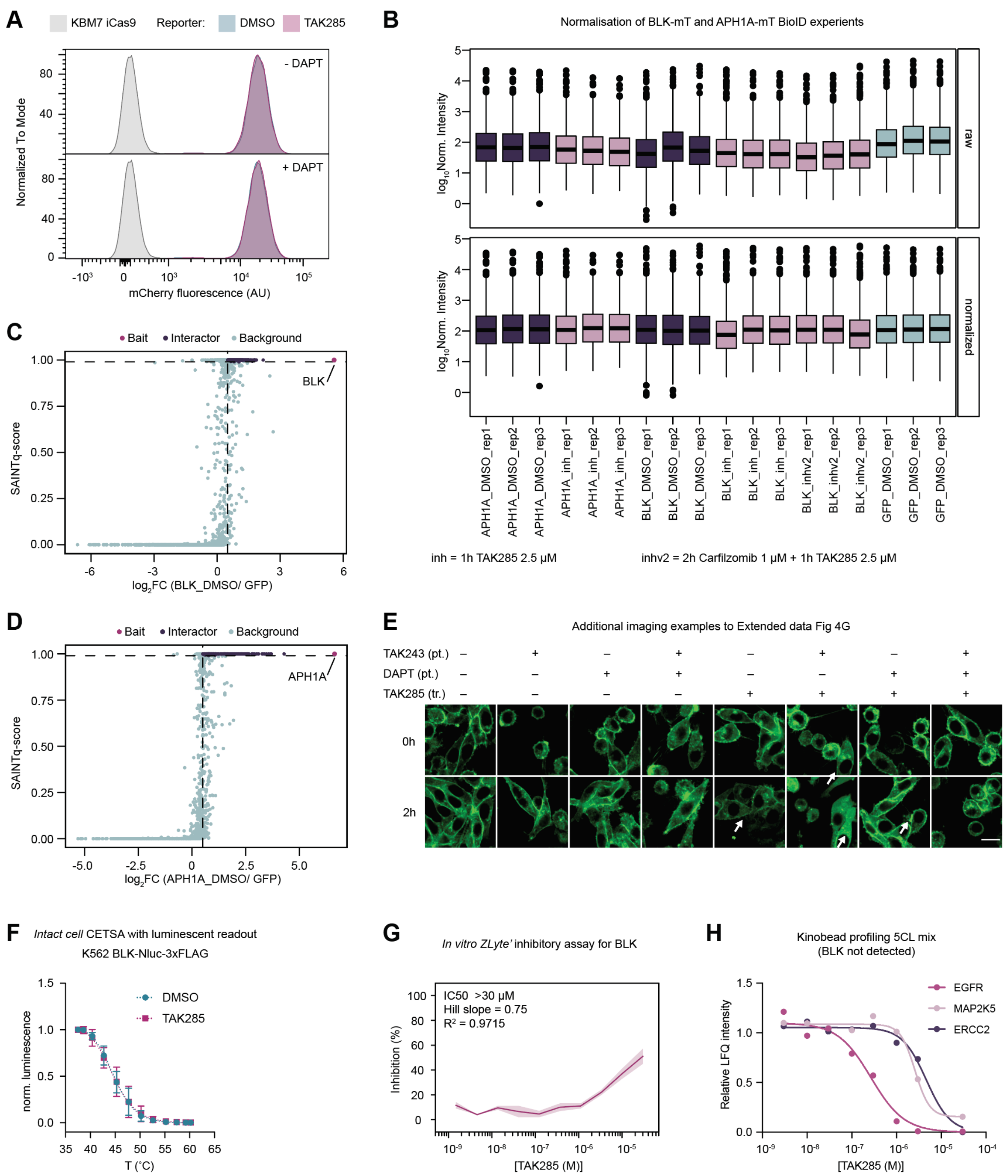


**Fig S5 Additional data to datasets related to BLK. A** Flow histogram matched to **Extended Data Fig 4C** showing no change of mCherry fluorescence upon DAPT (pre-treated 2h, 12.5 μM) or TAK285 (6h, 2.5 μM) treatment. **B** Distribution of protein intensities ($log_{10}$) before and after total ion current (TIC) normalization across BLK, APH1A and GFP negative controls including baseline (DMSO) and inhibitor treated conditions. Solid line represents the median, box limits show the IQR and its whiskers 1.5 x IQR. Outliers are visualized as black dots (n = 3). **C** Scatterplot of interaction partners after SAINTq scoring for BLK. The $log_2$FC against GFP negative controls (x-axis) was plotted against the SAINTq-score (y-axis). Proteins with a $log_2$FC of larger or equal to 0.5, a SAINTq score higher or equal to 0.99 with a BFDR smaller or equal to 0.01 and a CRAPome frequency below 10% were classified as interaction partners (pink). Proteins passing the scoring threshold but having a high CRAPome frequency were flagged together with the non-enriched proteins as background (light green) (n = 3). **D** Identical scatterplot of scoring characteristics for APH1A BioID experiments (n = 3). **E** Additional representative images of imaging experiment depicted in **Extended Data Fig 4G** (n = 2). **F** Luminescence-based CETSA experiment depicted as normalised luminescence to each treatment at 37˚C. K562 BLK-Nluc-3xFLAG cells were treated for 2h with TAK243 followed by 1h DMSO or 10 µM TAK285. $T^M$: DMSO = 44.60˚C, TAK285 = 44.43˚C; Hill-slopes: DMSO = -20.16, TAK285 = -18.52; R2 value: DMSO = 0.9829, TAK285 = 0.9718; no significant difference, multiple unpaired t-tests (n = 3). Values are shown as mean of the replicates ± SD. **G** ZLyte' *in vitro* inhibition assay data for BLK (m = 2). **H** Kinobead profiling data for the three hits identified for TAK-285 in the standard 5-cell line lysate mix. The relative LFQ intenstiy is depicted on the y-axis and the drug concentration on the x-axis. Following $K_D$s were determined: EGFR = 290 nM, ERCC2 = 4.70 µM, MAP2K5 = 2.55 µM. Data depicts the mean ± CI. n = biological replicates, m = technical replicates.

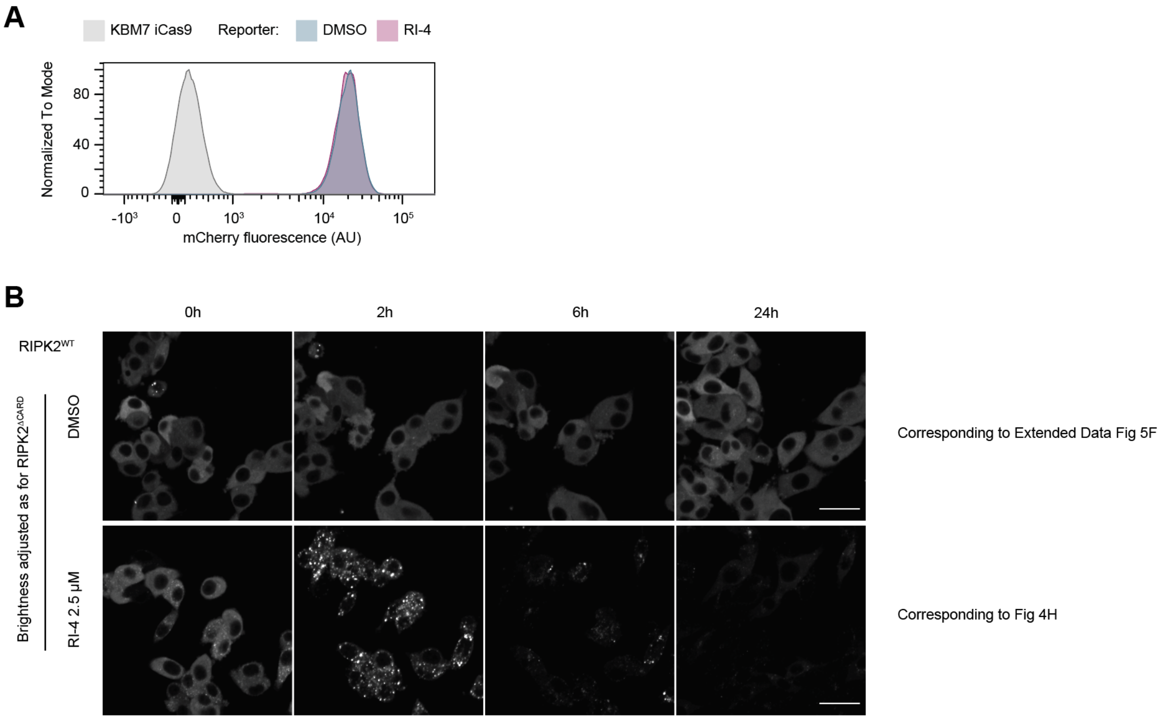


**Fig S6 Additional data to RIPK2 datasets. A** Flow histogram matched to **Fig 4A** showing no change of mCherry fluorescence upon RI-4 treatment (18h, 2.5 µM). **B** Data matched to the indicated figure panels with adjusted brightness values equal to the $RIPK2^{\Delta CARD}$ images.

**SI Fig S7-18** have been annotated in the following way: Uncropped images are shown as overlay of luminescent and colorimetric images. Replicates 2 and 3 have been annotated for clarity. Ladders are labelled with numeric values corresponding to the size (in kDA). For originals, the blots are labelled with their corresponding antibody. Labelled blots include additional information for each depicted band e.g. for protein fusions. Saturated pixels are depicted in red.

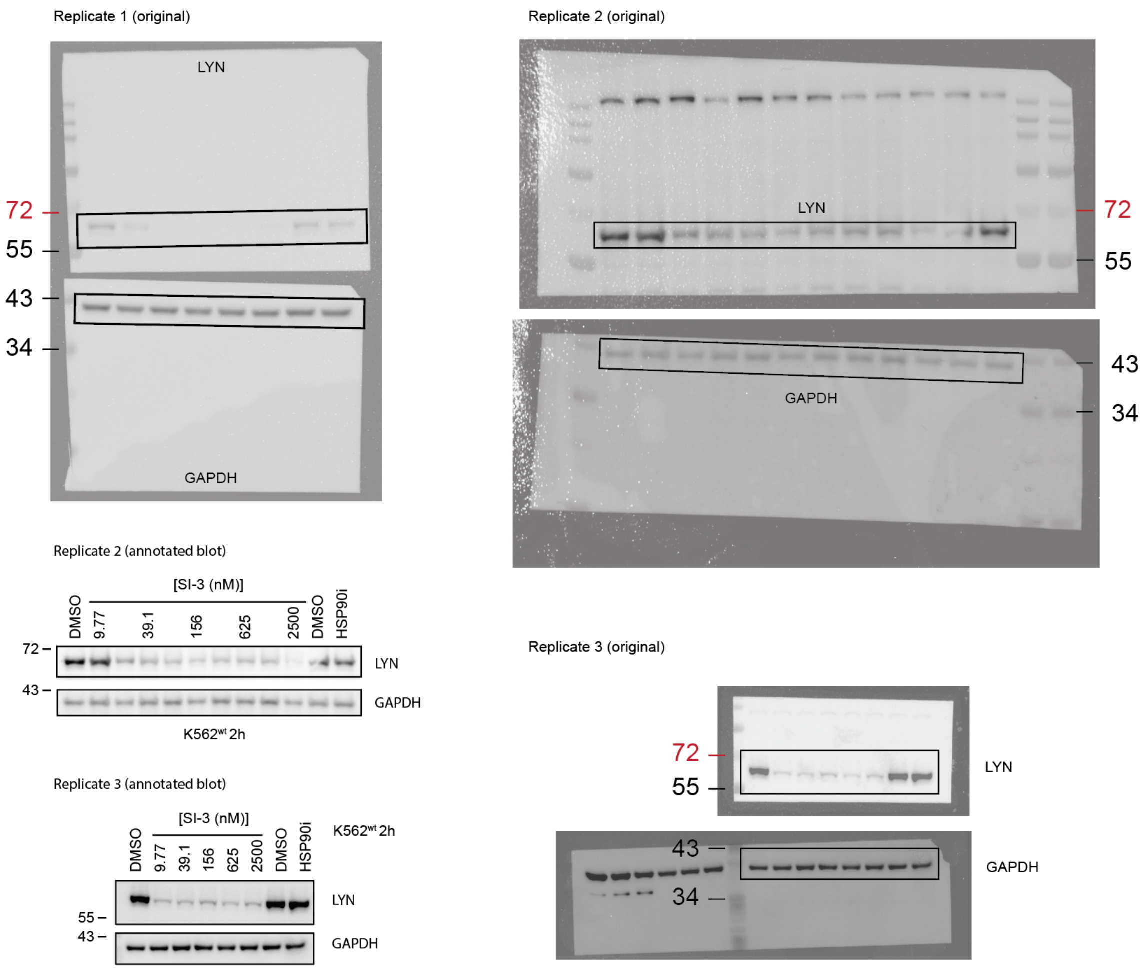


**Fig S7 Uncropped immunoblots and further replicates to Fig 2B.**

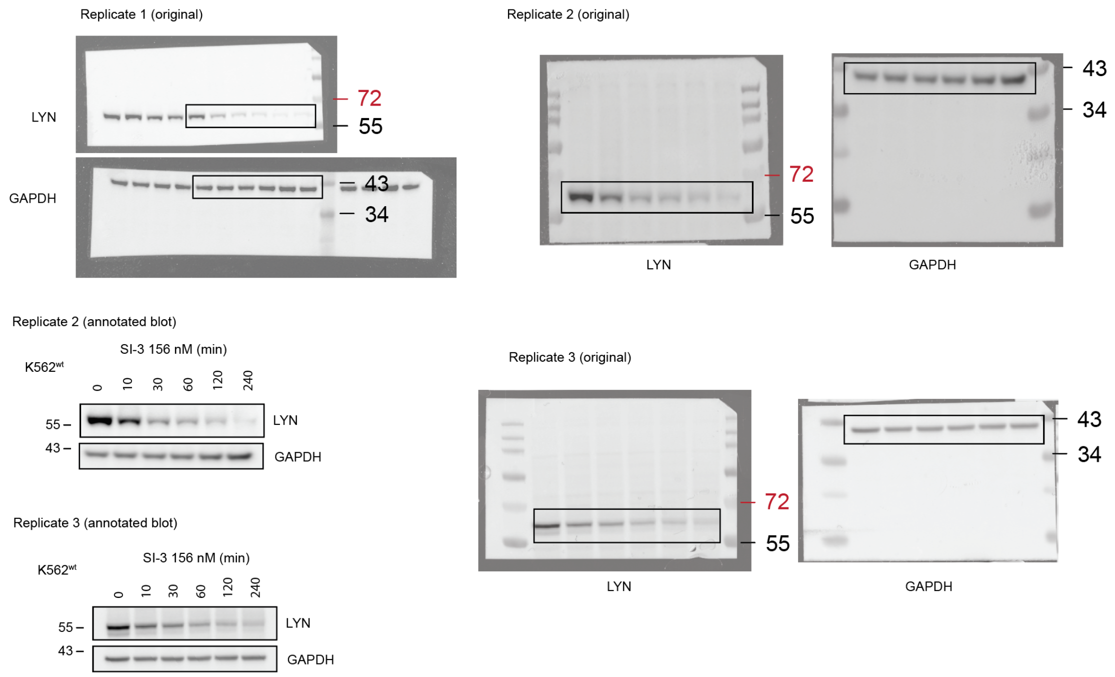


**Fig S8 Uncropped immunoblots and further replicates to Fig 2C.**

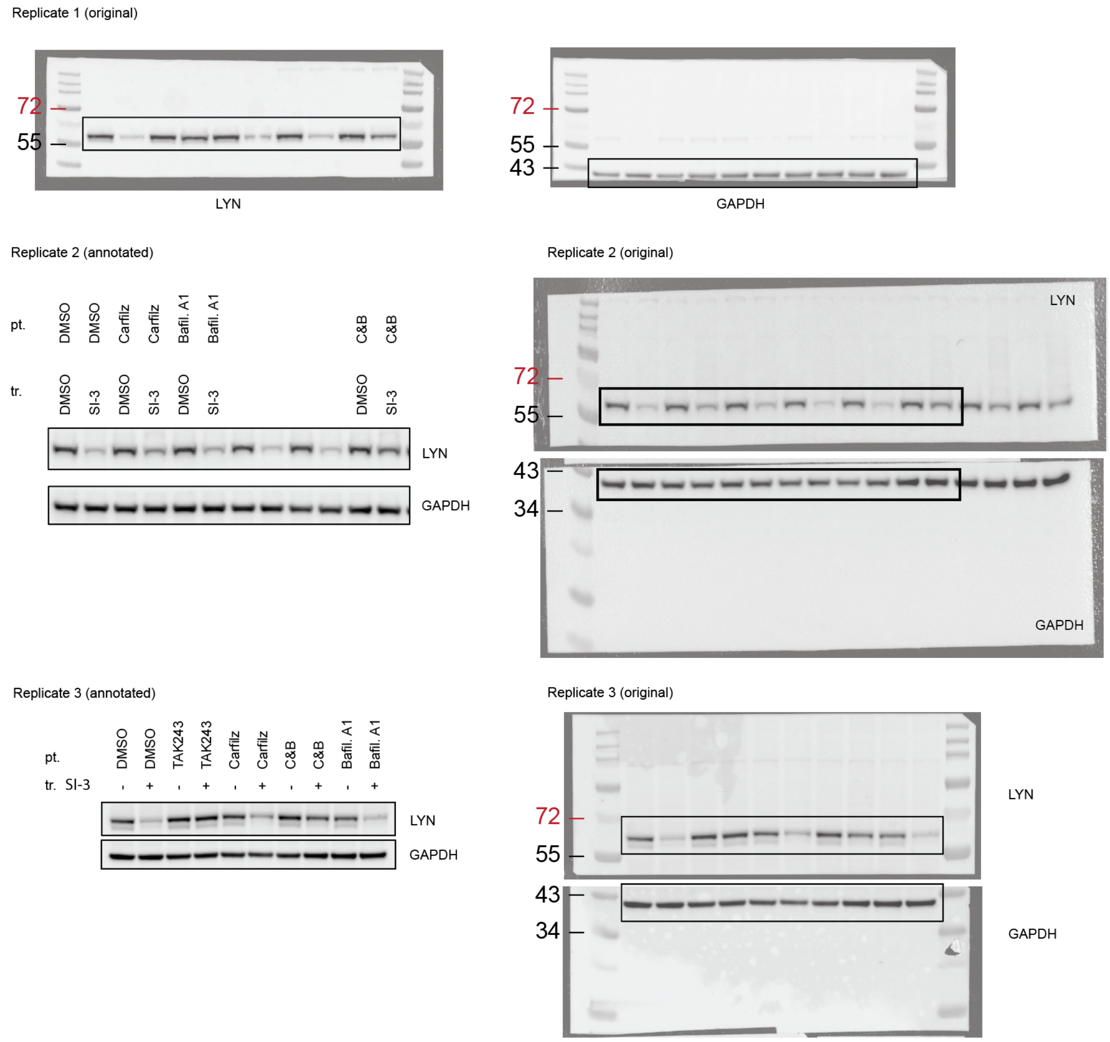


**Fig S9 Uncropped immunoblots and further replicates to Fig 2D.**

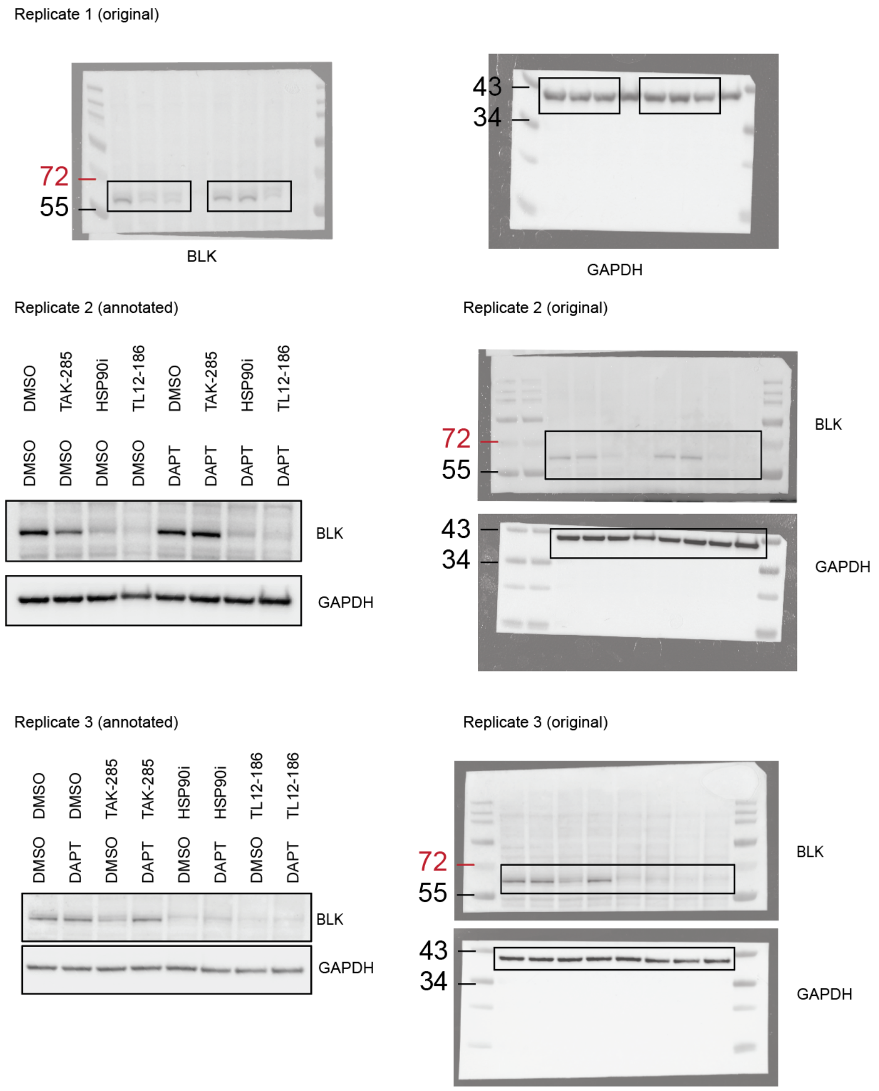


**Fig S10 Uncropped immunoblots and further replicates to Fig 3F.**

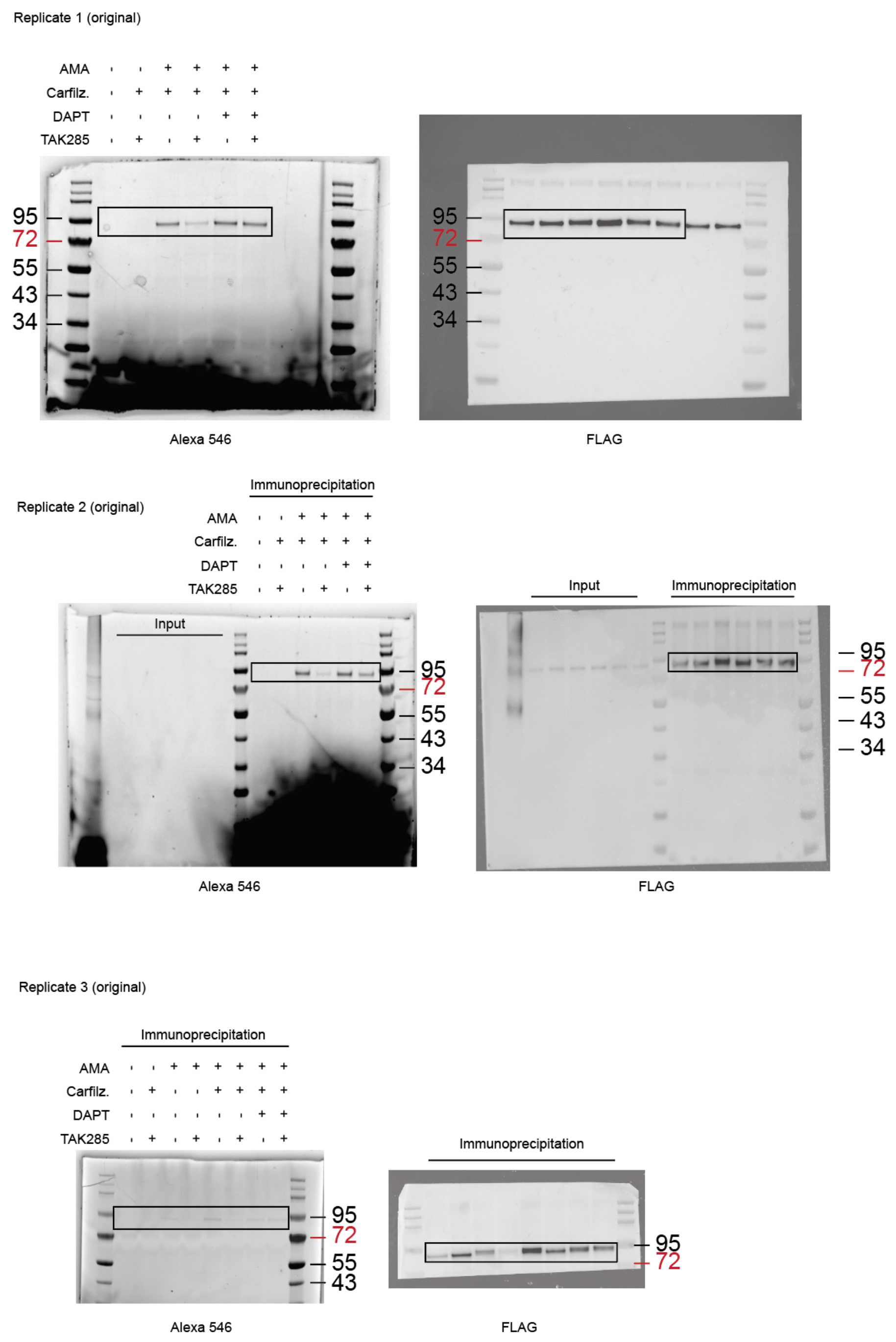


**Fig S11 Uncropped immunoblots and further replicates to Fig 3J.**

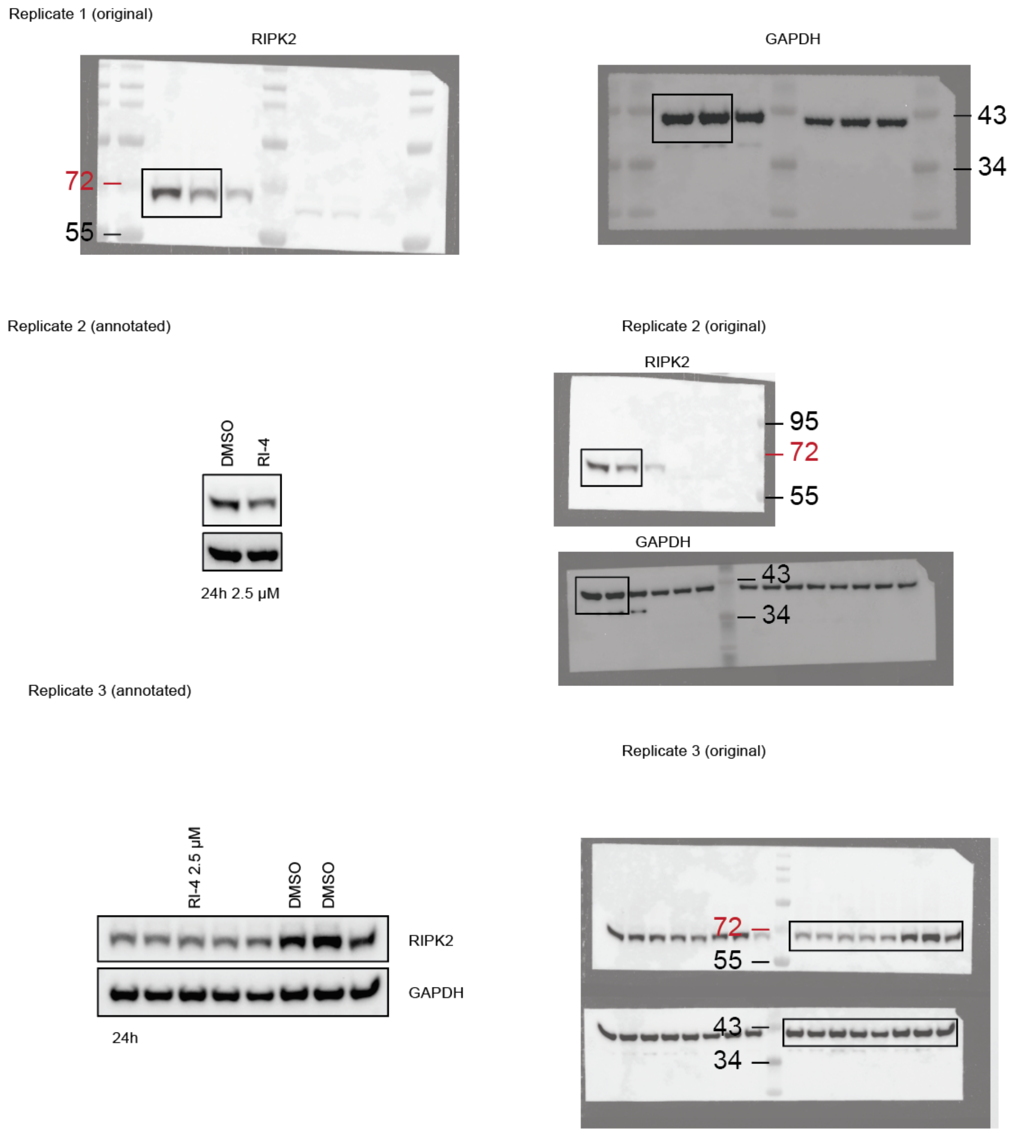


**Fig S12 Uncropped immunoblots and further replicates to Fig 4C.**

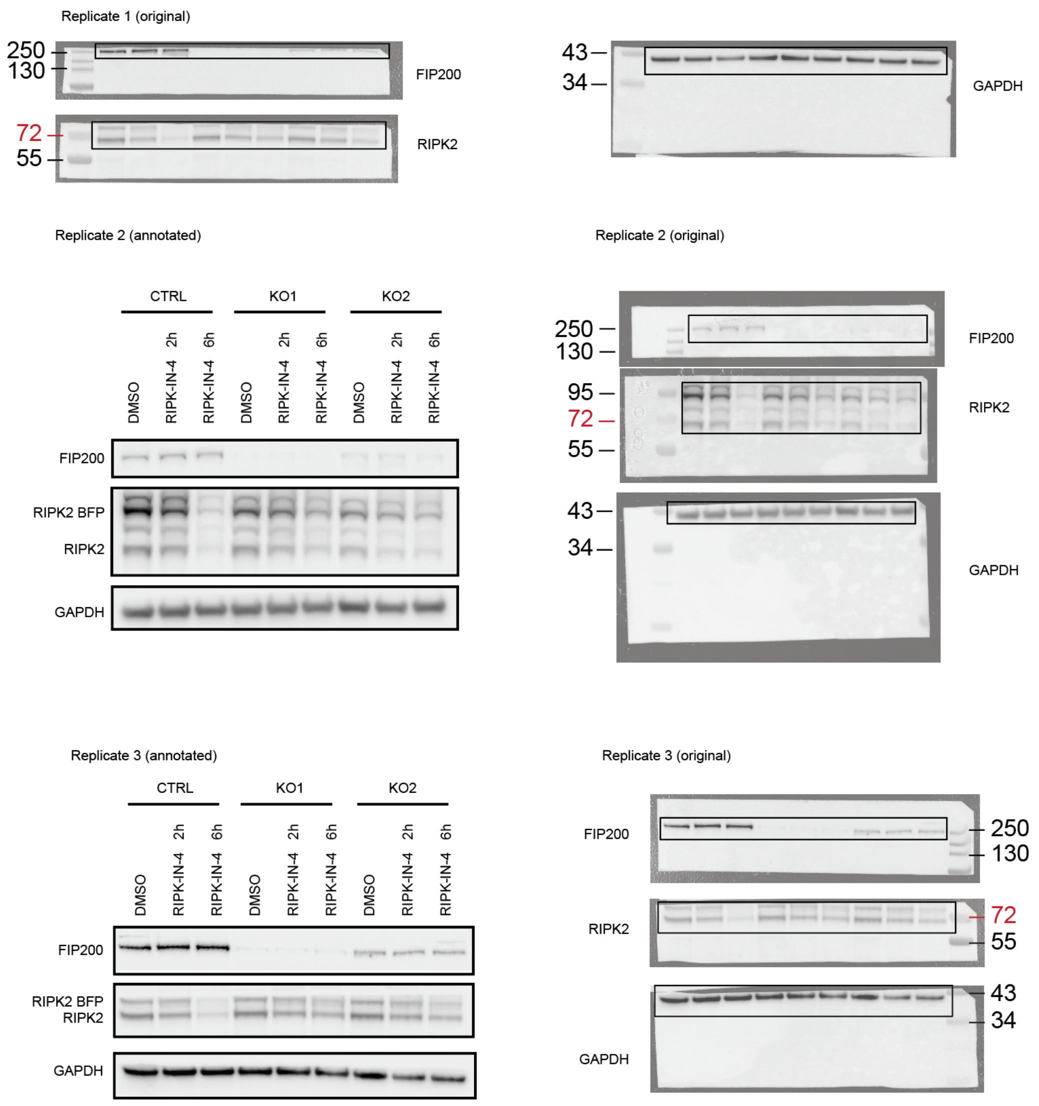


**Fig S13 Uncropped immunoblots and further replicates to Fig 4F.**

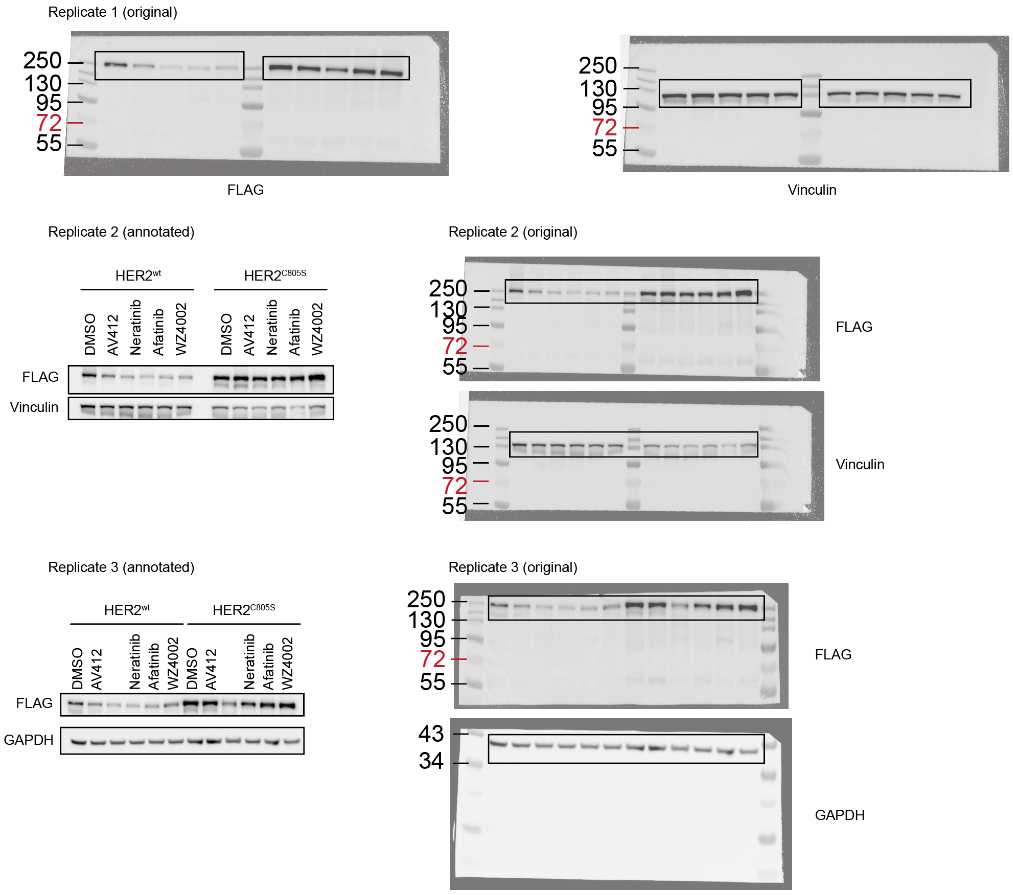


**Fig S14 Uncropped immunoblots and further replicates to Extended Data Fig 1D.**

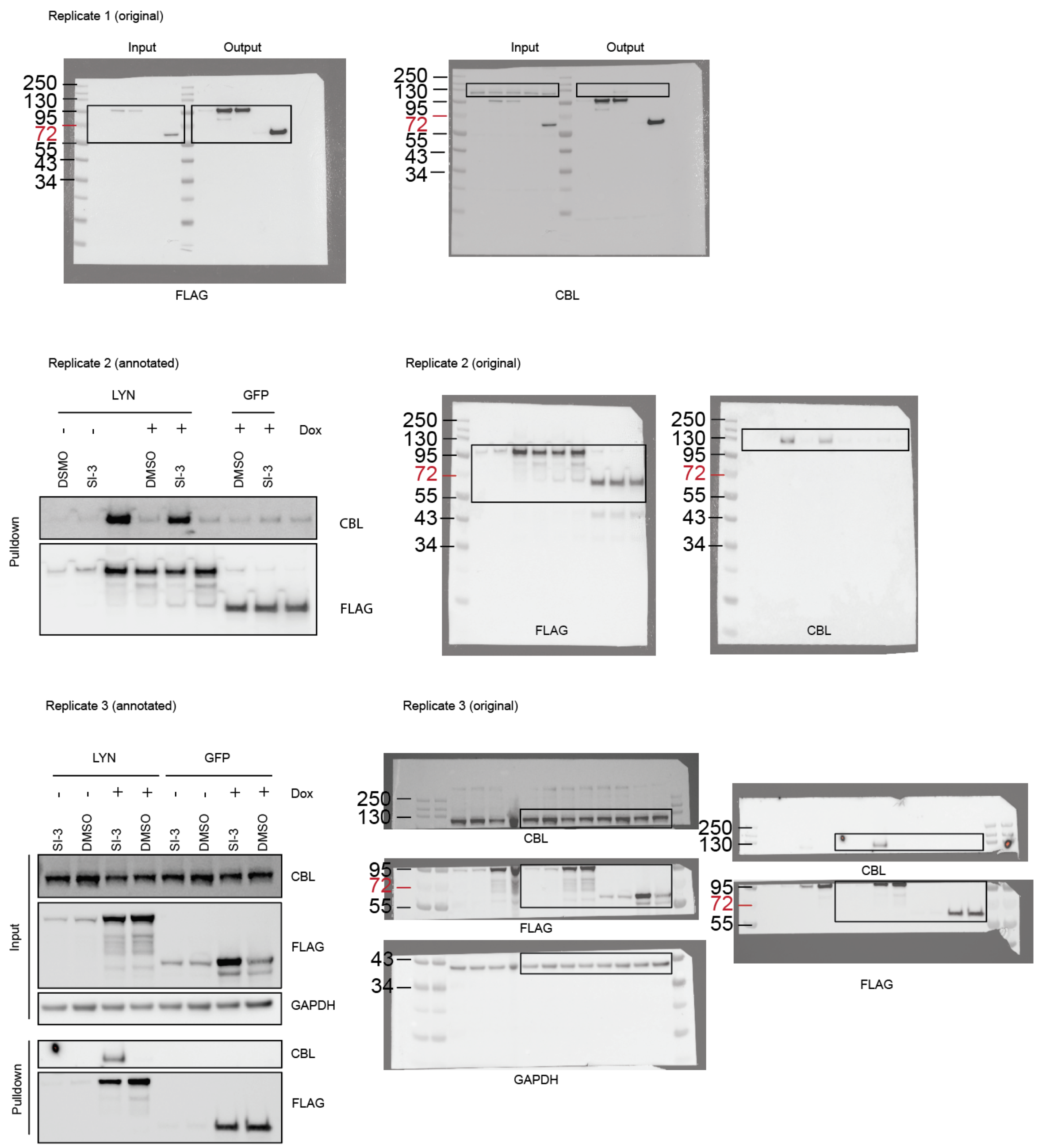


**Fig S15 Uncropped immunoblots and further replicates to Extended Data Fig 3G.**

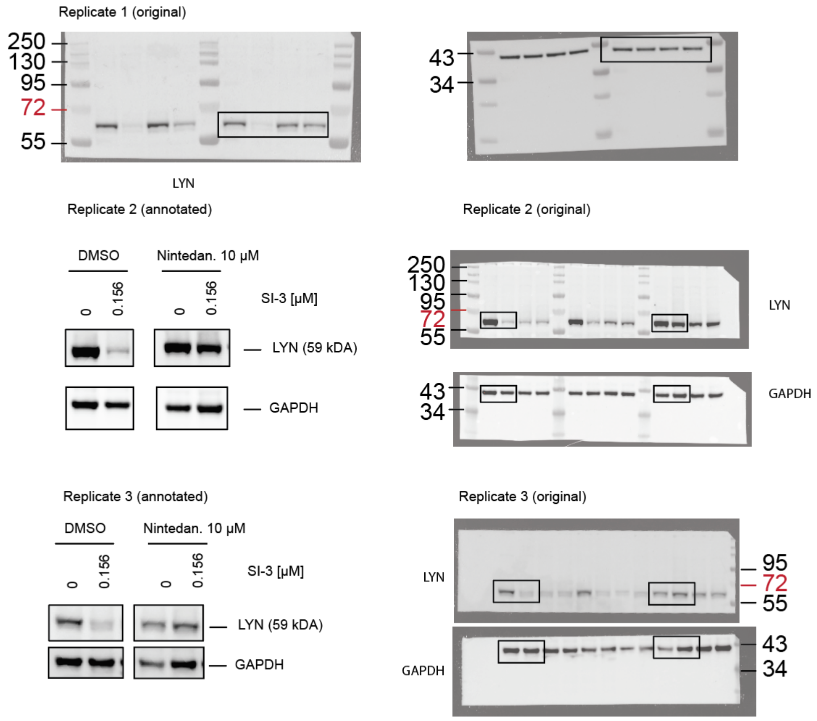


**Fig S16 Uncropped immunoblots and further replicates to Extended Data Fig 3K.**

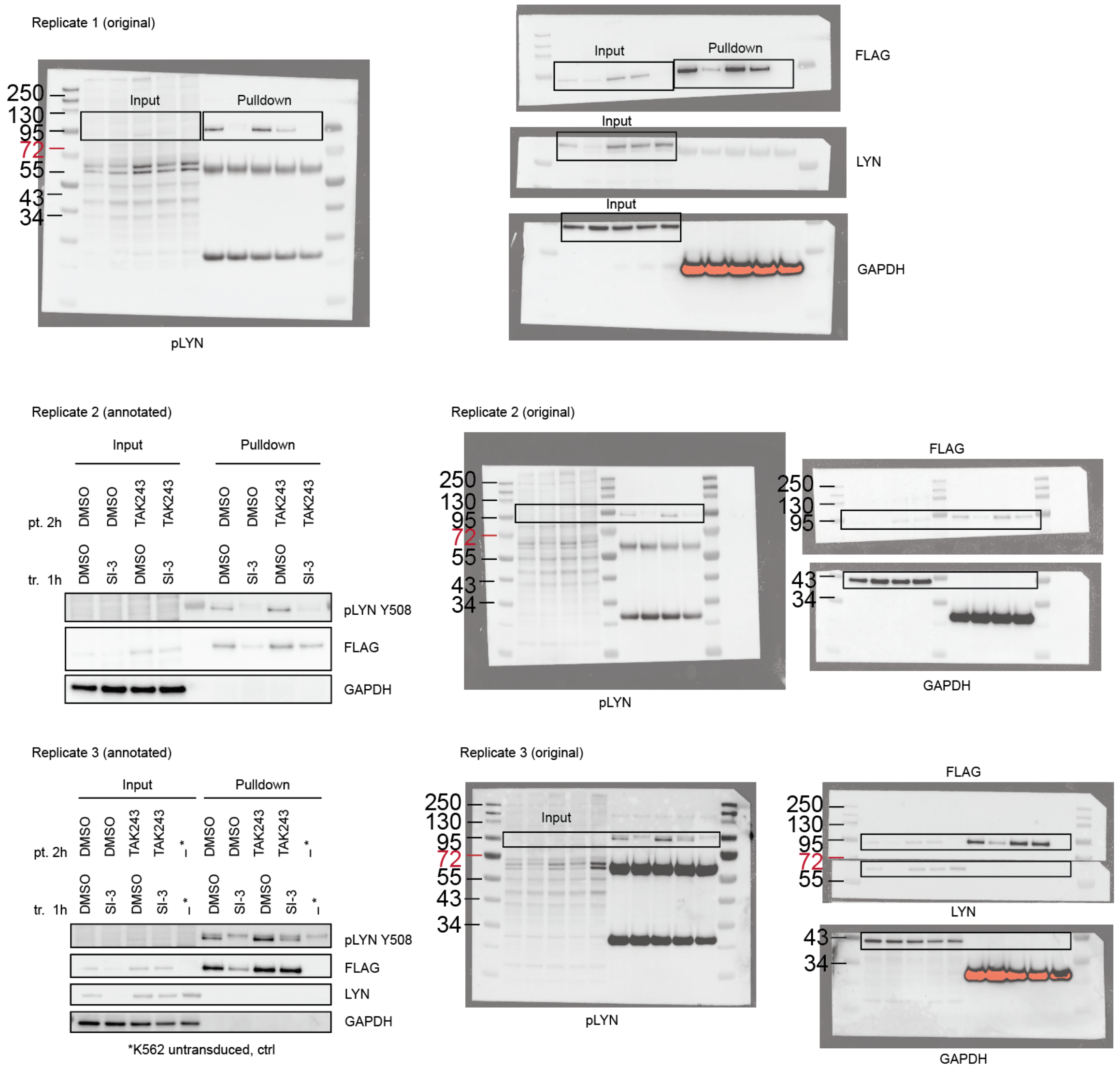


**Fig S17 Uncropped immunoblots and further replicates to Extended Data Fig 3L.**

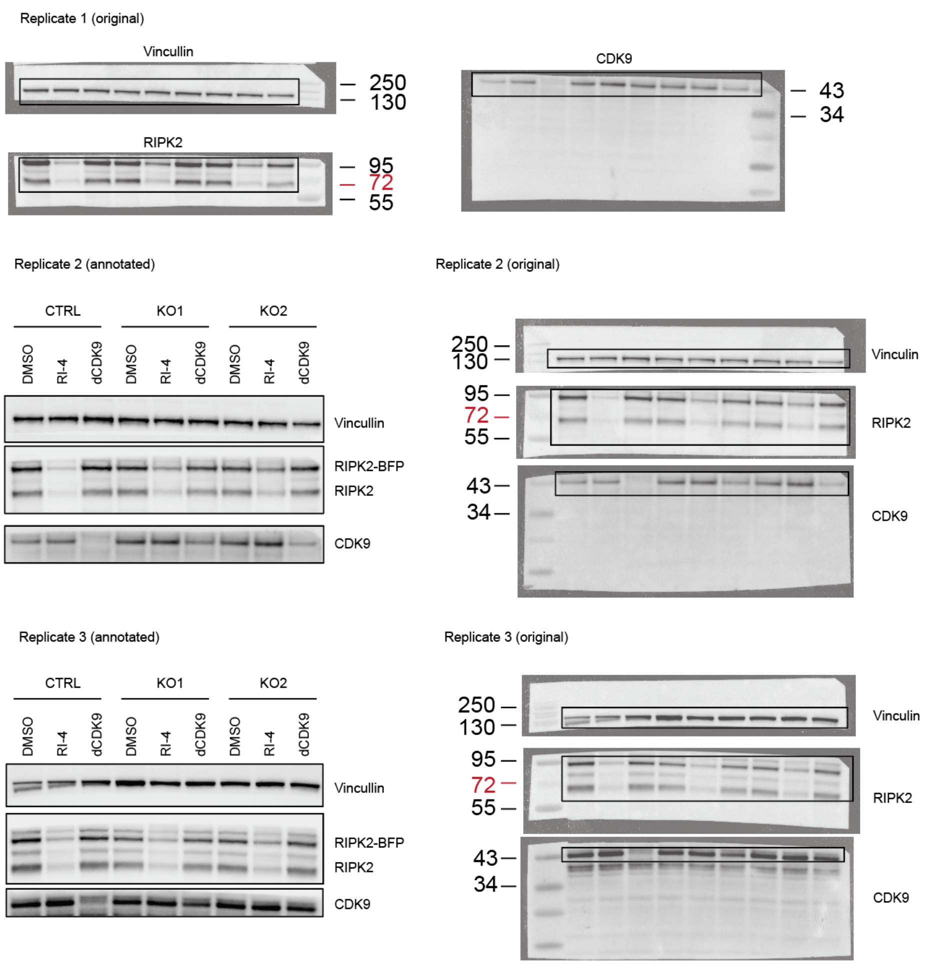


**Fig S18 Uncropped immunoblots and further replicates to Extended Data Fig 4D.**
