## Supplementary Document 1 for "Inhibitor-induced supercharging of kinase turnover via endogenous proteolytic circuits"

**Temporal trajectories of all hits**

Kinases defined as hits are highlighted in color, all other kinases in grey, the black line indicates the mean of all kinases. Errorbars are included as CI only for hit kinases.

Mutant kinases are abbreviated by their specific point mutation.

x-axis = Time (h) with ticks indicating 2,10 and 18h.

y-axis = POC with ticks indicating 0, 50, 100 and 150

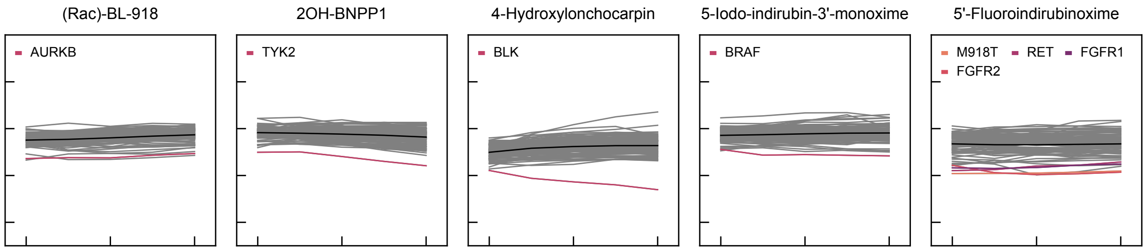


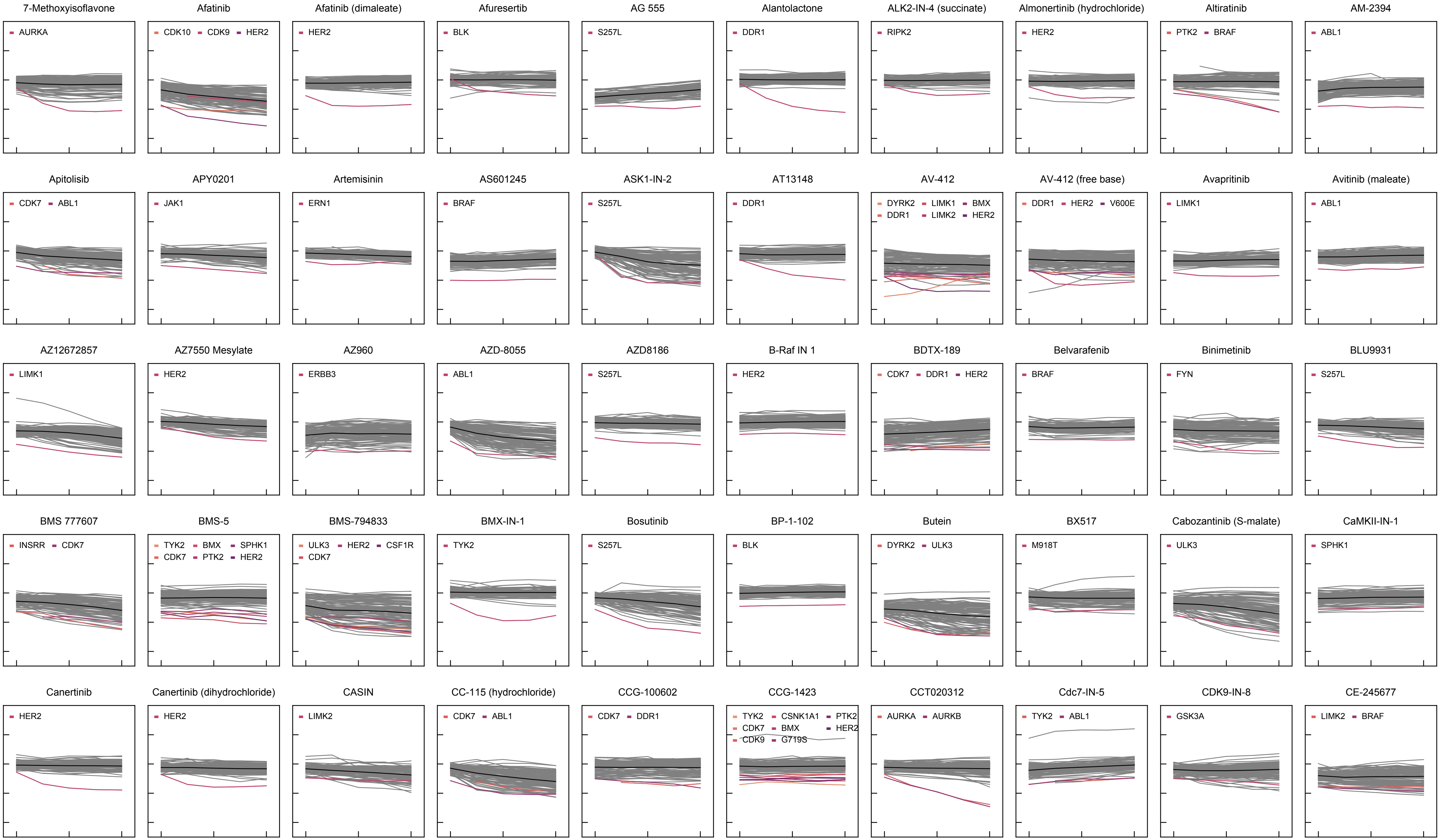

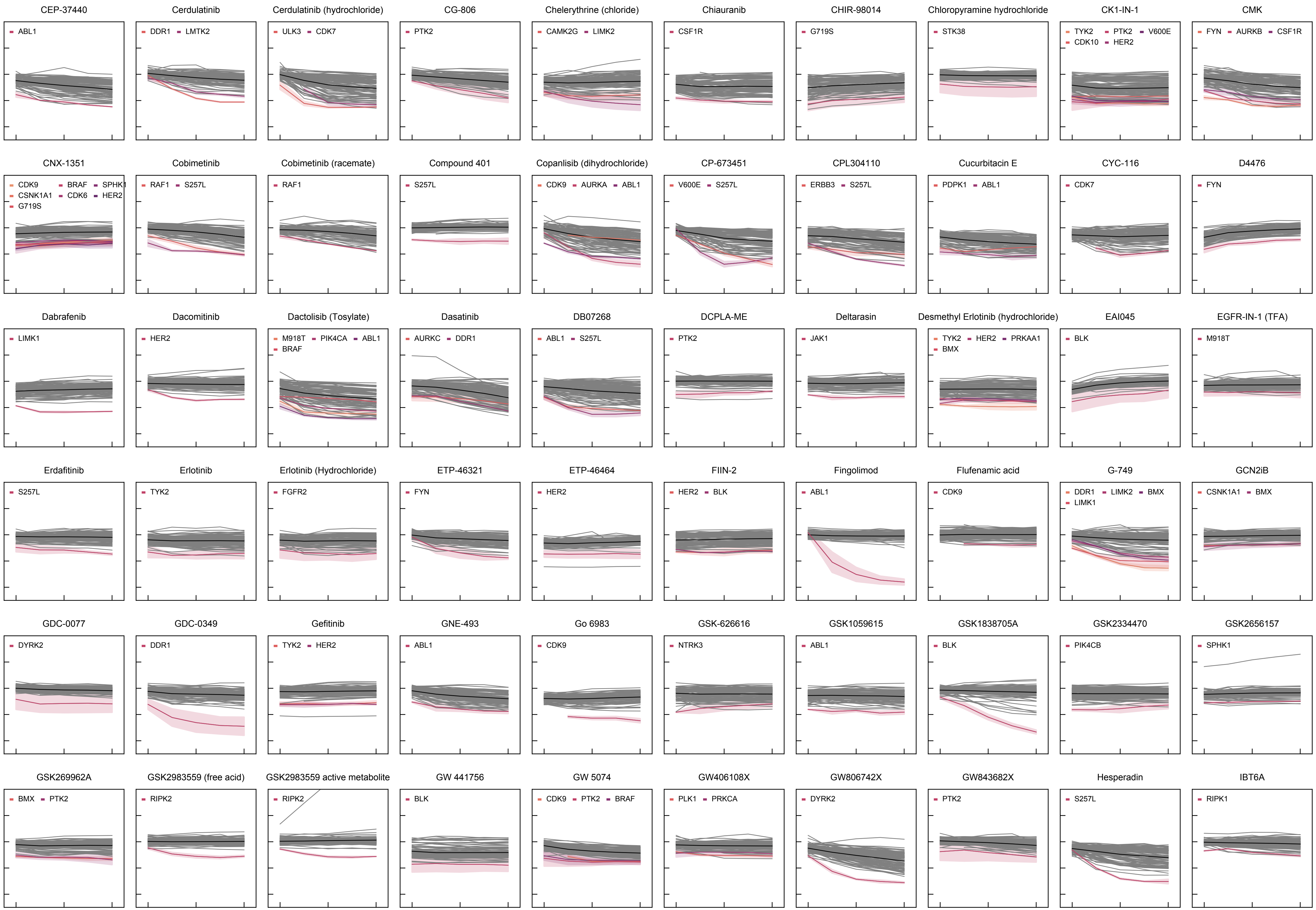

CEP-37440
ABL1
Cerdulatinib
DDR1
LMTK2
Cerdulatinib (hydrochloride)
ULK3
CDK7
CG-806
PTK2
Chelerythrine (chloride)
CAMK2G
LIMK2
Chiauranib
CSF1R
CHIR-98014
G719S
Chloropyramine hydrochloride
STK38
CK1-IN-1
TYK2
PTK2
V600E
CDK10
HER2
CMK
FYN
AURKB
CSF1R
CNX-1351
CDK9
BRAF
SPHK1
CSNK1A1
CDK6
HER2
G719S
Cobimetinib
RAF1
S257L
Cobimetinib (racemate)
RAF1
Compound 401
S257L
Copanlisib (dihydrochloride)
CDK9
AURKA
ABL1
CP-673451
V600E
S257L
CPL304110
ERBB3
S257L
Cucurbitacin E
PDPK1
ABL1
CYC-116
CDK7
D4476
FYN
Dabrafenib
LIMK1
Dacomitinib
HER2
Dactolisib (Tosylate)
M918T
PIK4CA
ABL1
BRAF
Dasatinib
AURKC
DDR1
DB07268
ABL1
S257L
DCPLA-ME
PTK2
Deltarasin
JAK1
Desmethyl Erlotinib (hydrochloride)
TYK2
HER2
PRKAA1
BMX
EAI045
BLK
EGFR-IN-1 (TFA)
M918T
Erdafitinib
S257L
Erlotinib
TYK2
Erlotinib (Hydrochloride)
FGFR2
ETP-46321
FYN
ETP-46464
HER2
FIIN-2
HER2
BLK
Fingolimod
ABL1
Flufenamic acid
CDK9
G-749
DDR1
LIMK2
BMX
LIMK1
GCN2iB
CSNK1A1
BMX
GDC-0077
DYRK2
GDC-0349
DDR1
Gefitinib
TYK2
HER2
GNE-493
ABL1
Go 6983
CDK9
GSK-626616
NTRK3
GSK1059615
ABL1
GSK1838705A
BLK
GSK2334470
PIK4CB
GSK2656157
SPHK1
GSK269962A
BMX
PTK2
GSK2983559 (free acid)
RIPK2
GSK2983559 active metabolite
RIPK2
GW 441756
BLK
GW 5074
CDK9
PTK2
BRAF
GW406108X
PLK1
PRKCA
GW806742X
DYRK2
GW843682X
PTK2
Hesperadin
S257L
IBT6A
RIPK1

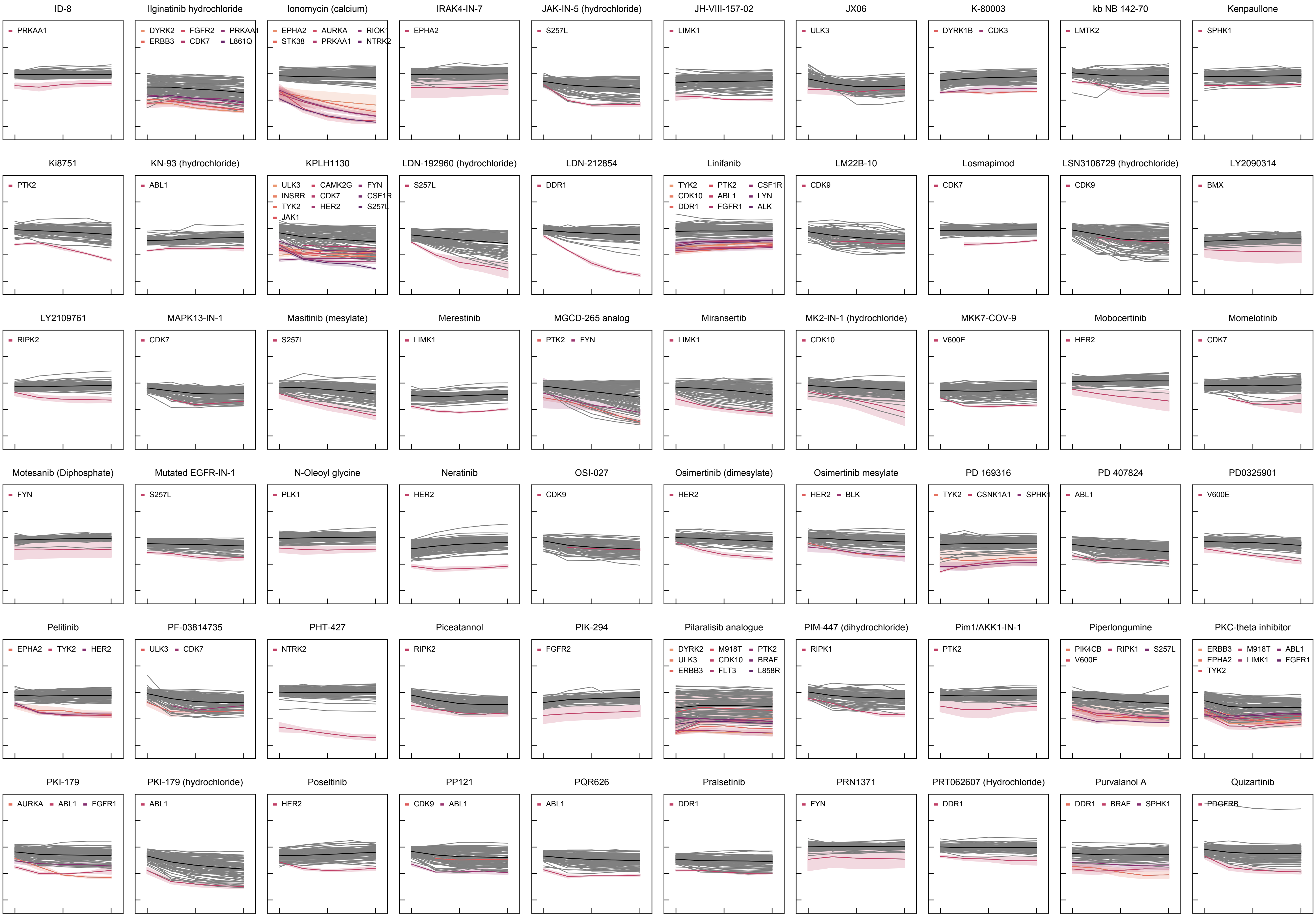
ID-8
PRKAA1
Ilginatinib hydrochloride
DYRK2 FGFR2 PRKAA1 ERBB3 CDK7 L861Q
Ionomycin (calcium)
EPHA2 AURKA RIOK1 STK38 PRKAA1 NTRK2
IRAK4-IN-7
EPHA2
JAK-IN-5 (hydrochloride)
S257L
JH-VIII-157-02
LIMK1
JX06
ULK3
K-80003
DYRK1B CDK3
kb NB 142-70
LMTK2
Kenpaullone
SPHK1
Ki8751
PTK2
KN-93 (hydrochloride)
ABL1
KPLH1130
ULK3 CAMK2G FYN INSRR CDK7 CSF1R TYK2 HER2 S257L JAK1
LDN-192960 (hydrochloride)
S257L
LDN-212854
DDR1
Linifanib
TYK2 PTK2 CSF1R CDK10 ABL1 LYN DDR1 FGFR1 ALK
LM22B-10
CDK9
Losmapimod
CDK7
LSN3106729 (hydrochloride)
CDK9
LY2090314
BMX
LY2109761
RIPK2
MAPK13-IN-1
CDK7
Masitinib (mesylate)
S257L
Merestinib
LIMK1
MGCD-265 analog
PTK2 FYN
Miransertib
LIMK1
MK2-IN-1 (hydrochloride)
CDK10
MKK7-COV-9
V600E
Mobocertinib
HER2
Momelotinib
CDK7
Motesanib (Diphosphate)
FYN
Mutated EGFR-IN-1
S257L
N-Oleoyl glycine
PLK1
Neratinib
HER2
OSI-027
CDK9
Osimertinib (dimesylate)
HER2
Osimertinib mesylate
HER2 BLK
PD 169316
TYK2 CSNK1A1 SPHK1
PD 407824
ABL1
PD0325901
V600E
Pelitinib
EPHA2 TYK2 HER2
PF-03814735
ULK3 CDK7
PHT-427
NTRK2
Piceatannol
RIPK2
PIK-294
FGFR2
Pilaralisib analogue
DYRK2 M918T PTK2 ULK3 CDK10 BRAF ERBB3 FLT3 L858R
PIM-447 (dihydrochloride)
RIPK1
Pim1/AKK1-IN-1
PTK2
Piperlongumine
PIK4CB RIPK1 S257L V600E
PKC-theta inhibitor
ERBB3 M918T ABL1 EPHA2 LIMK1 FGFR1 TYK2
PKI-179
AURKA ABL1 FGFR1
PKI-179 (hydrochloride)
ABL1
Poseltinib
HER2
PP121
CDK9 ABL1
PQR626
ABL1
Pralsetinib
DDR1
PRN1371
FYN
PRT062607 (Hydrochloride)
DDR1
Purvalanol A
DDR1 BRAF SPHK1
Quizartinib
PDGFRB

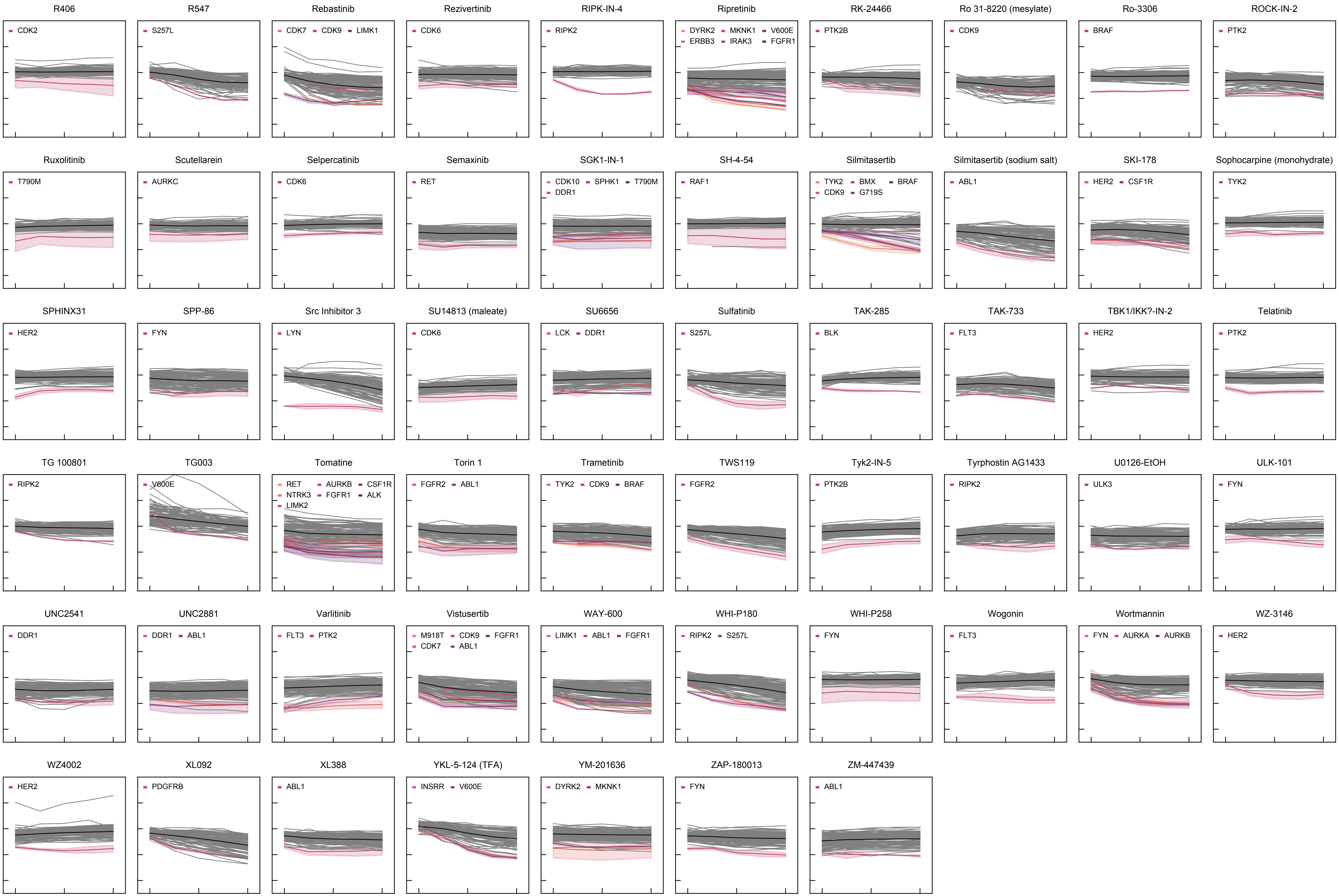
R406
CDK2
R547
S257L
Rebastinib
CDK7
CDK9
LIMK1
Rezivertinib
CDK6
RIPK-IN-4
RIPK2
Ripretinib
DYRK2
MKNK1
V600E
ERBB3
IRAK3
FGFR1
RK-24466
PTK2B
Ro 31-8220 (mesylate)
CDK9
Ro-3306
BRAF
ROCK-IN-2
PTK2
Ruxolitinib
T790M
Scutellarein
AURKC
Selpercatinib
CDK6
Semaxinib
RET
SGK1-IN-1
CDK10
SPHK1
T790M
DDR1
SH-4-54
RAF1
Silmitasertib
TYK2
BMX
BRAF
CDK9
G719S
Silmitasertib (sodium salt)
ABL1
SKI-178
HER2
CSF1R
Sophocarpine (monohydrate)
TYK2
SPHINX31
HER2
SPP-86
FYN
Src Inhibitor 3
LYN
SU14813 (maleate)
CDK6
SU6656
LCK
DDR1
Sulfatinib
S257L
TAK-285
BLK
TAK-733
FLT3
TBK1/IKK?-IN-2
HER2
Telatinib
PTK2
TG 100801
RIPK2
TG003
V600E
Tomatine
RET
AURKB
CSF1R
NTRK3
FGFR1
ALK
LIMK2
Torin 1
FGFR2
ABL1
Trametinib
TYK2
CDK9
BRAF
TWS119
FGFR2
Tyk2-IN-5
PTK2B
Tyrphostin AG1433
RIPK2
U0126-EtOH
ULK3
ULK-101
FYN
UNC2541
DDR1
UNC2881
DDR1
ABL1
Varlitinib
FLT3
PTK2
Vistusertib
M918T
CDK9
FGFR1
CDK7
ABL1
WAY-600
LIMK1
ABL1
FGFR1
WHI-P180
RIPK2
S257L
WHI-P258
FYN
Wogonin
FLT3
Wortmannin
FYN
AURKA
AURKB
WZ-3146
HER2
WZ4002
HER2
XL092
PDGFRB
XL388
ABL1
YKL-5-124 (TFA)
INSRR
V600E
YM-201636
DYRK2
MKNK1
ZAP-180013
FYN
ZM-447439
ABL1
